# Sub-communities of the vaginal microbiota in pregnant and non-pregnant women

**DOI:** 10.1101/2021.12.10.471327

**Authors:** Laura Symul, Pratheepa Jeganathan, Elizabeth K. Costello, Michael France, Seth M. Bloom, Douglas S. Kwon, Jacques Ravel, David A. Relman, Susan Holmes

**Affiliations:** Department of Statistics, Stanford University, 390 Jane Stanford Way, Stanford, CA 94305, USA; Department of Mathematics and Statistics, McMaster University, 1280 Main Street, West Hamilton, Ontario L8S 4K1, Canada; Department of Medicine, Stanford University School of Medicine, 300 Pasteur Drive, Stanford, CA 94305 USA; Institute for Genome Sciences, University of Maryland School of Medicine, 670 W. Baltimore Street, Baltimore, MD 21201, USA; Department of Microbiology and Immunology, University of Maryland School of Medicine, 685 West Baltimore Street, HSF-I Suite 380, Baltimore, MD 21201, USA; Division of Infectious Diseases, Massachusetts General Hospital, 55 Fruit Street, Boston MA 02114, USA; Harvard Medical School, 25 Shattuck St, Boston, MA 02115, USA; Ragon Institute of MGH, MIT, and Harvard, 400 Technology Square, Cambridge MA 02139, USA; Department of Microbiology & Immunology, Stanford University School of Medicine, 299 Campus Drive, Stanford, CA 94305, USA; Infectious Diseases Section, Veterans Affairs Palo Alto Health Care System, 3801 Miranda Avenue, Palo Alto, CA 94304, Palo Alto, CA 94304, USA; The Vaginal Microbiome Research Consortium (VMRC)

**Keywords:** Vaginal microbiota, multi-omics, menstrual cycle, pregnancy

## Abstract

Diverse and non-*Lactobacillus*-dominated vaginal microbial communities are associated with adverse health outcomes such as preterm birth and the acquisition of sexually transmitted infections. Despite the importance of recognizing and understanding the key risk-associated features of these communities, their heterogeneous structure and properties remain ill-defined. Clustering approaches are commonly used to characterize vaginal communities, but they lack sensitivity and robustness in resolving substructures and revealing transitions between potential sub-communities. Here, we address this need with an approach based on mixed membership topic models, using longitudinal data from cohorts of pregnant and non-pregnant study participants. We identify several non-*Lactobacillus*-dominated sub-communities common to both cohorts and independent of reproductive status. In non-pregnant individuals, we find that the menstrual cycle modulates transitions between and within sub-communities. In addition, a specific non-*Lactobacillus*-dominated sub-community, which was associated with preterm delivery in pregnant participants, was also more common during menses, a time of elevated vaginal inflammation in non-pregnant participants. Overall, our analyses based on mixed membership models reveal substructures of vaginal ecosystems which may have important clinical and biological associations.

## Introduction

Several critical aspects of women’ s health are linked to the structure of the vaginal microbiota (1–3). Vaginal microbiotas dominated by beneficial *Lactobacillus* species are associated with positive health outcomes (3). A paucity of *Lactobacillus* and a diverse array of strict and facultative anaerobes, however, are associated with negative health outcomes such as preterm birth (4, 5) and susceptibility to sexually transmitted infections (6–9), including HIV (10–12). Longitudinal studies of vaginal microbiota composition have revealed its dynamic nature: microbiota composition frequently changes over time (4, 13, 14). In non-pregnant individuals, a virtually complete replacement of the microbiota is sometimes observed, typically around the time of menses (13, 15). While complete replacement is rare, more modest (*i*.*e*., of a fraction of the microbiota composition), or slower (*i*.*e*., over a few days or weeks) changes in composition are relatively common in both pregnant and non-pregnant individuals (4, 13, 14). The microbiota of pregnant women may appear more stable than that of non-pregnant individuals; however, differences in sampling frequencies (*e*.*g*., weekly during pregnancy *vs* daily outside of pregnancy) might not allow us to fully characterize the differences in microbiota dynamic. Non-*Lactobacillus* dominated microbiotas are generally less stable than *Lactobacillus* dominated ones (4, 13, 14). Some *Lactobacillus* species, such as *L. crispatus*, better resist invasion or replacement by non-*Lactobacillus* species and create greater vaginal ecosystem stability during and outside pregnancy (14, 16, 17). Other *Lactobacillus* species, such as *L. iners*, are more frequently associated with non-optimal communities (14, 16, 17). Non-optimal vaginal microbiotas (*i*.*e*., non-*Lactobacillus*-dominated microbiota) are typically highly heterogeneous within and between individuals (4, 14, 16). It remains, however, poorly understood whether non-optimal microbiota composition is random (*i*.*e*., individual-specific) or if distinct sub-communities (*i*.*e*., consortia of bacteria interacting with each other) exist within these diverse microbiotas. If such sub-communities do exist, it remains to be seen whether they are differentially associated with characteristics of the host or with specific negative health outcomes, such as preterm birth.

Efforts to address this question have so far relied on clustering approaches. Various clustering methods are commonly applied to taxonomic abundance tables to define community structure. This has led to the adoption of the concepts of community state types (CST) or community types (CTs) (18, 19). More recently, in order to define reference sub-CSTs (*i*.*e*., dataset- or study-independent state types), large composite datasets have been clustered, and several non-*Lactobacillus*-dominated clusters (sub-CSTs) have been identified across populations of non-pregnant women (20). Clustering serves as a useful dimensionality reduction tool for describing complex microbiota compositions. However, clustering-based categorization of samples may fail to capture clinically-relevant structures. For example, the vaginal microbiota of two women could belong to the same cluster because their microbiotas both show a bare majority of *L. iners* (*e*.*g*., 60%), but be accompanied by *L. crispatus* in one case, and by a diverse panel of non-*Lactobacillus* species in the other case. The two situations may appear similar (*i*.*e*., each may be assigned to CST III), but they may be driven by different mechanisms and have different health implications. In addition, clustering based approaches fail to model *transition* or *intermediary states* between clusters (Fig 1). Modeling *transitions* is especially important in the context of the vaginal microbiota as its composition may change several times over a few months, weeks or even a few days, as observed in non-pregnant, menstruating individuals (4, 14–16). However, because samples are assigned only to a single cluster (Fig1a), transitions between clusters may appear identical (*i*.*e*., described by the same sequence of clusters) while the underlying microbiota trajectories were drastically different in rate (progressive vs abrupt) or in the nature of the intermediate compositions. Finally, while clustering approaches can identify sets of species that frequently co-occur, they are not well suited to identify subsets of species that may have similar functions but that are not frequently found together (Fig 1b). These discrepancies between the clustering assumptions and our understanding of the composition and dynamics of the vaginal microbiota highlight the need for better-suited dimension reduction statistical models.

**Figure 1:**
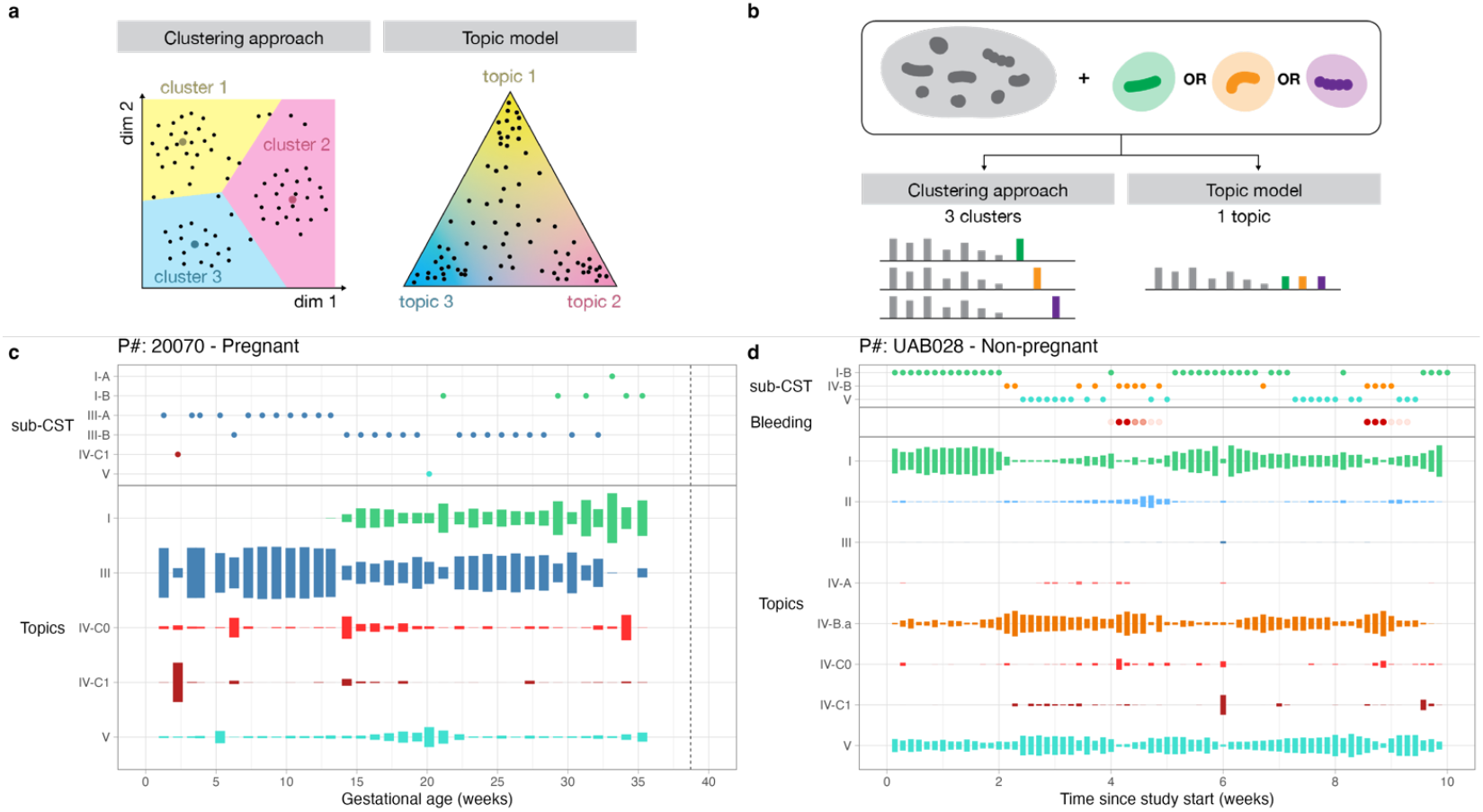
Topic models are mixed membership models that reveal transitions between states. **(a)** Schematics contrasting sample characterization in a lower dimensional space by clustering methods versus topic models. In both schematics, each dot is a sample. Larger colored dots in the clustering schematic indicate centroids. **(b)** Schematic illustrating the phenomenon of “functional equivalence” and how clustering methods versus topic models represent it. We consider two or more species potentially “functionally equivalent” if they tend to occupy the same ecological niche (thrive in similar environments and with other species) but are rarely found together because they may compete for the same resources. **(c-d)** Examples of time-series displays of changes in microbiota composition summarized by clusters membership (sub-CST - top) or topic proportions (bottom) in a pregnant (panel c) and non-pregnant (panel d) participant. Topics were labeled such that their name matched the (sub)CST with the most similar composition (see Fig. 2c).

Topic models, first developed to infer population structure (21) and later formally described as Latent Dirichlet Allocation (LDA) in the context of natural language processing (22), have recently been proposed for analyzing microbial communities and identifying sub-communities (23). In contrast to clustering-based categorization, where each sample is assigned to a single category based on the closest cluster, samples are modeled as mixtures of topics (sub-communities), and each topic is characterized by a particular distribution of bacterial species or strains. For example, if a sample were described as 70% topic 1 and 30% topic 2, this would mean that the species subsumed in topic 1 accounted for 70% of the sample, while the species in topic 2 accounted for the remaining 30%. Some species can be found in several topics (*e*.*g*., a species can co-exist within two distinct sub-communities). Topics may be composed of a few species or strains (sparse topics) or include a larger number. In addition to providing a more realistic model of microbiota composition, topic models present the advantage of not requiring any normalization of the taxa count tables (typically the number of 16S rRNA genes sequenced in each sample) as they are hierarchical Bayesian models explicitly accounting for library sizes.

In this study, we sought to deepen our understanding of the fine structure of non-optimal vaginal microbiotas by applying topic models (mixed membership models) to longitudinal samples acquired from pregnant and non-pregnant women. We examined the similarities and differences in sub-community composition between cohorts and compared them to previously identified reference clusters. We then investigated the clinical relevance of the identified sub-communities and their association with host characteristics, pregnancy status, phase of the menstrual cycle (in non-pregnant individuals), and the risk of preterm birth (in pregnant individuals). The concentrations of vaginal metabolites (both host- and bacteria-produced) and cytokines (host-produced) were also quantified longitudinally in non-pregnant individuals but at a lower temporal resolution (five samples from 40 non-pregnant participants) and were analyzed for correlations with the menstrual cycle.

## Results

### Topic analysis identifies nine sub-communities in the vaginal microbiota of pregnant and non-pregnant women

We analyzed data from 2,179 vaginal samples collected weekly from 135 pregnant individuals enrolled at two sites in the United States (Stanford University, Stanford, CA and University of Alabama, Birmingham, AL) and 1,534 vaginal samples collected daily from 30 non-pregnant individuals enrolled at the University of Alabama, Birmingham (Methods, Table S1 for demographic data). Topic models were fit to the count data of 16S rRNA amplicon sequence variants (ASVs) agglomerated by taxonomic assignment.

Topic analysis requires choosing K, the number of topics, to model the provided count data. K can be estimated using cross-validation or, as recently proposed (24), by performing topic alignment across models with different resolutions (*i*.*e*., with different K, Fig 2a). In contrast to cross-validation, this latter approach shows how topics at higher resolution relate to topics at lower resolution and provides several diagnostic scores. These scores characterize each topic across degrees of resolution and allow us to evaluate whether the data deviate from LDA assumptions. Our topic alignment suggested that 9 topics provided the best compromise between dimension reduction and accurate modeling of taxonomic counts (Methods, SI, Fig 2a-b). If a coarser resolution were desired, the alignment refinement scores suggested that K = 5 topics would be the most suited as topics at higher resolutions were sub-topics of these five topics (SI, Fig 2b).

**Figure 2:**
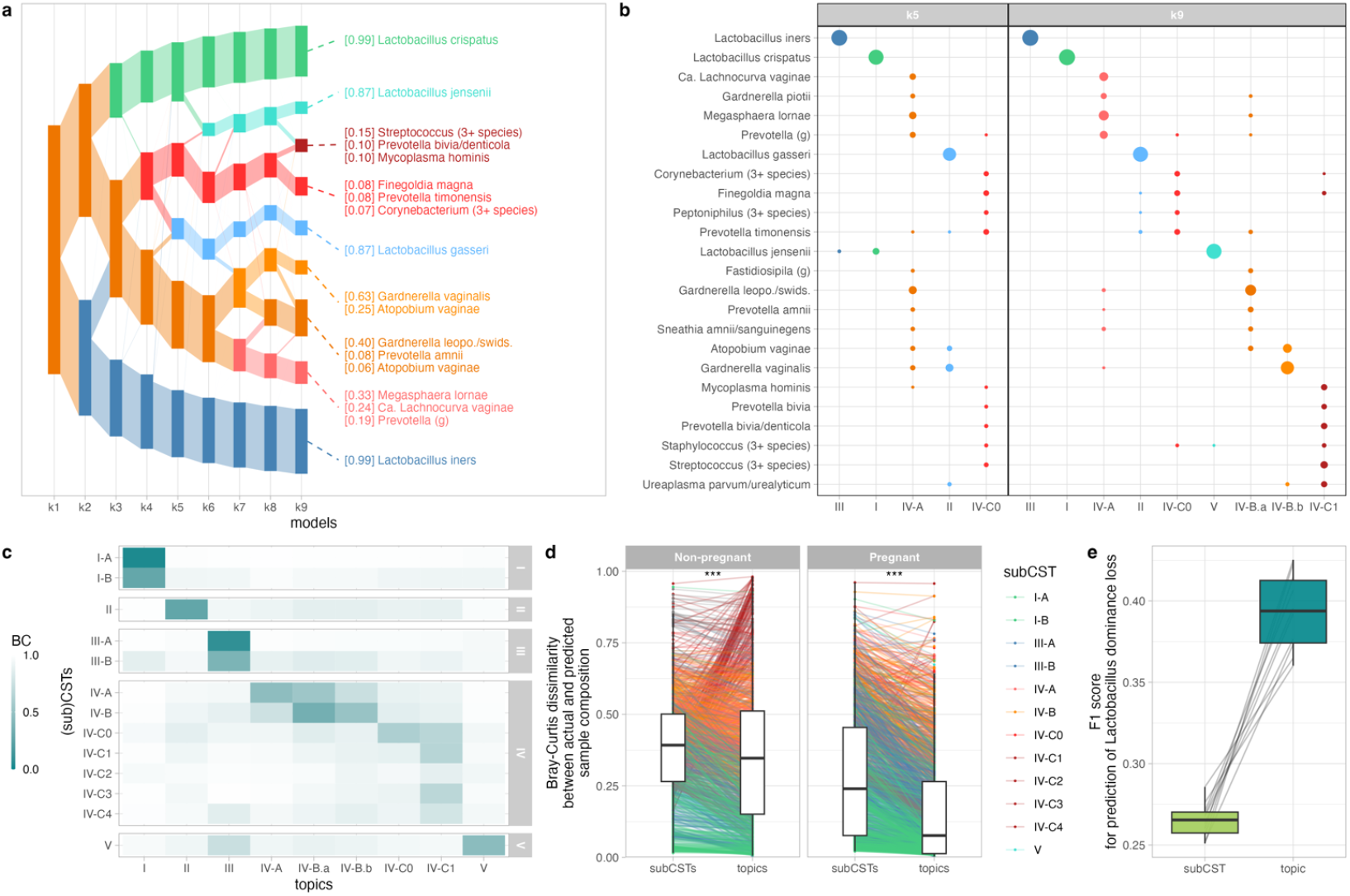
Sub-communities identified by topic models. **(a)** Alignment of topics (rectangles) for models fitted with an increasing number of topics (x-axis). The height of the rectangles is scaled according to the total proportion of the corresponding topic in all samples: taller rectangles represent more prevalent topics. Topics are connected across models (x-axis) according to their alignment weight, which reflect their similarities (Methods). Topics of the k = 9 model are annotated with their most prevalent species, and the numbers in brackets in front of each species indicate the proportion of that species in the topic. The annotations included the three most prevalent species that made up at least 5% of the topic composition. **(b)** Topic composition for k = 5 (coarse representation) or k = 9 (optimal tradeoff between dimension reduction and descriptive accuracy) topics (side-by-side panels). The proportion of each species (y-axis) within each topic (x-axis) is encoded by the size of the dots. These proportions sum to 1 for each topic. For readability and conciseness of the figure, species were included if they accounted for at least 0.5% of a topic composition. **(c)** Comparison of the topic (x-axis) and sub-CST (y-axis) compositions. Compositions were compared using the Bray-Curtis dissimilarity. Topics and sub-CSTs with similar compositions are characterized by a low divergence and a darker hue. **(d)** Bray-Curtis dissimilarity between actual sample composition and predicted sample composition (y-axis) by sub-CSTs or topics (x-axis) in non-pregnant (left panel) and pregnant (right panel) individuals. Each line is a sample, colored by its sub-CST membership. Stars in each panel indicate statistical significance of a one-sided paired t-test (***: < 0.001) **(e)** F1 scores (y-axis) for the prediction of Lactobacillus dominance loss (*i*.*e*., total proportion of Lactobacillus falling below 50%) at the next sample when the loss is predicted from sub-CST membership (light green) or topic memberships (dark turquoise). The F1 score is the harmonic mean of the precision and the sensitivity of the predictions. Distributions were obtained from 10 independent training-testing sets (Methods, SI). Thin lines connect F1 scores from the same training-testing set.

At K = 9, four of these nine topics were dominated by one of the four most common *Lactobacillus* spp. (*L. crispatus, L. gasseri, L. iners*, and *L. jensenii*, Fig 2a-b). The composition of the five remaining topics did not include any *Lactobacillus* spp. (Fig 2a-b). These five non-Lactobacillus topics could be grouped into two groups based on the topic alignment: one group contained three topics which included *Gardnerella, Atopobium*, and *Megaspaera* spp., while the other group contained *Finegoldia, Corynebacterium*, and *Streptococcus* (Fig 2a-b).

### Topics provide a more succinct, yet more accurate, description of microbiota composition than sub-CSTs

To evaluate the generalizability of the identified sub-communities, we compared the topic composition with the composition of the 12 “reference” clusters (sub-CSTs, Valencia centroids) described previously and identified in a composite dataset of non-pregnant individuals’ samples (20) (Fig 2c). To compare topic and cluster compositions, we computed the Bray-Curtis dissimilarities between the two compositions after harmonizing taxonomic assignments (Fig 2c, Methods, SI). Topics were labeled to match the (sub-)CST label of the cluster to which they were most similar (Methods) (Fig. 1c-d, Fig. 2b). The comparison showed that two *L. crispatus*-dominated sub-CSTs (I-A and I-B) have high similarity with the single *L. crispatus-dominated* topic (I). Similarly, two *L. iners*-dominated sub-CSTs (III-A and III-B) match a single *L. iners*-dominated topic (III). This is because CST I-A and I-B (or III-A and III-B) describe microbiotas that are either fully dominated by *L. crispatus* (subCST I-A) or *L. iners* (subCST III-A) versus those dominated by *L. crispatus* or *L. iners* but also hosting other species (sub-CST I-B or III-B). In contrast, because topic models allow samples to be composed of several topics, a single topic is sufficient to account for *L. crispatus* (topic I) or *L. iners* (topic III) counts. Samples in which *L. crispatus* co-exists with *L. iners* will be represented by a mix of topics I and III, while a sample where *L. crispatus* co-exists with a *Gardnerella* species by a mix of topics I and IV-A/B. CST II and V have a one-to-one optimal match with topics II and V.

When comparing the non-*Lactobacillus* sub-CSTs and topics, we observed that (i) sub-CST IV-A and IV-B are represented by three topics (IV-A, IV-B.a, and IV-B.b), which can, in part, be explained by differences in taxonomic assignment used for topics (*e*.*g*., *Gardnerella* species are undifferentiated in sub-CSTs, while, here, some *Gardnerella* ASVs were matched to different species), and (ii) a single topic (IV-C1) matches four sub-CSTs (IV-C1 – IV-C4). This is because these four sub-CSTs only differ in the proportion of 4 seemingly mutually exclusive species (*Streptococcus, Enterococcus, Bifidobacterium*, and *Staphylococcus*), with one of these four species dominating each sub-CST; the prevalence of the remaining species is similar across the four IV-C1-4 sub-CSTs. In contrast, because topic models allow for synonyms, topic IV-C1 embeds these species within a single topic, as illustrated in Fig 1b.

We next examined three potential benefits of using topic mixed-memberships instead of clustering categorization (sub-CSTs). Our first conjecture was that topics would provide a more accurate representation of sample compositions than sub-CSTs. The second was that this effect would be primarily driven by samples from unstable microbiotas. Our third conjecture held that topic membership would better predict whether an individual is at risk of losing Lactobacillus dominance at the next time-point.

To test our first conjecture (*i*.*e*., accuracy of representation), we compared the Bray-Curtis dissimilarity between the actual sample compositions and the sample compositions predicted by topic mixed memberships or by sub-CST membership. The predicted composition of a sample is either the composition of the centroid of the sample’ s sub-CST or the average topic composition (displayed in figure 2b) weighted by the proportion of each topic in that sample (Methods). The Bray-Curtis dissimilarity between actual sample composition and predicted sample composition was smaller when sample compositions were predicted by topics (Fig 2d). This effect was stronger in pregnant participants (mean difference = 0.12, paired t-test p-value < 0.001) than in non-pregnant participants (mean difference = 0.02, p-value < 0.001). The smaller mean difference in non-pregnant women compared to pregnant women can partially be explained by samples belonging to sub-CSTs IV-C1-4. These samples were dominated by one of the four seemingly mutually exclusive species mentioned above (*Streptococcus, Enterococcus, Bifidobacterium*, and *Staphylococcus*), considered synonyms in topic models, and found in a single topic. When samples from sub-CST IV-C1-4 were omitted, the mean difference in dissimilarity in non-pregnant women increased from 0.02 to 0.07.

Our second conjecture was that the composition of samples from stable microbiotas (*i*.*e*., the microbiota composition remains largely unchanged over time) would be equally well described by sub-CSTs or by topics because these microbiotas would have stabilized over more robust sub-communities that can be well captured by clustering approaches. In contrast, we expected that samples from unstable microbiotas would be better described by topic mixed memberships because the transitions between well-defined sub-communities can be captured better by varying memberships. Our results supported this expectation in pregnant participants, but not in non-pregnant participants (Fig S1). To test this expectation, we used the Bray-Curtis dissimilarities computed above and compared their differences (sub-CSTs *vs* topics) in samples from stable vs unstable microbiotas. Samples were considered to harbor stable microbiotas if they belonged to a group of at least 5 consecutive samples whose Bray-Curtis dissimilarity was less than 0.25 (similar results were obtained for 0.15 and 0.35 thresholds – see Table S2) and were considered to harbor unstable microbiotas or transition states otherwise. In pregnant participants, the mean difference in dissimilarities was 0.08 for samples from stable microbiotas and 0.14 for samples from unstable microbiotas (one-sided t-test p-value < 0.001). In non-pregnant participants, these differences were approximately the same in samples from both stable (0.03) and unstable (0.02) microbiotas.

We next evaluated our third conjecture which was that topic memberships better identify individuals at risk of losing *Lactobacillus* dominance, defined here as overall *Lactobacillus* proportions falling below 50%. Past studies have shown that individuals whose microbiota is categorized as CST III (*L. iners*-dominated) are more at risk of losing *Lactobacillus* dominance than those in other *Lactobacillus*-dominated CSTs (I, II, and V) (17, 25) but this risk has not been evaluated with a more refined definition of microbiota composition. To do so, we trained logistic regression models to predict whether an individual would lose their *Lactobacillus* dominance. Prediction performances were then evaluated on an independent test set and the procedure was repeated ten times using random splits of the data into training and test sets (Methods). Since only 11% of *Lactobacillus* dominated microbiotas switch to non-*Lactobacillus* dominated ones (*i*.*e*., we are predicting rare events), the F1 score, which is the harmonic mean of the prediction’ s precision and sensitivity, was used to compare prediction performances (Fig 2e). This comparison shows that topic memberships better predict the risk of losing *Lactobacillus* dominance than sub-CST memberships do (median F1 score of 0.4 *vs* 0.27, Wilcoxon test *p-* value < 0.002). Specifically, topic-based predictions are more precise (*i*.*e*., have a lower false positive rate) than sub-CST-based predictions (precision of 0.26 vs 0.16, *p-*value < 0.002, Fig S2).

Given these results and the three advantages conferred by topic-based description of microbiota composition, we next explored the demographic associations and functional relevance of the identified sub-communities.

### Topic composition varies with demographic characteristics and pregnancy status

The samples used in this study were collected from three cohorts: non-pregnant women recruited at the University of Alabama Birmingham between 2009 and 2010, pregnant women recruited at the same institution between 2013 and 2015, and pregnant women recruited at Stanford University also between 2013 and 2015. Participants’ race and recruitment site were significantly associated with differential proportions of several topics. The microbiotas of Black participants and participants recruited at UAB were more likely to contain topics III (*L. iners*-dominated), IV-A, and IV-B.a (both non-*Lactobacillus*-dominated) (fig3a-c). Topics III and IV-A were also more prevalent in pregnant participants, while topics IV-B.b and IV-C1 were less prevalent than in non-pregnant participants (fig3a-c).

### Topics IV-C0 and IV-C1 increase during menses; topic IV-C1 is also associated with preterm birth

The proportions of both topics IV-C0 and IV-C1 increased during menses (p-values smaller than 0.001 and 0.01 resp., fig 3c). In contrast, the proportion of topic I (*L. crispatus*-dominated, p-value < 0.01) decreased during menses. Consistent with previous findings (4), topic I (*L. crispatus*-dominated) was associated with term delivery, while topic IV-C1 had a strong but mildly significant (p = 0.051) association with preterm delivery.

**Figure 3:**
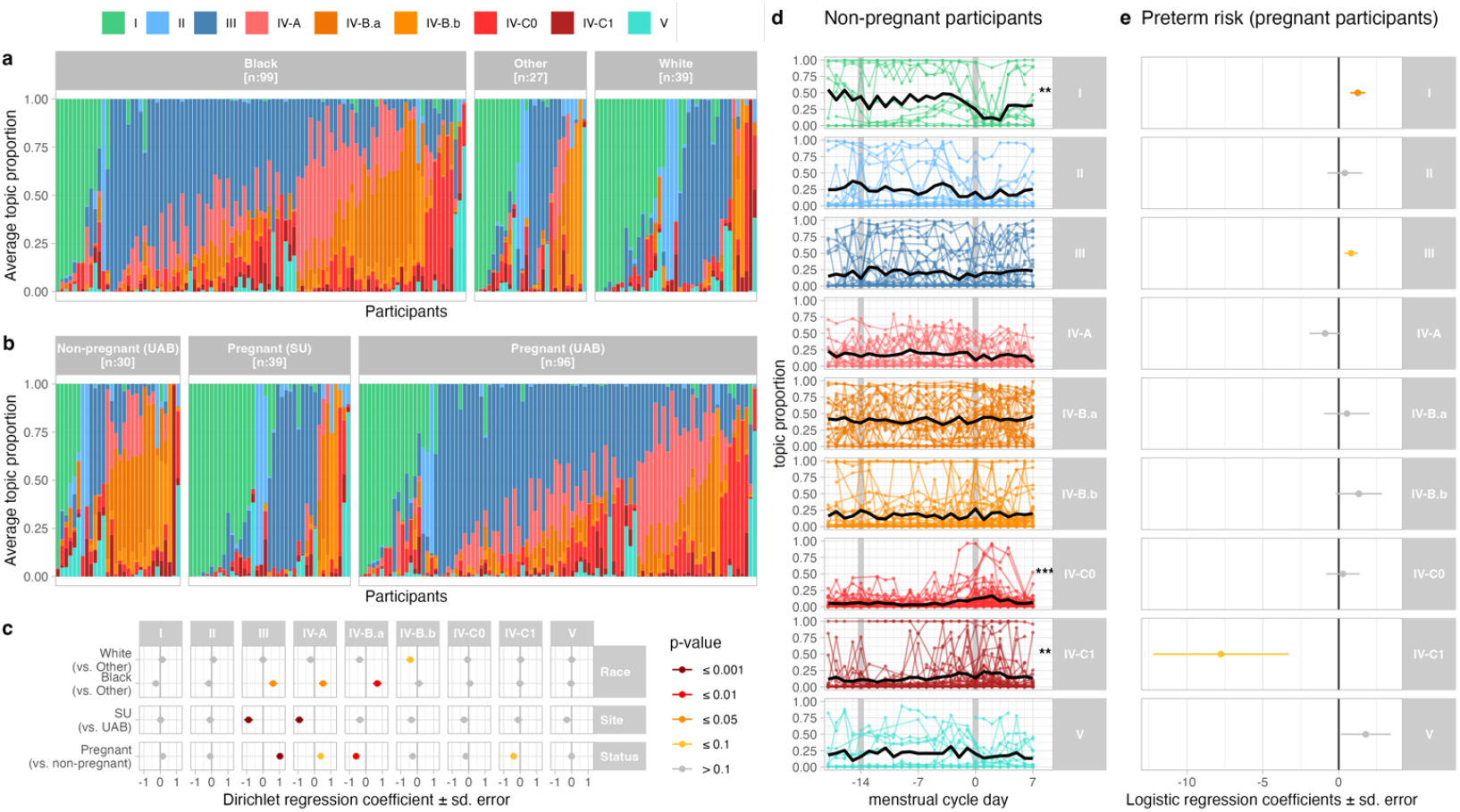
Sub-communities and demographic and reproductive characteristics. **(a-b)** Topic composition per racial group (a) or cohort (b). Vertical bars show the longitudinal average topic (color) proportion for each participant (x-axis). Participants are ordered by their most prevalent topic. **(c)** Dirichlet regression estimated coefficients (x-axis) quantifying the associations between race, study site, pregnancy status (y-axis) and topic proportions (horizontal panels). Colors indicate the strength of the statistical significance of the associations (p < 0.001: dark purple; p < 0.01: red; p < 0.05: orange; p < 0.1: yellow; p > 0.1: gray). **(d)** Topic proportions throughout the menstrual cycle (cycle day 0 indicates the first day of menses – see Fig 4a). Each dot is a sample. Lines connect samples from the same participant and cycle. Thick black lines show the average topic proportions across all participants. Stars on the right indicate the statistical significance of the associations between topic proportions and menstrual cycle (***: p < 0.001, **: p < 0.01). **(e)** Logistic regression estimated coefficients (x-axis) quantifying the association between average topic proportion and preterm birth in pregnant individuals. Colors are as in panel (c).

### The menstrual cycle shapes the vaginal microbial composition

Prompted by the observation that the proportions of several topics varied with the menstrual cycle, we further investigated longitudinal associations between menstrual cycle and microbiota composition. Among the 30 non-pregnant participants, 26 had reported vaginal bleeding patterns that allowed for identification of at least one menstrual cycle within the ten study weeks (Methods), and for 20 participants, we had data over two consecutive menstrual cycles. Cycles were standardized starting from 18 days before menses to 7 days after the first day of menses, given that the luteal phase (after ovulation) is known to vary less in duration than the follicular phase (before ovulation) (26, 27) (Fig 4a, Methods). Ovulation was assumed to occur around 2 weeks before the first day of menses based on average luteal phase duration (26, 27).

**Figure 4:**
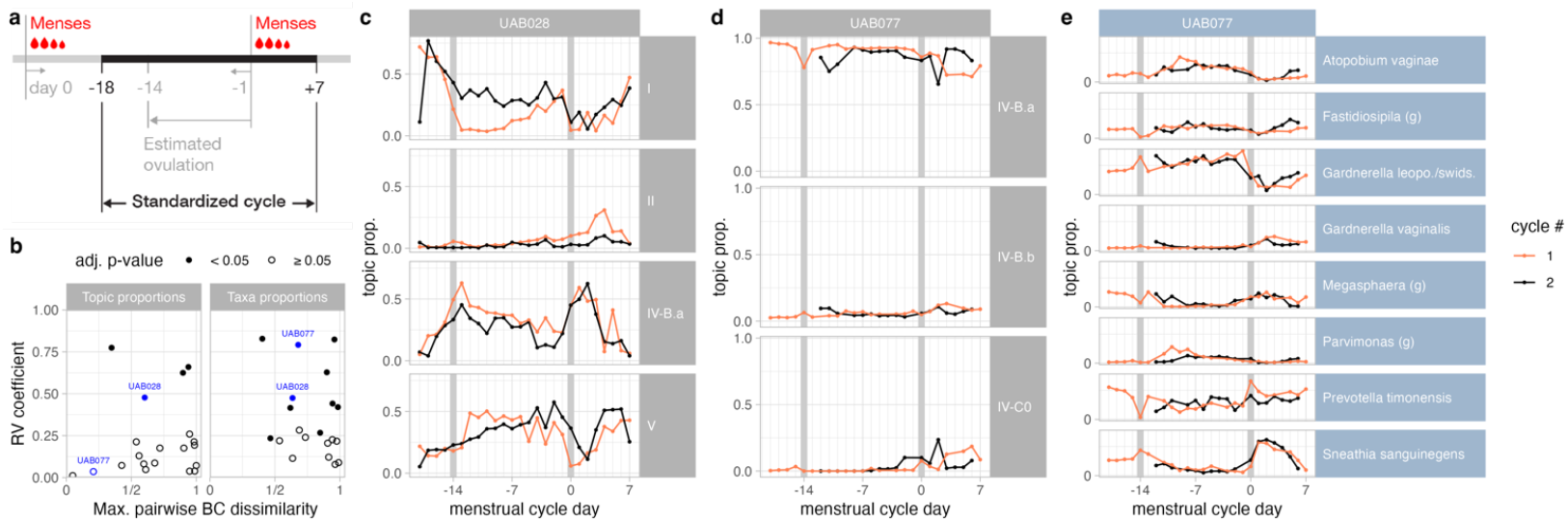
The menstrual cycle shapes the microbial composition. **(a)** Schematic illustrating the features of standardized cycles. **(b)** Scatter plot, in which each dot is a participant, showing the RV coefficient of agreement (y-axis) between the relative proportions of topics (left panel) or taxa (right panels) of a participant’ s consecutive cycles and the magnitude of change in microbiota composition throughout the cycle measured by the maximum of the pairwise Bray-Curtis dissimilarity between the average topic or taxa proportions for each cycleday (x-axis). Participants selected for panels c-e are highlighted in blue. **(c-d)** Topic composition of two participants with data available for at least two full menstrual cycles. The first menstrual cycle is displayed in orange and the second in black. These two participants were selected to show the diversity of temporal profiles. The time series display shows topic proportion (y-axis) on each cycle day (x-axis). For each study participant, topics were included if their median proportion across cycles was higher than 1% and their maximal proportion was higher than 5%. **(e)** Same display as in panels c-d but where the y-axis shows the relative abundance of each taxon for that participant. Taxa were included following the criteria used to select topics in panels c-d.

The vaginal microbiota structure of 4/20 participants (20%), characterized by topic proportions, showed a statistically significant agreement between consecutive cycles (Fig 4b-d) as measured by the RV coefficient (adj. p-value < 0.05, Methods). However, while the topic proportions may remain relatively stable throughout cycles, the underlying taxa composition may vary (*e*.*g*., for participant UAB077, fig 4d-e). Half (10/20) of the participants had a statistically significant agreement between their taxa proportions in two consecutive cycles (Fig 4b, right panel).

We further investigated whether the vaginal environment, characterized by pH values and vaginal metabolite and cytokine concentrations, varied with the menstrual cycle. Consistent with past results (18), the vaginal pH of *Lactobacillus-*dominated samples (*i*.*e*., proportions of Lactobacillus > 50%) was lower (4.4, 90% 4.0-5.3) than that of non-*Lactobacillus-*dominated samples (5.0, 90% 4.0-5.8). The pH remained stable throughout the cycle (*Lactobacillus*-dominated: 4.3, 90% 4.0-5.3; non-*Lactobacillus* dominated: 4.9, 90% 4.0-5.5), except during menses when it increased by about 0.5 units in *Lactobacillus*-dominated (4.7, 90% 4.0-5.8) and non-*Lactobacillus*-dominated samples (5.4, 90% 4.4-7.0) (Fig 5a).

**Figure 5:**
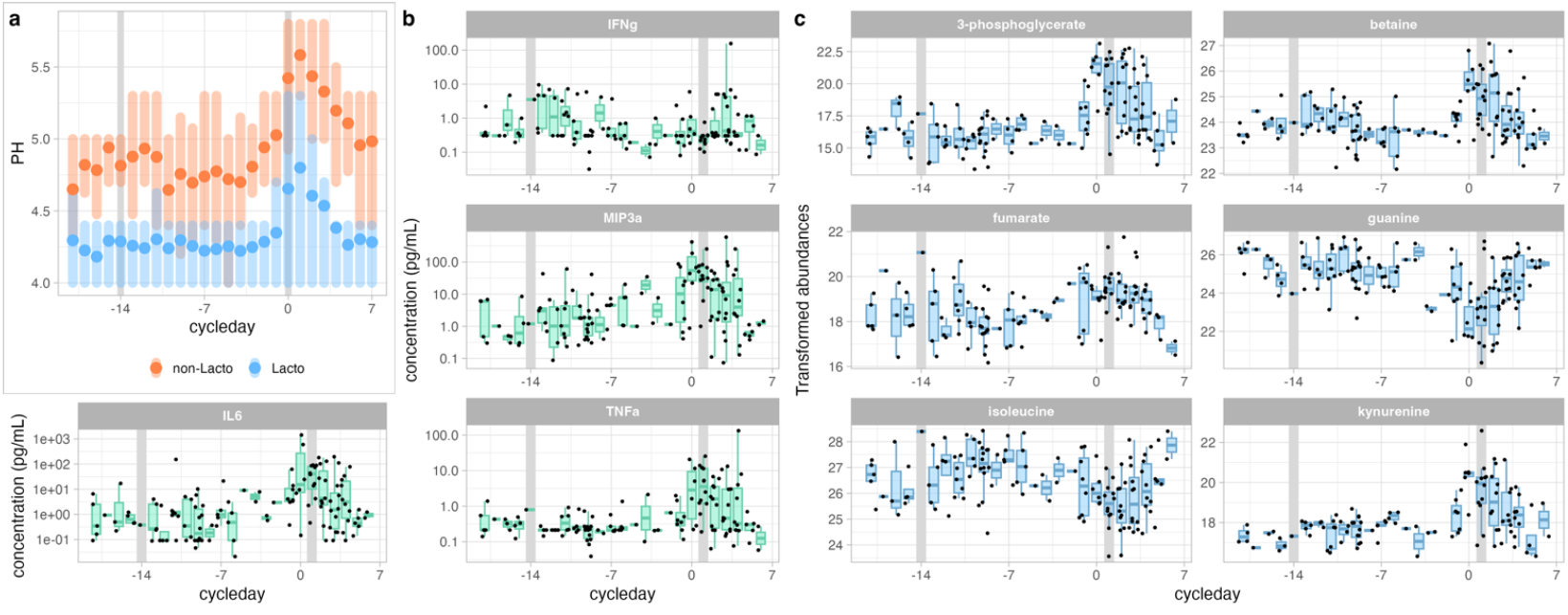
Vaginal pH, cytokines, and metabolites throughout the menstrual cycle. **(a)** Distribution of vaginal pH throughout the menstrual cycle in *Lactobacillus*-dominated samples (blue) and non-*Lactobacillus*-dominated samples (orange). Dots indicate the means, while the shaded vertical bars span from the 25^th^ to the 75^th^ percentiles. **(b-c)** Concentration (y-axis) of four cytokines (panels in b) and six metabolites (panels in c) with significant concentration variations throughout the menstrual cycle (x-axis). Each black dot is a sample.

Half of the cytokines (10 out of 20, p-values < 0.01, adjusted for multiple testing) showed a significant association with the menstrual cycle. Most cytokines (e.g., IL6 or TNFα) peaked during menses, while two of them (IFNγ and IL13) showed elevated abundance about the time of ovulation (Fig 5b, Fig S3). 18% of metabolites (60 out of 336) were also significantly associated with the menstrual cycle (Fig 5c, Fig S4). Most (72%) had increased or decreased abundances in the late luteal phase or during menses (between cycle day-3 and 5, Fig S4).

## Discussion

In this study, we used topic models, a mixed membership method, to identify bacterial sub-communities within vaginal microbiota samples from both pregnant and non-pregnant US women. We identified four *Lactobacillus*-dominated sub-communities corresponding to the four *Lactobacillus*-dominated community state types (CST), and five non-*Lactobacillus* sub-communities (*i*.*e*., topics), refining the structure of samples traditionally assigned to community state type (CST) IV (18). This CST is particularly relevant clinically as a paucity of *Lactobacillus* species is associated with bacterial vaginosis (BV), increased risk of preterm birth, and susceptibility to acquiring sexually transmissible infections (3, 5, 6, 10–12, 25, 28).

These five non-*Lactobacillus* sub-communities were found to belong to two groups. One group contained three topics (IV-A, IV-B.a, IV-B.b) and was characterized by the co-occurrence of ASVs taxonomically assigned to species from the *Gardnerella, Megasphaera, Atopobium, Fastidiosipila*, and *Sneathia* genera and of *Prevotella amnii*. The other group contained two topics (IV-C0 and IV-C1). It was characterized by the co-occurrence of species from the *Corynebacterium, Finegoldia, Peptoniphilus, Bifidobacterium, Staphylococcus*, and *Streptococcus* genera, and of *Prevotella bivia/denticola* and *timonensis*. These two groups align with sub-groups previously identified with a clustering approach that aimed to identify reference community state types in non-pregnant women from a large collated dataset (20): sub-CST IV-A and B belong to the first group of 3 topics, and sub-CSTs IV-C0-4 to the second group. This study thus confirms that non-*Lactobacillus*-dominated microbiotas present sub-structures that may have clinical relevance.

The main difference between the approach used here (topic analysis) and clustering approaches traditionally used to identify sub-groups in the vaginal microbiota lies in the *mixed membership* nature of topic models, thereby allowing samples to be associated with multiple topics in different proportions. This property offers the advantage of revealing longitudinal transitions between sub-communities and the rate at which they occur, which is impossible with clustering approaches. We showed here that, in pregnant participants, stable microbiotas were almost equally well characterized by clusters and topics; in contrast, unstable microbiotas composition was better represented by mixed topic memberships than by sub-CSTs. We also observed that topic memberships could better predict the risk that a participant’ s microbiota would lose its *Lactobacillus* dominance and switch to a sub-optimal microbiota composition.

In this study, we compared topic- and clustering-based sample descriptions in cases in which sub-communities (mixed) memberships were used as explanatory variables; the actual microbiota composition or the risk of losing *Lactobacillus* dominance were our response variables. We expect that colleagues might also find advantages in using sub-communities mixed memberships (topic-based sample description) as a *multivariate response variable* to identify host or intervention related factors associated with specific transitions or intermediate states. In contrast to univariate alternative or clustering, this might better reflect the potential multiple etiologies of vaginal dysbiosis.

Another difference between topic models and clustering approaches is that topic models allow for “synonyms”, which may reflect *potential functional equivalences* in a microbial community context. Indeed, if two species are found interchangeably (but not simultaneously) with a specific combination of other species, these two species will be found in the same topic. In contrast, clustering approaches tend to create two clusters, one containing each of these two species, potentially artificially increasing the number of functionally relevant sub-communities. This matches our observations as a single topic encapsulates four sub-CSTs (IV-C1-4) (20) characterized by four mutually exclusive taxa that co-occur with the same set of other species. These four taxa belong to the genera *Streptococcus, Enterococcus, Bifidobacterium*, and *Staphylococcus* and these sub-communities are found with higher prevalence in non-pregnant individuals, often during menses.

Topic models used in this study are unsupervised methods, and, like clustering, topic models identify dataset-specific features. This means that sub-communities identified in samples from a different cohort may differ from those identified in this study. However, we expect these sub-communities to be reproducibly observed in other (North American) populations since the sub-communities revealed by our analysis were found in individuals from three distinct cohorts, encompassing both pregnant and non-pregnant individuals. Further, the agreement between the topic composition and the composition of sub-CSTs, which had been identified from non-pregnant individuals’ samples, supports the generalizability of our findings. Deeper sequencing methods (*e*.*g*., metagenomics) may allow a more precise taxonomic characterization of microbiota samples, which may, in turn, enable further refinement of these sub-communities.

We found several associations between these subcommunities and the demographic characteristics or reproductive status of participants. Specifically, Black women were more likely to have a microbiota containing *L. iners* (topic III) and non-*Lactobacillus* subcommunities from the first group (topics IV-A, IV-B.a, and IV-B.b). Regarding differences associated with participants’ reproductive state, non-*Lactobacillus* topics from the second group (topics IV-C0 and IV-C1) were more prevalent in non-pregnant individuals than in pregnant women. They were especially more frequent during menses, a time characterized by elevated vaginal inflammation, as 40% of the measured cytokines had higher concentrations during menses. In pregnant individuals, topic IV-C1 showed a mildly significant association with the risk of preterm birth. It remains to be investigated whether vaginal inflammation is also elevated in pregnant individuals with a higher abundance of this sub-community. Our available data did not allow us to answer this question.

Almost all non-pregnant participants with data available for two menstrual cycles showed a high between-cycle correlation in their vaginal microbiota variation. Most topics or taxa, however, reached their maximal relative abundance at different menstrual cycle phases in different individuals. These inter-individual differences may be an artifact of the compositional nature (*i*.*e*., relative abundances) of our data or could be due to (i) inter-individual differences in menstrual timing (for example, one participant might have a 10-day luteal phase while another one might have a 14-day luteal phase); (ii) inter-individual differences in hormone levels (or the rates of change in these levels); or (iii) the set of species present in each individual and how each of these species might respond differently to the menstrual cycle while competing for resources. Future clinical studies including hormonal measurements would allow a better understanding of the relationships between hormonal changes and microbiota composition. Similarly, additional data would be necessary to understand if abrupt hormonal changes, the presence of blood, or the use of menstrual protections such as pads or tampons drive the substantial changes in vaginal microbiota composition observed during menses.

These abrupt changes in microbiota composition around menses were accompanied by changes in vaginal cytokine and metabolite levels. As mentioned above, 8 out of 20 measured cytokines had elevated levels during menses (and 2 around ovulation), and 70% of the 60 metabolites that varied with the menstrual cycle peaked or dropped during menses. For example, kynurenine peaked during menses while isoleucine dropped. Kynurenine is a tryptophan catabolite via a pathway involving IDO1-mediated degradation. It is known to play a role in blood vessel dilatation during inflammatory events (29). The elevated levels of kynurenine during menses found in our study are thus consistent with these roles and with past studies showing varying levels of kynurenine in serum and urine through the cycle (30, 31). In our vaginal samples, isoleucine, a branched-chain amino acid with important metabolic functions (32), was found with the highest levels in the luteal phase and lowest during menses. Interestingly, serum levels of isoleucine show opposite trends (33). The menstrual changes in cytokine concentrations were consistent with those identified in previous studies in non-pregnant individuals (34, 35). In addition to the 8 cytokines that had elevated levels around menses, IFNγ and IL13 had elevated levels around ovulation. Further studies in which the clinical and reproductive state of participants is more accurately measured would allow one to confirm these findings and to further investigate the associations between local inflammation and microbiota composition.

## Conclusions

Topic analysis revealed bacterial sub-communities (topics) shared across pregnant and non-pregnant women, confirming the existence of sub-structures in non-*Lactobacillus*-dominated microbiota and their possible clinical relevance. Compared to clustering approaches traditionally used to categorize microbial composition, topics provide an expanded characterization of the heterogeneity of the previously described risk-associated community state type IV (CST IV), a high-resolution view of transitions between communities, and they better predict the loss of *Lactobacillus* dominance. We found that the menstrual cycle had a strong impact on the vaginal microbiota and on vaginal levels of 60 metabolites and half (10/20) of the measured cytokines. Specifically, one sub-community with increased prevalence during menses, a time of elevated vaginal inflammation, was also mildly associated with the risk of preterm birth. *In vitro* studies will provide further functional insights into the identified sub-communities, their ecological network, and their effects on the vaginal epithelium.

## Material and Methods

### Cohorts and sample collection

#### Daily samples from non-pregnant participants

The samples were obtained from 30 participants recruited at the University of Alabama, Birmingham (UAB) as part of the UMB-HMP study, which enrolled participants regardless of their BV diagnosis between 2009 and 2010 (15). Participants with symptomatic BV were treated using standard-of-care practices (15). These 30 participants were selected to represent women with stable *Lactobacillus*-dominated microbiota, stable non-*Lactobacillus*-dominated microbiota, and unstable microbiota (i.e., with samples dominated by *Lactobacillus* and others dominated by non-*Lactobacillus*). Each participant self-collected daily vaginal swabs for 10 weeks, resulting in a maximum of 10 × 7 = 70 samples per individual. For further detail about recruitment criteria and sample collection, see (15).

#### Weekly samples from pregnant women

We used the samples from both cohorts presented previously (4). 39 pregnant individuals were recruited at Stanford University (SU), and 96 pregnant individuals were recruited at the University of Alabama, Birmingham (UAB) between 2013 and 2015. Pregnant participants from both cohorts were enrolled from the fourth month of their pregnancy (earliest enrollment at week 8, latest at week 22), and vaginal swabs were collected weekly (approximately) until delivery. There was an average of 16 samples per participant and 2179 samples in total. The distributions of age, BMI, and race were significantly different between the two cohorts (Table S1). Participants recruited at UAB were part of a pool of individuals for which intramuscular progesterone injections (17-OHPC) were indicated or recommended. UAB participants received that treatment throughout pregnancy. The treatment is intended to reduce the risk of preterm birth in pregnant women with a singleton pregnancy and who have a history of singleton spontaneous preterm birth. 9/39 (23 %, SU) and 41/96 (43 %, UAB) participants delivered preterm, defined as a delivery before 37 weeks of gestation.

#### Metabolite and cytokine samples

Metabolites and cytokine concentrations were quantified in a subset of the non-pregnant samples. Specifically, 5 samples per non-pregnant participant were selected such that they were separated by approximately 2 weeks. In addition, 5 samples each were from 10 additional non-pregnant participants of the UMB-HMP study but recruited at different sites (Emory University and the University of Maryland Baltimore). In total, metabolites and cytokines were quantified in 200 samples from 40 non-pregnant individuals.

### Ethics

All participants provided written informed consent. Ethical approval was obtained from the Institutional Review Boards of Stanford University (IRB protocol no. 21956), the University of Alabama (protocol no. X121031002), Birmingham, Emory University, and the University of Maryland Baltimore. All research was conducted in compliance with relevant guidelines and regulations.

### Vaginal microbiota sequencing

#### Daily samples from the 30 non-pregnant participants recruited at UAB (1534 samples)

The V3-V4 regions of the 16S rRNA gene were amplified and then sequenced with the Illumina HiSeq/MiSeq platforms.

#### Weekly samples from pregnant participants of both cohorts (SU and UAB) (2179 samples)

Raw sequence data from samples from pregnant participants were generated and processed as described in (4). In brief, genomic DNA was extracted from vaginal samples using a PowerSoil DNA isolation kit (MO BIO Laboratories). Barcoded primers 515F/806R (36) were used to amplify the V4 variable region of the 16S rRNA gene from each sample. Pooled amplicons were sequenced on the Illumina HiSeq platforms at the Roy J. Carver Biotechnology Center, University of Illinois, Urbana-Champaign.

Demultiplexed raw sequence data from Illumina HiSeq/MiSeq were resolved to amplicon sequence variants (ASVs) as described in the DADA2 Workflow for Big Data (https://benjjneb.github.io/dada2/bigdata.html) (37).

#### Taxonomic assignment

Automated taxonomic calls were made using DADA2’ s implementation of the RDP naive Bayesian classifier (38) and a Silva reference database (version 132) (39). The assignment of sequences of the most abundant ASVs were refined and standardized by using BLAST and NCBI RefSeq type strains. This is the case for *Lactobacillus, Candidatus* Lachnocurva vaginae (previously referred to as BVAB1), *Gardnerella*, and *Megasphaera lornae* species-level assignments, following recently published work on these species (40, 41). *Gardnerella* ASVs were tagged as G1, G2, or G3 *sensu* (4) based on exact matching of the ASV sequences. Tables with the taxonomic assignments are available (see data availability section).

#### Taxonomic agglomeration of ASV counts

ASV counts were aggregated based on their taxonomic assignment such that the counts of ASVs with the same taxonomic assignment were summed.

### Metabolite concentration quantification

Untargeted metabolomics was performed on 200 non-pregnant participant samples by ultra-high-performance liquid chromatography/tandem mass spectrometry (Metabolon, Inc.). Metabolite identification was performed at Metabolon based on an internally validated compound library, and results were expressed in relative concentrations, following the same protocol as in (42). All samples were shipped and analyzed in a single batch.

#### Data transformation

We transformed the raw metabolite relative concentrations using a variance stabilizing method (43). Raw data included the concentrations of 853 metabolites. However, the abundance of 517 metabolites was missing in more than 50% of the samples. We removed these metabolites from the analysis. Despite this, measurements for most of the remaining 336 metabolites were still missing in at least one sample. Metabolites might be missing because their abundance was lower than the detection limits or because the overall quality of a sample was lower. A sample with more than 60% missing metabolites was further excluded for the rest of the analysis.

### Cytokine concentration quantification

Vaginal cytokines were quantified in the 200 non-pregnant participant samples using a Luminex-based assay with a custom kit of 20 analytes (IFNγ, IL-1a, IL-1b, IL-4, IL-5, IL-6, IL-8, IL-10, IL-12p70, IL-13, IL-17, IL-21, IL-23, IP-10, ITAC, MIG, MIP-1a, MIP-1b, MIP-3a, and TNFα) following the same protocol as in (12). The assay was run on a Luminex FLEXMAP 3D instrument. For measurements that were below the limit of quantification for a given cytokine, values were imputed at half the lower limit of quantification (LLOQ / 2). For measurements that were above the limit of quantification for a given cytokine, values were imputed as equal to the upper limit of quantification (ULOQ). Values reported here represent medians of two technical replicates. The medians were calculated after imputation in one or both replicates (if necessary), as described above. Missing cytokine values represent technical failures of the assay for that analyte.

#### Data transformation

Raw cytokine abundances were log-transformed. Raw data included the abundance of 20 cytokines. Most of the cytokines could be quantified (11/4000 data points were missing).

### Data integration into a multi-assay experiment (MAE) object

All analyses were performed in the R software environment (44). Specific packages used for the analyses are referred to in the next sections. The raw datasets were loaded and minimally processed before being formatted into SummarizeExperiment objects of the SummarizedExperiment bioconductor package (45), then combined into a single S4 object using the MultiAssayExperiment bioconductor package (46).

### Identifying bacterial sub-communities using topic analysis

Microbial communities were estimated based on LDA (latent Dirichlet allocation) (22, 23). LDA models were fitted to the data for K (the number of topics) = 1 to 25 using the R package “topicmodels” (47). Models were fitted on the taxonomically agglomerated ASV counts directly, without any prior normalization; the library size being one of the parameters of this Bayesian framework.

Topics were aligned across K using the topic alignment method described in (24). To identify robust topics across K, we used the alignment summary scores for topic coherence as defined in the same reference.

### Comparison of topic composition with subCST composition

Both sub-CSTs centroids (20) and topics are described as compositional data: for each sub-CST or topic, the proportion of each species is provided such that the proportions sum to one per sub-CST/topic. However, the taxonomic assignment used by France et al. (20) differs from the assignment used here. For example, sub-CSTs taxonomy does not differentiate between *Gardnerella* species or uses “BVAB1” when we use *Ca*. Lachnocurva vaginae. Consequently, to compare topics with sub-CSTs, we proceeded in two steps. First, we harmonized the taxonomic assignments between the two methods (*e*.*g*., proportions of the different *Gardnerella* species were aggregated). A dictionary of the matched taxonomic assignment is available in the supplementary material. We then computed the Bray-Curtis dissimilarity between the composition of each topic and sub-CST centroid.

### Assignment to Valencia reference sub-CST

Per France et al. (20), samples were assigned to the sub-CST that maximizes the Yue and Clayton similarity between the sample composition and the sub-CST centroids.

### Microbiota composition prediction from sub-CST and topic membership

To compare how well sample composition was represented by sub-CST categories (fixed composition) or topics (fewer topics than sub-CSTs, but mixed memberships), we compared the Bray-Curtis dissimilarity between the actual sample compositions and the sample compositions predicted by topic mixed memberships or by sub-CST membership. The predicted composition of a sample is either the composition of the centroid of the sample’ s sub-CST or the average of topics composition (displayed in figure 2b) weighted by the proportion of each topic in that sample (*i*.*e*., 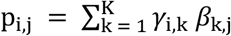 where p_i,j_ is the proportion of taxa j in sample i, k is the topic index going from 1 to K, the total number of topics, *γ*_i,k_ is the proportion of topic k in sample i, and *β*_k,j_ is the proportion of taxa j in topic k).

### Microbiota local stability

Samples were classified as belonging to a stable microbiota if they were part of a series of 5 consecutive samples with a Bray-Curtis dissimilarity smaller than a given threshold. Otherwise, the microbiota was considered unstable.

### Predicting the risk of losing Lactobacillus dominance

To predict the risk of losing *Lactobacillus* dominance at the next time-point in participants’ longitudinal time series, a logistic regression model was fitted to the data. The explanatory variables were either the sub-CST category of the sample or the topic proportion at the current time point. The response variable was a binary variable indicating if the next sample belonged to a *Lactobacillus*-dominated sub-CST or not. *Lactobacillus* dominance was defined as a total proportion of *Lactobacillus* larger than 50%. The models were fitted on a training set (a random sample comprising 80% of the total dataset) and prediction performances were evaluated on the remaining 20% of the dataset. The procedure was repeated independently 10 times. Because the loss of *Lactobacillus* dominance is rare (10% of cases), we weighted the sample to give more weight (10 folds) to the minority class when training the models, and we used the F1 score, the harmonic mean between precision and sensitivity, to evaluate predictive performances. To test for differences in the sub-CST-vs topic-based prediction performances, a non-parametric Wilcoxon Rank sum test was used.

### Associations between topic composition and demographic variables

A Dirichlet regression was used to test if race, study site, or pregnancy were associated with differential topic proportions. Because most participants’ race was Black or White, the race was transformed into a three-category variable: Black, Other, and White, with “Other” serving as the reference. Pregnancy was a binary variable (pregnant *vs*. non-pregnant), and so was the study site: Stanford University (SU) *vs*. University of Alabama Birmingham (UAB). The model used is *p* = *β* + *α*_R_R + *α*_P_P + *α*_S_S + ε where *p* is the vector of topic proportions lying on the K-dimension simplex. Coefficients were obtained using the DirichletReg package in R (48).

### Identification of phases of the menstrual cycle

Menstrual cycles were identified from bleeding flows reported daily by participants on a scale from 0 (none) to 3 (heavy). A hidden semi-Markov model was specified to account for empirically observed distributions of cycle length and bleeding patterns across the menstrual cycle, including spotting between menses (49). Data of participants who reported too few days with bleeding (i.e., less than 3/70 study days) or too many (i.e., more than 30/70 study days) were excluded from the menstrual cycle analyses. Once cycles were identified (see Fig S5), cycle days were numbered forward and backward from the first day of the period. To align the two major menstrual events (i.e., ovulation and menses) across participants and given that the luteal phase has been well documented to vary less than the follicular phase (27), cycles were standardized starting from day-18 (i.e., 18 days before the start of the next cycle) and ending on day +7 (i.e., 7 days after the first day of the menses). This definition ensures that the standardized cycles would include the days leading to ovulation, estimated to happen around days-12 to -14 (27), and allows for the best possible alignment of the two major menstrual events (ovulation and menses) in the absence of hormonal and/or ovulation markers.

### Testing for differential abundance throughout the menstrual cycle

To identify metabolites, cytokines, or topics with differential abundance (metabolites or cytokines) or differential probabilities of being present at specific phases of the menstrual cycle, a linear model (for abundances) or logistic regression (proportions) was fitted to circular splines parameterized with 4 degrees of freedom (R package “pbs”). Analysis of deviance was used to report p-values of the F-statistics and corrected for multiple testing using the Benjamini-Hochberg method.

### Associations between topic proportions and preterm birth

To test if topic proportions were associated with preterm birth, a logistic regression model was fitted on the data. Explanatory variables were the per-participant topic proportion averages, and the response variable was a binary variable indicating whether participants delivered preterm or not.

### Correlation in vaginal microbiota composition between two consecutive cycles

To evaluate how the menstrual cycle affects the vaginal microbiota composition, we compute the RV coefficient (50) and associated permutation test p-value (51) between the topic or taxa proportions of the first cycle and of the second cycle. To quantify the magnitude of change in microbiota composition throughout the cycle (x-axes of fig 4b), we first compute the average topic or taxa proportion across cycles for each cycleday. Then, the pairwise Bray-Curtis dissimilarities are computed so that the compositions of each cycleday are compared against each other. The maximum value is used to quantify the magnitude of change throughout the menstrual cycle for each participant.

### Availability of data and materials

The sequence data for samples from non-pregnant study participants are available in the NCBI Sequence Read Archive (SRA) under BioProject accession numbers <u>PRJNA208535</u> (samples beginning with UAB) and <u>PRJNA575586</u> (samples beginning with AYAC and EM). Sequence data from samples from pregnant study participants are available on the SRA (accession no. <u>PRJNA393472</u>). The raw data and R code enabling the reproduction of the analyses are available at https://purl.stanford.edu/gp215vr4425. The code is also provided in the SI.

## Acknowledgments

The authors thank Anna Robaczewska for generating the amplicon libraries, and Nomfuneko Mafunda, Brooke Spencer, and Leah Froehle for running the cytokine quantification assays. They also thank Dr. K. Sankaran and Dr. J. Fukuyama for advice about the visualization of topic analyses, and all other members of the VMRC consortium for fruitful discussions and interactions.

