## Supplementary Material for "Sub-communities of the vaginal microbiota in pregnant and non-pregnant women"

Supplementary Information

L. Symul, P Jeganathan, E. Costello, M. France, S.M. Bloom, D.S. Kwon, J. Ravel, D.A. Relman, S. Holmes

Last updated on 28 June, 2023

This document is the pdf-rendering of the analyses performed in R for the manuscript "Sub-communities of the vaginal ecosystem in pregnant and non-pregnant women". It contains supplementary figures and tables referred to in the main text, the code and workflow of the analyses, and many additional figures and tables that the reader might find useful. For reproducibility purposes, the last section documents the softwares and packages versions used for the analyses.

**Reproducibility Receipt** **88**

5 Identified and included standardized menstrual cycles from bleeding data. Each line is a non-pregnant participant, each dot is a day. Color indicates bleeding intensity from 0 to 3. Crossed days are days excluded for menstrual-cycle related analyses. Days are excluded either because cycles could not be identified from the participant's data or because they fall outside identified standardized menstrual cycles. Small blue dots indicate days that were identified as part of a 'menstrual period'. . . . 8
6 Distribution of total ASV counts per 16S rRNA sequencing assay. . . . 11
7 Batch effect between the metabolites concentrations in the samples of the Pregnant vs non-Pregnant subjects and the metabolomics batches . . . 12
8 Distribution of the proportion of Lactobacillus in samples from Pregnant and Non-pregnant participants. . 17
9 Time-series of the proportion of Lactobacillus species in non-pregnant subjects. . . . 18
10 The frequency of missing data is higher for metabolites with lower abundances. . . . 19
11 Correlation between lactate concentrations and proportion of Lactobacillus . . . 20

12 Time-series of subjects with missing data for lactate. . . . 21
13 Metabolites samples quality in each cohort. . . . 22
14 Histogram of transformed cytokine abundances. . . . 24
15 Cytokines distributions . . . . 24
16 Bleeding intensities reported by non-pregnant subjects. . . . 28
17 Graph of the hidden semi-Markov model (HSMM) used to detect menstrual cycles from bleeding records. . 29
18 Specified sojourn distribution for our menstrual cycle HSMM . . . . 30
19 Fitting the HSMM: likelihood over the EM steps. . . . 30
20 Sojourn distributions of the fitted HSMM. . . . 31
21 Identified and included standardized menstrual cycles from bleeding data. . . . 34
22 pH throughout the menstrual cycle (boxplot) . . . . 35
23 pH throughout the menstrual cycle (ribbon) . . . . 35
24 pH and proportion of Lactobacillus . . . . 37
25 Nugent score throughout the menstrual cycle . . . . 37
26 Nugent score and proportion of Lactobacillus . . . . 38
27 Alignment diagnostic scores. Colors denote the various alignments (several alignments were performed for increasing max K). . . . . 44
28 Topic alignments. Please note that topics might have different colors in both aligment. . . . . 46
29 Topic composition for the optimal K and a coarser K . . . . 47
30 Topic alignment colored by paths (a), topic coherence (b), or topic refinement (c). . . . . 48
31 Coherence and refinement scores of topics at two resolutions. . . . . 49
32 Composition of Valencia centroids. Adapted from France et al, 2020. . . . . 50
33 Bray-Curtis dissimilarity between Valencia centroids and topics. . . . . 52
34 Bray-Curtis dissimilarity between the actual sample compositions and those predicted by topic mixed membership or sub-CST membership. . . . . 54
35 Distribution of the differences between the Bray-Curtis dissimilarity between actual sample compositon and that predicted by sub-CST and topics for samples in stable microbiota episodes or samples belonging to transitions or unstable microbiotas. . . . . 56
36 Prediction metrics (panels) for each method (x-axis) . . . . 60
37 F1 score per sub-CST for each method. . . . . 61
38 Predicted risk of losing lactobacillus dominance per method and CST. . . . . 62
39 Topic composition per racial groups. . . . . 64
40 Topic composition per cohort (location and pregnancy status). . . . . 65
41 Topic distribution per status in all subjects. . . . . 69
42 Topic distribution per status in all subjects. . . . . 70
43 Topic distribution per Race in all subjects. . . . . 71
44 Metabolites associated with the menstrual cycle. . . . . 84
45 Metabolites with a strong association with the menstrual cycle. . . . . 86
46 Cytokines whose concentration varies with the menstrual cycle. . . . . 87

### 1 Supplementary tables and figures

Table 1: Demographic characteristics of study participants by cohort. UMD stands for University of Maryland (UMD-led cohort, but participants recruited at UAB), SU for Stanford University, and UAB for University of Alabama, at Birmingham. The distributions of all demographical variables are significantly different across cohorts.

| metric | category | Non-pregnant (AYAC/EM) | Non-pregnant (UAB) | Pregnant (SU) | Pregnant (UAB) |
|---|---|---|---|---|---|
| | n | 10 | 30 | 39 | 96 |
| Age | min | 28 | 19 | 25 | 17 |
| | 5th perc. | 28.9 | 20 | 26.9 | 20.7 |
| | median | 36 | 29 | 32 | 25 |
| | 95th perc. | 38.1 | 43 | 38.3 | 34 |
| | max | 39 | 45 | 42 | 38 |
| Race | Asian | 1 (10%) | 0 | 3 (8%) | 1 (1%) |
| | Black | 6 (60%) | 19 (63%) | 1 (3%) | 79 (82%) |
| | Hispanic/Latino | 1 (10%) | 3 (10%) | 0 | 0 |
| | White | 2 (20%) | 8 (27%) | 22 (56%) | 9 (9%) |
| | Other | 0 | 0 | 13 (33%) | 7 (7%) |
| BMI | unknown | 10 (100%) | 30 (100%) | 0 | 14 (15%) |
| | underweight (< 18.5) | 0 | 0 | 3 (8%) | 5 (5%) |
| | healthy weight [18.5, 25[ | 0 | 0 | 21 (54%) | 30 (31%) |
| | overweight [25, 30[ | 0 | 0 | 8 (21%) | 9 (9%) |
| | obese (> 30) | 0 | 0 | 7 (18%) | 38 (40%) |
| Delivery | extremely preterm (< 28 w) | 0 | 0 | 0 | 12 (12%) |
| | very preterm (< 32 w) | 0 | 0 | 1 (3%) | 7 (7%) |
| | moderate preterm (< 37 w) | 0 | 0 | 8 (21%) | 22 (23%) |
| | term | 0 | 0 | 30 (77%) | 55 (57%) |
| | NA | 10 (100%) | 30 (100%) | 0 | 0 |

```r
load(
  file = "../results/suppl_figs/local_stability_and_topics_vs_clusters.Rdata"
)

g_local_stability_and_topics_vs_clusters
```

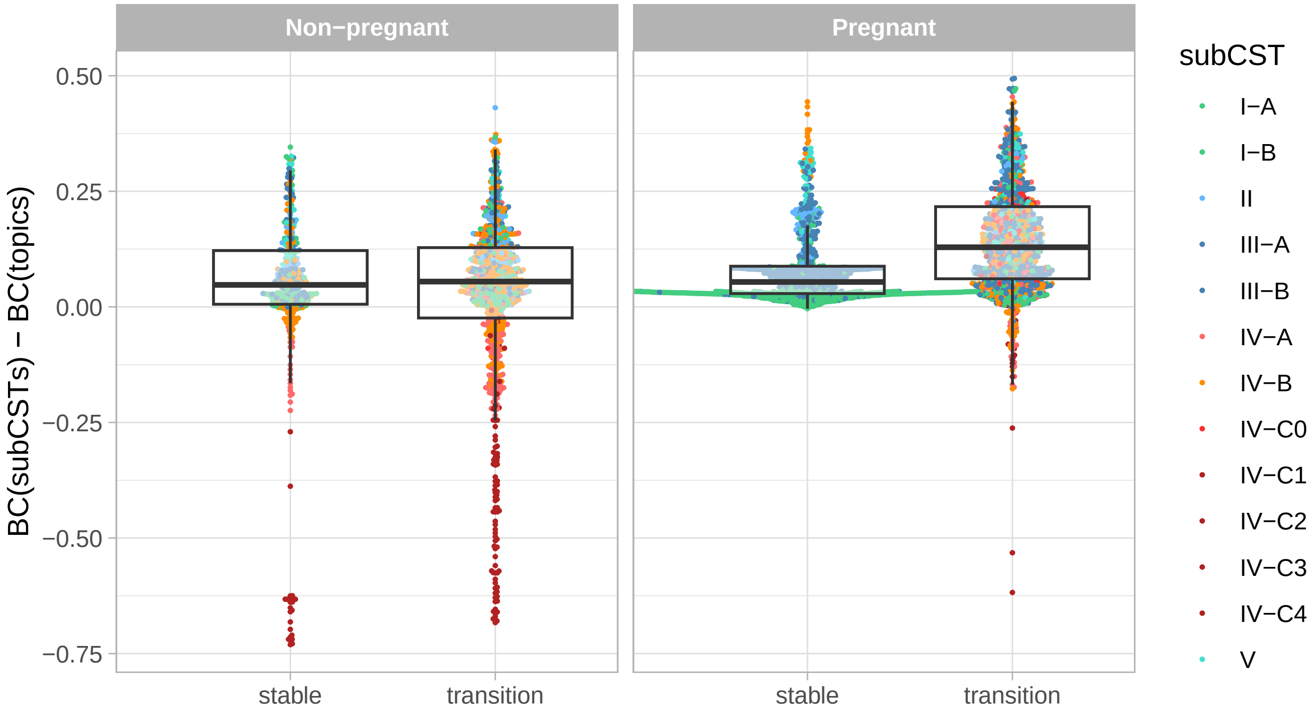

Figure 1: Distribution of the differences between the Bray-Curtis dissimilarity between actual sample compositon and that predicted by sub-CST and topics for samples in stable microbiota episodes or samples belonging to transitions or unstable microbiotas.

```
load(file = "../results/suppl_figs/table_sensitivity.Rdata")

t_sensitivity %>%
  kable(., format = "latex", booktab = TRUE, linesep = "",
        caption = "Average differences between the Bray−Curtis dissimilarity (3rd and 4th columns) between ac
  kableExtra::kable_styling(latex_option = "HOLD_position") %>%
  kableExtra::collapse_rows(columns = 1, latex_hline = "major")
```

Table 2: Average differences between the Bray-Curtis dissimilarity (3rd and 4th columns) between actual and predicted sample composition when sample composition is predicted with topic or cluster membership(s) for various thresholds (1st column) differentiating between stable microbiotas and transition states in samples from pregnant or non-pregnant participants (2nd column). The 5th column provides the p-value from a one-sided t-test.

| threshold | Status | mean BC diff. (stable microbiotas) | mean BC diff. (transitions) | p-value |
|---|---|---|---|---|
| 0.15 | Non-pregnant | 0.0453 | 0.0225 | > 0.1 |
| | Pregnant | 0.0638 | 0.1421 | ≤ 0.001 |
| 0.25 | Non-pregnant | 0.0300 | 0.0231 | > 0.1 |
| | Pregnant | 0.0832 | 0.1422 | ≤ 0.001 |
| 0.35 | Non-pregnant | 0.0436 | 0.0042 | > 0.1 |
| | Pregnant | 0.0999 | 0.1361 | ≤ 0.001 |

```
load(file = "../results/suppl_figs/pred_metrics.Rdata") #
g_pred_metrics
```

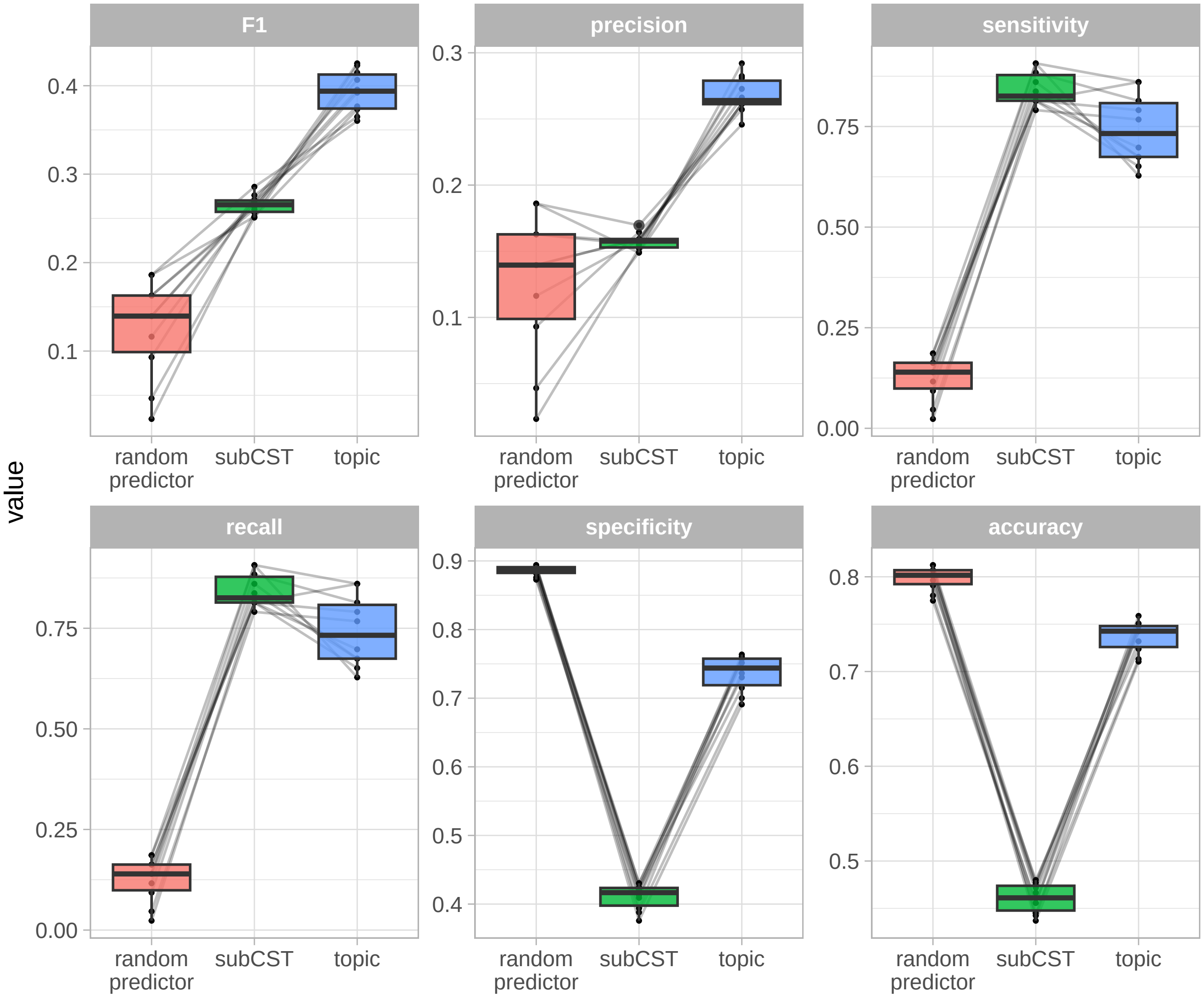

Figure 2: Prediction metrics (panels) for each method (x-axis)

```
load(file = "../results/suppl_figs/I_MC.Rdata")

g_I_MC
```

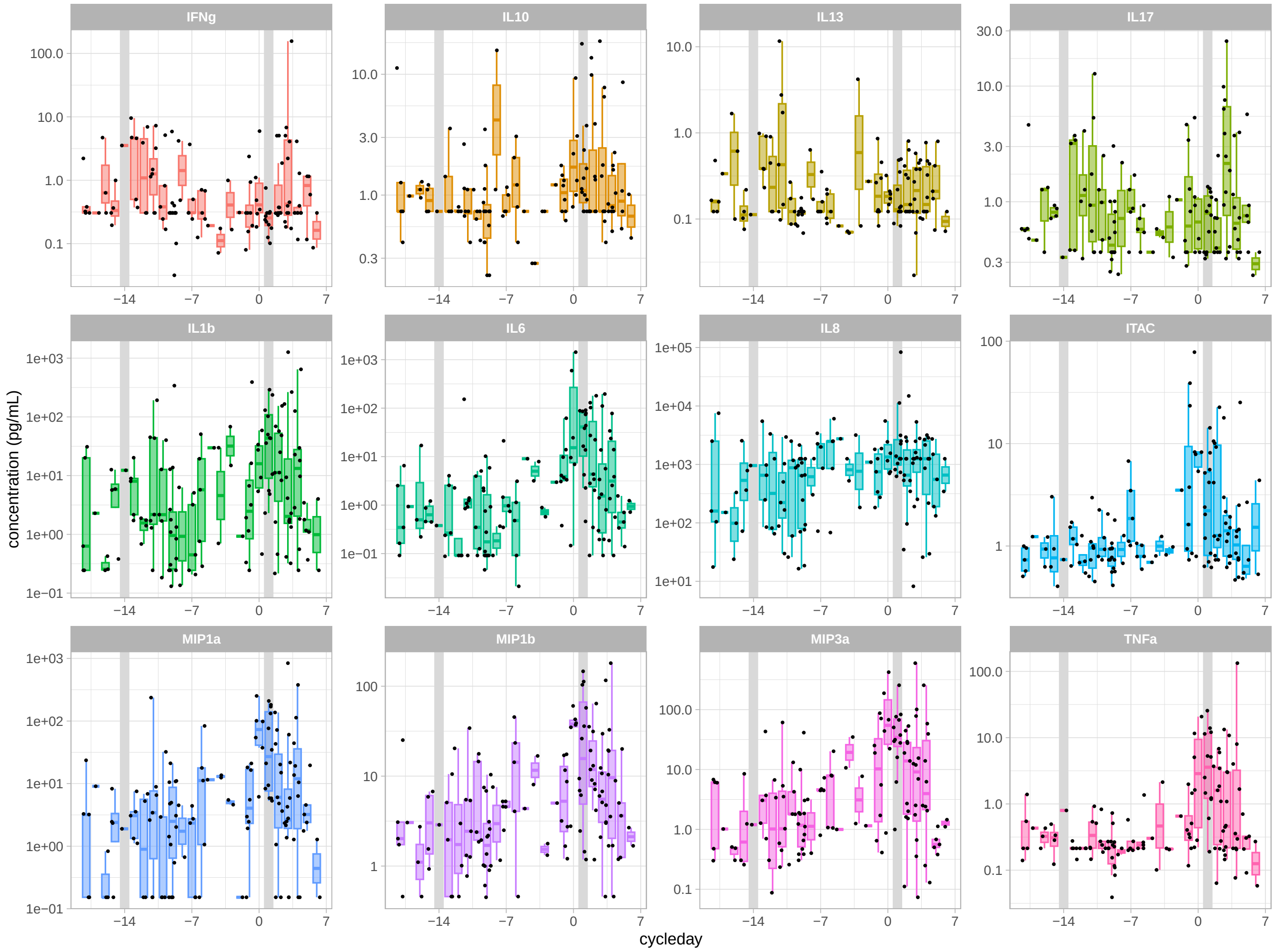

Figure 3: Cytokines whose concentration varies with the menstrual cycle.

```r
load(file = "../results/suppl_figs/MB_MC.Rdata")

g_MB_MC
```

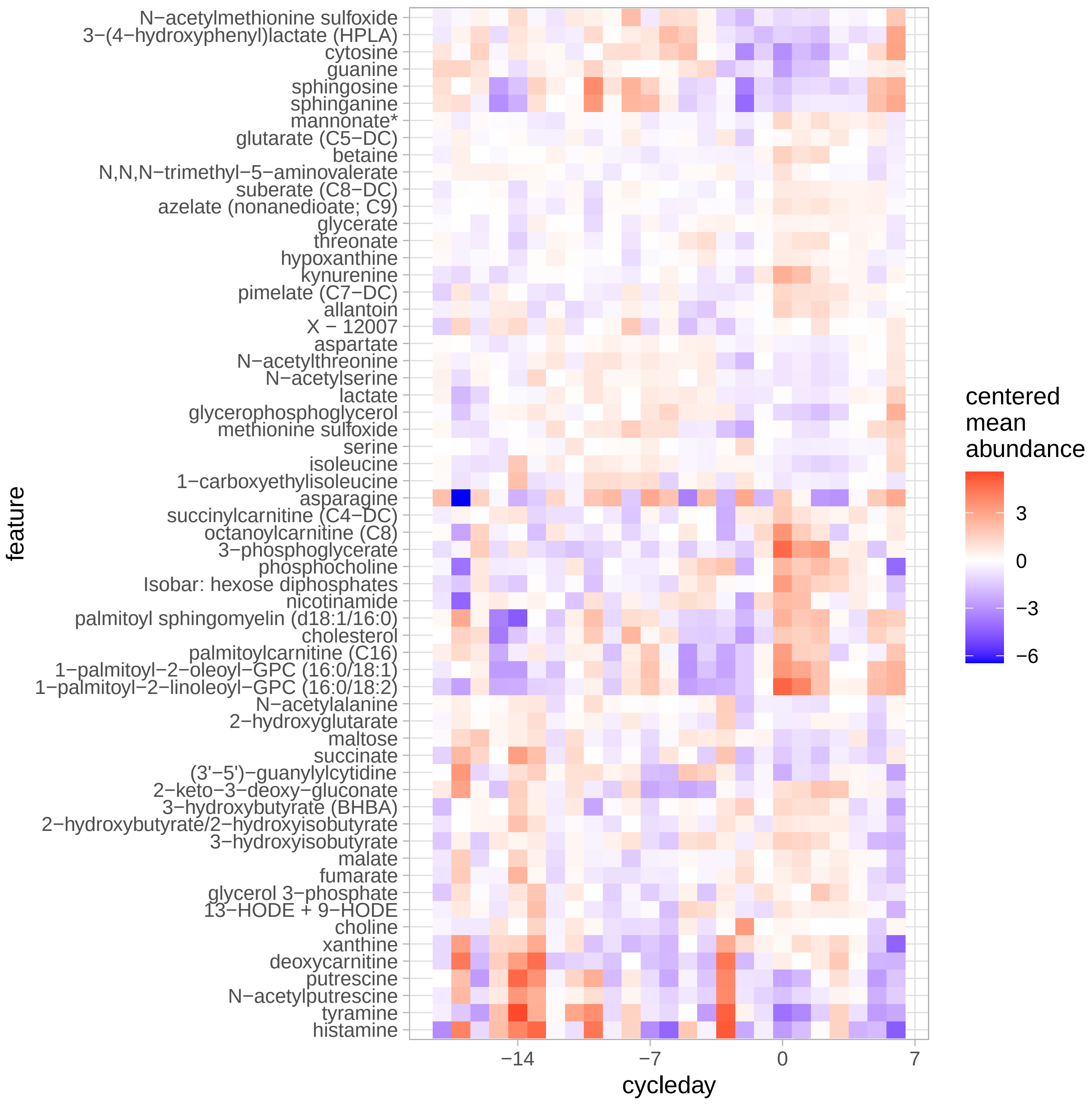

Figure 4: Metabolites associated with the menstrual cycle.

```
cycle_file <- "../results/suppl_figs/cycles.Rdata"

if (file.exists(cycle_file)) {
  load(cycle_file)
  g_cycles
}
```

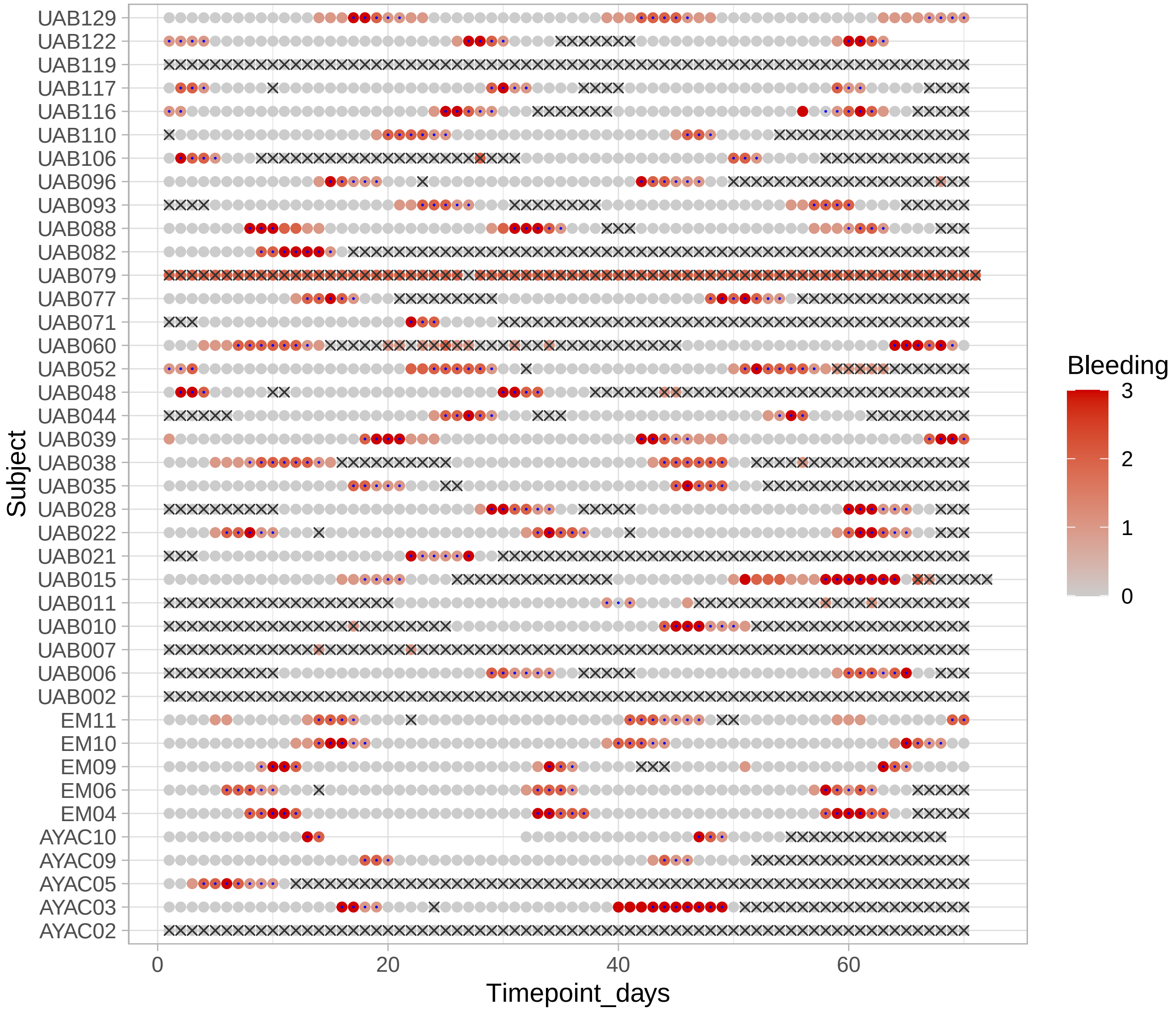

Figure 5: Identified and included standardized menstrual cycles from bleeding data. Each line is a non-pregnant participant, each dot is a day. Color indicates bleeding intensity from 0 to 3. Crossed days are days excluded for menstrual-cycle related analyses. Days are excluded either because cycles could not be identified from the participant's data or because they fall outside identified standardized menstrual cycles. Small blue dots indicate days that were identified as part of a 'menstrual period'.

### 2 Data Preparation

In this section, we first describe how the samples have been collected and quantification methods. We then pre-process each assay and store them in a `SummarizedExperiment` object. These assays are then collected in a `MultiAssayExperiment` object. Finally, the data is augmented with various variables that are useful for downstream analyses.

#### 2.1 Data description: Groups, Cohorts, and Sites; Domains and Assays.

**Groups (Status), Cohorts and Sites**: Vaginal swabs were collected from two groups in three cohorts:

- one cohort of 40 non-pregnant subjects recruited at 3 sites (UAB, EM, and AYAC) and whose data were provided by the Ravel's lab (UMD),
- two cohorts of 135 pregnant subjects in total recruited at the Stanford center (39 subjects) or at the Alabama center (96 subjects).

**Domains**: Some of these swabs were then processed for three domains:

- microbiota composition (16S rRNA sequencing),
- metabolite relative concentrations (targeted metabolomics by Metabolon)
- cytokine concentrations (Luminex assay).

**Sampling frequency**: Swabs were collected daily for ten weeks in non-pregnant women and approximately weekly throughout pregnancy for pregnant subjects. Among all of these samples, 5 samples per subjects from 40 pregnant participants (Stanford and UAB) and the 40 non-pregnant participants were selected and labeled as "VMRC" samples.

**16S rRNA Assays**

Swabs were sequenced using different technologies or protocols:

| Name | Status | Cohorts | Resolution | 16S rRNA region | N subjects | N samples |
|---|---|---|---|---|---|---|
| S2017 | Pregnant | Stanford and UAB | ASV | V3 | 135 | 2179 |
| UMD-ASV | Non-pregnant | UMD | ASV | V3V4 | 30 | 1534 |
| UMD-OTU | Non-pregnant | UMD | OTU | V1V3 | 10 | 747 |
| VMRC | both | Stanford and UMD | ASV | V3 | 80 | 393 |

For the topic analysis, we use the data from the `S2017` and `UMD-ASV` assays.

**Metabolite Assays**

Metabolite relative concentrations in the VRMC samples were quantified by Metabolon.

**Cytokine Assays**

Cytokines from the VRMC samples were quantified using a Luminex-based assay with a custom kit of 20 analytes (IFNg, IL-1a, IL-1b, IL-4, IL-5, IL-6, IL-8, IL-10, IL-12p70, IL-13, IL-17, IL-21, IL-23, IP-10, ITAC, MIG, MIP-1a, MIP-1b, MIP-3a, and TNFa). The assay was run on a Luminex FLEXMAP 3D instrument.

- Samples were run on various dates in 2019 and 2020 by technicians in the lab depending on date of receipt and kit availability from Milliplex. The technicians were Nomfuneko Mafunda, Brooke Shields, and Leah Froehle.
- Values of measurements that were **below the limit of quantification for a given cytokine** were imputed at half the lower limit of quantification (LLOQ / 2).
- Values of measurements that were **above the limit of quantification for a given cytokine** were imputed as equal to the upper limit of quantification (ULOQ).
- Values reported here represent medians of two technical replicates. The medians were calculated values were imputed for one or both replicates (if necessary) as described above.
- There are a couple of samples for which a given cytokine has a value of `NA`. These represent technical failures of the assay and should be regarded as genuine missing data (i.e., they were purely stochastic technical failures of an analyte for that sample and therefore should not introduce any systematic biological bias).

#### 2.2 SummarizedExperiment (SE) objects for each assay

SummarizedExperiment (SE) objects consist of three tables:

1. `assay` data: each column is a sample and each row is a feature (*e.g.*, an ASV sequence, a cytokine, or a metabolite)
2. `rowData`: "metadata" associated with the feature (*e.g.*, taxonomy tables for the ASV sequences or the KEGG pathways for the metabolites)
3. `colData`: "metadata" associated with each sample *specific to that assay* (*e.g.*, the sampling depth)

While the `colData` slot of each `SE` object contains sample-level metadata specific to each assay, we also build an additional table that contains sample-level non-assay specific metadata. Such information include the status of a participant (pregnant/non-pregnant) or the timepoint at which the samples was collected. This table will become the overarching `colData` of the `MultiAssayExperiment` (`MAE`) object that combine all of these assays in a single object (see next section).

##### 2.2.1 SE objects for the 16S rRNA assays

For each 16S rRNA assay, we load the raw data (provided as `phyloseq` objects or raw text files) to build the `assay` table (named `counts_[assay_name]`), the `rowData` (named `tax_[assay_name]` and which contains the taxonomy table) and the `colData` (named `colData_[assay_name]` and which contains the assay-specific sample information) . Because we want to use the same taxonomy for each ASV-assay (VM16S_S2017, VM16S_UMD_ASV and VRMC), we match the ASV from each ASV assay (`VM16S_S2017`, `VM16S_UMD_ASV` and `VM16S_VMRC`) to the same taxonomic databases. The result of these matches are stored in a taxonomic table common to each assay.

**16S rRNA - ASV - Pregnant - all samples (VM16S_S2017)**

```
counts_and_cols_S2017 <-
  get_counts_and_cols_S2017(file = "../data/processed_coll_date_fixed.rda")
```

**16S rRNA - ASV - Non-Pregnant - large subset (UMD-ASV)**

```
counts_and_cols_UMD_ASV <-
  get_counts_and_cols_UMD_ASV(file = "../data/UMD_Gates_V3V4_ASVdata_non_zero.csv")
```

**16S rRNA - ASV -(VRMC samples: P and N-P)**

```
counts_and_cols_VMRC <-
  get_counts_and_cols_VMRC(file = "../data/ps_vmrc_core16S_final.rds")
```

**16S rRNA - OTU - Non-Pregnant - all samples (UMD-OTU)**

```
counts_and_cols_UMD_OTU <-
  get_counts_and_cols_UMD_OTU(file = "../data/gates_UMD_16S_data_StR_CSTs_040919_MC_info.csv")
```

**Comparison of sequencing depth**

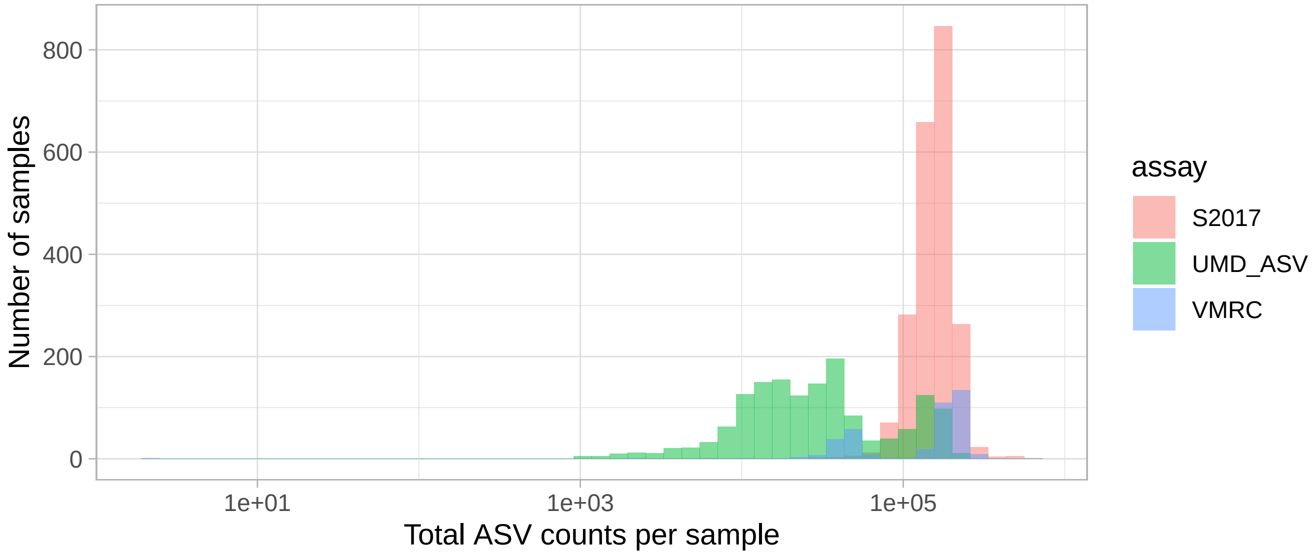

Figure 6: Distribution of total ASV counts per 16S rRNA sequencing assay.

###### Taxonomy tables

E. Costello matched the ASV sequences from each ASV assay to obtain taxonomic information for each ASV. The get_tax_tables_all_ASV_assays function load the taxonomic table, create an ASV_usename (a summary of the taxonomic information) for each ASV of each assay, and creates the taxonomy tables matching each assay.

```
tax_tables <-
  get_tax_tables_all_ASV_assays(
    tax_table_RDS_file = "../data/tax_table_draft_with_manual_annotations.rds",
    counts_and_cols_S2017,
    counts_and_cols_UMD_ASV,
    counts_and_cols_VMRC
  )

rownames(counts_and_cols_S2017$counts) <- rownames(tax_tables$tax_VM16S_S2017)
rownames(counts_and_cols_UMD_ASV$counts) <- rownames(tax_tables$tax_VM16S_UMD_ASV)
rownames(counts_and_cols_VMRC$counts) <- rownames(tax_tables$tax_VM16S_VMRC)
```

```
tax_tables$tax_VM16S_S2017 %>%
  as.data.frame() %>%
  filter(Genus == "Gardnerella", str_detect(ASV_assay_key, "G[1-3]")) %>%
  write_csv("../misc/Gardnerella_ASV.csv")
```

```
tax_VM16S_UMD_OTU <-
  get_tax_tables_UMD_OTU(counts_and_cols_UMD_OTU)

rownames(counts_and_cols_UMD_OTU$counts) <- rownames(tax_VM16S_UMD_OTU)
```

###### Summarized Experiments objects

```
se_VM16S_S2017_ASVseq <-
  SummarizedExperiment(assay = counts_and_cols_S2017$counts,
                       rowData = tax_tables$tax_VM16S_S2017,
                       colData = counts_and_cols_S2017$colData)

se_VM16S_UMD_ASV_ASVseq <-
  SummarizedExperiment(assay = counts_and_cols_UMD_ASV$counts,
                       rowData = tax_tables$tax_VM16S_UMD_ASV,
                       colData = counts_and_cols_UMD_ASV$colData)
```

```r
se_VM16S_VMRC_ASVseq <-
  SummarizedExperiment(assay = counts_and_cols_VMRC$counts,
                       rowData = tax_tables$tax_VM16S_VMRC,
                       colData = counts_and_cols_VMRC$colData)

se_VM16S_UMD_OTU <-
  SummarizedExperiment(assay = counts_and_cols_UMD_OTU$counts,
                       rowData = tax_VM16S_UMD_OTU,
                       colData = counts_and_cols_UMD_OTU$colData)
```

##### 2.2.2 SE object for the metabolomics data

In this section, we create a "SummarizedExperiment" (SE) object to store the raw metabolite abundance data. In the Data Transformation section, we will transform and impute the missing data.

Because the samples from the two groups (P and NP) were not processed exactly the same way, we expect batch effects between these two groups. In addition, samples were sent to Metabolon into two batches. We thus also expect batch effects from this additional technical difference. Because of the differences in how P and NP swabs were prepared, it looks difficult to compare the metabolite concentrations between these two groups. However, it looks like the batch effects from the Metabolon processing might be correctable. Here, we add the data from each file (i.e. P and NP) as separate assay and in the Data Transformation section, we transform, correct batch effects and impute the missing data.

```r
MB_batches <- read_csv("../data/Gates_metabolomics_batch_accounting.csv", show_col_types = FALSE)
MB_batches <-
  MB_batches %>%
  mutate(SampleID = UID %>% str_remove_all(" [0-9]*$") %>% str_replace_all(" ","_"),
         Status = ifelse(Cohort == "Stanford", "Pregnant", "Non-pregnant")) %>%
  select(Status, SampleID, Metabolomics_batch)

se_MB_P <- make_SE_for_metabolites(file = "../data/gates_metabolomics_022420_stanford.csv", batches = MB_batc
se_MB_NP <- make_SE_for_metabolites(file = "../data/gates_metabolomics_061219_UMD.csv", batches = MB_batches)
```

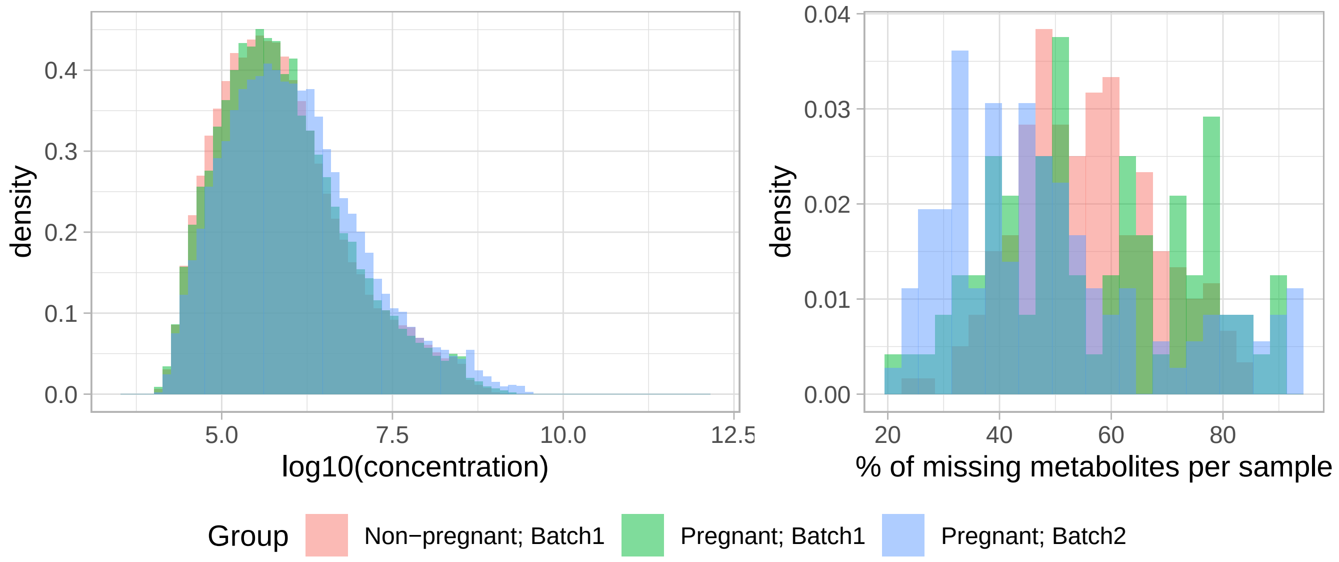

Figure 7: Batch effect between the metabolites concentrations in the samples of the Pregnant vs non-Pregnant subjects and the metabolomics batches

##### 2.2.3 SE object for the immunology data

```r
se_I <-
  make_SE_for_immunology_data(
```

```
    file = "../data/VMRC_Luminex_data_2021_01_27.txt"
  )
```

#### 2.3 SE for the clinical and survey data from the NP participants.

```
se_CS_NP <-
  make_SE_for_clinical_and_survey_data(
    file = "../data/GATES_UMD_full16S_updated_metadata_112221.csv"
  )
```

#### 2.4 MultiAssayExperiment (MAE) object combining all assays

In this section, we create a multiple component container, a MultiAssayExperiment (MAE) object that contains all SE objects created in the previous section.

**colData**: the colData for the MAE object are the Subject/Sample information specific to the Subject/Sample but not specific to any assay

```
MAE_colData <-
  combine_and_augment_MAE_colData(
    MAE_colData_P = get_ColData_P(file = "../data/processed_coll_date_fixed.rda"),
    MAE_colData_NP = get_ColData_NP(file = "../data/GATES_UMD_full16S_updated_metadata_112221.csv")
  )
```

**sampleMap** (sample map for each assay)

```
assay_names <-
  c("VM16S_S2017_ASVseq", "VM16S_UMD_OTU",
    "VM16S_UMD_ASV_ASVseq","VM16S_VMRC_ASVseq",
    "MB_P","MB_NP","I", "CS_NP")

sample_map <-
  create_sample_map(assay_names = assay_names)
```

```
## VM16S_S2017_ASVseq
## VM16S_UMD_OTU
## VM16S_UMD_ASV_ASVseq
## VM16S_VMRC_ASVseq
## MB_P
## MB_NP
## I
## CS_NP
```

**Creating the MAE object**

```
assay_list <-
  list(
    VM16S_S2017_ASVseq = se_VM16S_S2017_ASVseq,
    VM16S_UMD_OTU = se_VM16S_UMD_OTU,
    VM16S_UMD_ASV_ASVseq = se_VM16S_UMD_ASV_ASVseq,
    VM16S_VMRC_ASVseq = se_VM16S_VMRC_ASVseq,
    MB_P = se_MB_P,
    MB_NP = se_MB_NP,
    I = se_I,
    CS_NP = se_CS_NP
  )

mae <-
  MultiAssayExperiment::MultiAssayExperiment(
    experiments =
      MultiAssayExperiment::ExperimentList(assay_list),
    colData = MAE_colData,
    sampleMap = sample_map
  )
```

**MAE colData augmentation**

```
# we add if a sample has MB, I or is a VRMC sample

colData(mae)$has_MB_data =
  colData(mae)$SampleID %in% c(colnames(mae[["MB_P"]]), colnames(mae[["MB_NP"]]))

colData(mae)$has_I_data =
```

```
  colData(mae)$SampleID %in% colnames(mae[["I"]])

colData(mae)$is_VRMC_sample =
  colData(mae)$has_MB_data |
  colData(mae)$has_I_data |
  (colData(mae)$SampleID %in%  se_VM16S_VMRC_ASVseq$SampleID)

mae <- add_CS_NP_to_mae_ColData(mae)
```

###### Saving the MAE object

```
saveRDS(mae, file = "../results/mae_1_data_integration.Rds" )
```

##### 2.5 Data transformation and imputation

```
mae = readRDS("../results/mae_1_data_integration.Rds")
```

In this section, assays containing transformed and/or imputed data for each original assay are added to the MAE object.

###### 2.5.1 Agglomerating ASVs counts at the species levels

In this section, we add new assays to our MultiAssayExperiment object which contains the 16S counts agglomerated at the species levels. Topic models will be fitted to this agglomerated data. We fit topic models on the species-level counts and not on the ASV counts directly because some ASVs are only present in a few participants but may account for a large proportion of the samples in these participants. Since topic models are ignorant of the taxonomic labels, these participant-specific ASVs would very likely be found in different topics than their more widespread counterparts. So, while these participant-specific ASVs *might* represent different strains of the same species, this is more likely to bias downstream analyses correlating topics with various phenotypes or health outcomes than informing us about potential functional differences between these ASVs and more widespread ones with the same taxonomic assignments.

```
mae <- add_tax_glom_assay(mae, assay_name = "VM16S_S2017_ASVseq")
mae <- add_tax_glom_assay(mae, assay_name = "VM16S_UMD_ASV_ASVseq")
mae <- add_tax_glom_assay(mae, assay_name = "VM16S_VMRC_ASVseq")
```

###### 2.5.2 Assay combining P and NP samples (S2017 and UMD 16S assays)

We add an assay to the MultiAssayExperiment object that combines the samples from pregnant and non-pregnant participants.

```
mae <-
  add_combined_VM16S_assays(
    mae,
    assay1 = "VM16S_S2017_ASVseq_tax_glom",
    assay2 = "VM16S_UMD_ASV_ASVseq_tax_glom",
    new_assay_name = "VM16S_combined"
    )
```

###### 2.5.3 16S rRNA: proportions

Because size effects are difficult to estimate, ASV or species proportions are usually not a reliable indicator of true variations in counts. However, it is still useful to compute the ASV/species proportion to analyze the share of Lactobacillus species in the microbiota. Consequently, for each 16S rRNA assay, we compute the proportion of each ASV per sample.

```
assay_names = c("VM16S_combined","VM16S_VMRC_ASVseq_tax_glom", "VM16S_UMD_OTU")

for(a in assay_names){
  A = assay(mae, a)
  rA = t(t(A)/colSums(A))
  new_assay = list()
  new_assay[[str_c(a,"_p")]] = rA
  mae = c(mae, new_assay, mapFrom = match(a, names(experiments(mae))))
}

saveRDS(mae, file = "../results/mae_2_data_transformation_16S.Rds")
```

###### 2.5.4 Proportion of Lactobacillus and non-Lactobacillus species in each sample

```
prop_Lacto_combined <- identify_Lacto_prop(mae, assay_name = "VM16S_combined_p")
prop_Lacto_OTU <- identify_Lacto_prop(mae, assay_name = "VM16S_UMD_OTU_p")
```

```r
prop_Lacto <-
  bind_rows(
    prop_Lacto_combined,
    prop_Lacto_OTU %>% filter(!(SampleID %in% prop_Lacto_combined$SampleID))
    )

m <- match(colData(mae)$SampleID, prop_Lacto$SampleID)
# (colData(mae)$SampleID == prop_CST4$SampleID[m]) %>% all(., na.rm = TRUE)
colData(mae)$prop_Lacto = prop_Lacto$prop_Lacto[m]
colData(mae)$prop_nonLacto = prop_Lacto$prop_CST4[m]

saveRDS(mae, file = "../results/mae_3_prop_Lacto_nonLacto.Rds")
```

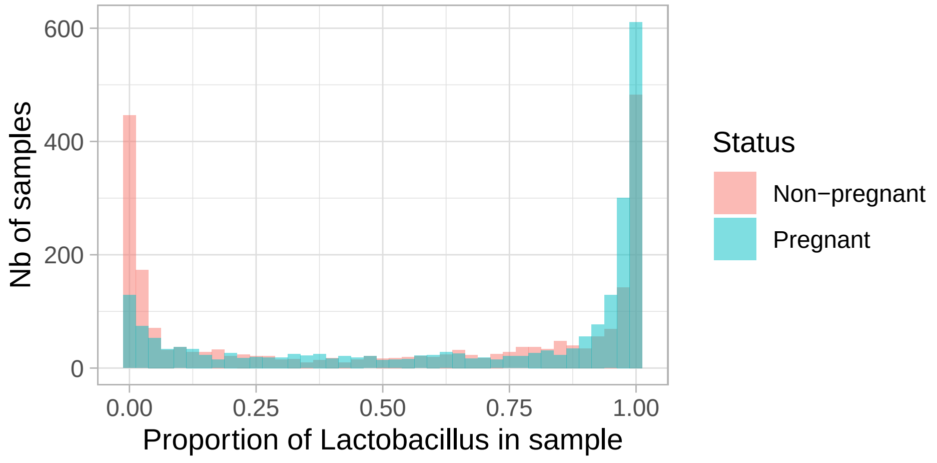

Figure 8: Distribution of the proportion of Lactobacillus in samples from Pregnant and Non-pregnant participants.

```r
ggplot(colData(mae) %>%
         as.data.frame() %>%
         filter(Status == "Non-pregnant") %>%
         select(Subject, Timepoint_days, Bleeding, prop_Lacto, prop_nonLacto) %>%
         pivot_longer(cols = starts_with("prop_") ,
                      names_to = "ASV_cat",
                      values_to = "prop") %>%
         mutate(ASV_cat =
                  ifelse(ASV_cat == "prop_Lacto","Lacto","non-Lacto") %>%
                  factor(., levels = c("non-Lacto","Lacto"))
         ),
       aes(x = Timepoint_days)) +
  geom_bar(aes(y = prop, fill = ASV_cat), stat = "identity") +
  geom_point(aes(y = -0.1, col = Bleeding), size = 0.5) +
  scale_color_gradient(low = "transparent", high = "red") +
  scale_fill_discrete("") +
  facet_wrap(Subject ~ .) +
  ylab("Nb of samples") +
  theme(legend.position = "bottom")
```

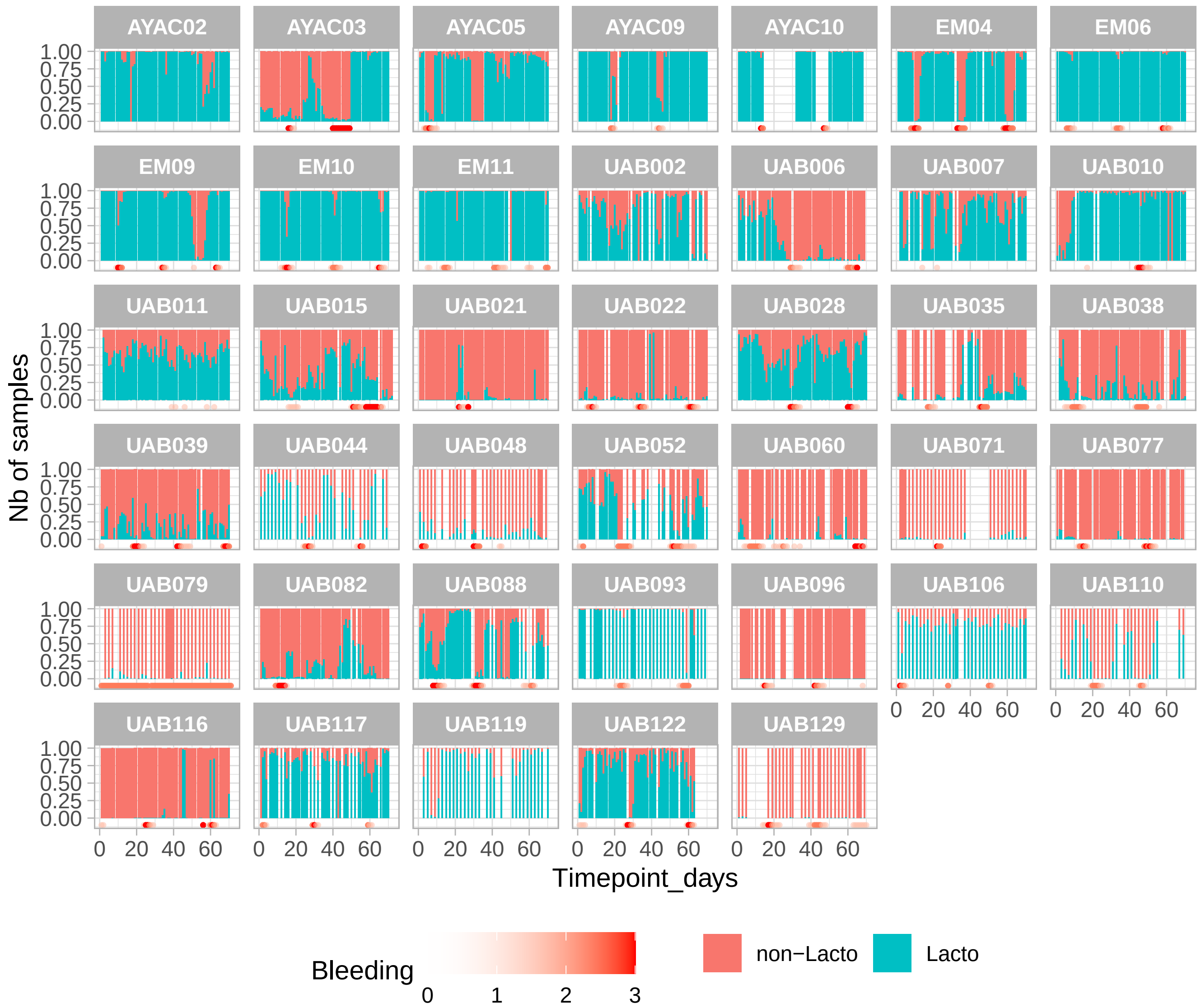

Figure 9: Time-series of the proportion of Lactobacillus species in non-pregnant subjects.

##### 2.5.5 Metabolites: transformation and imputation

Here, we first transform the metabolites data with a variance-stabilizing method and remove effects from metabolomics batches in P women samples. We then check the distribution of missing data, and impute the missing data with a KNN-imputation.

**Transformation**

```
# MB_P = assay(mae, "MB_P")
# MB_NP = assay(mae, "MB_NP")
#
# MB_NP_t_vsn = MB_NP %>% as.matrix() %>% justvsn() %>% as.matrix()
# MB_P_t_vsn = MB_P %>% as.matrix() %>% justvsn() %>% as.matrix()

MB_NP_t <- transform_and_correct_MB_per_batch(mae, "MB_NP")
MB_P_t <- transform_and_correct_MB_per_batch(mae, "MB_P")

mae <- c(mae, MB_P_t = MB_P_t, mapFrom = match("MB_P", experiments(mae) %>% names()))
mae <- c(mae, MB_NP_t = MB_NP_t, mapFrom = match("MB_NP", experiments(mae) %>% names()))
```

**Missing data**

```
# scatter plot of missing frequency vs metabolite mean abundance
ggplot(
  rbind(
    tibble(Status = "Pregnant", mean = rowMeans(MB_P_t, na.rm = TRUE), f_missing = rowMeans(is.na(MB_P_t))),
    tibble(Status = "Non-pregnant", mean = rowMeans(MB_NP_t, na.rm = TRUE), f_missing = rowMeans(is.na(MB_NP_
  ),
  aes(x = 100* f_missing, y = mean, col = Status)
) +
  geom_point() +
  xlab("% of samples in which a metabolite is missing") +
  ylab("Mean abundance of metabolites") +
  ggtitle("The frequency of missing data is correlated with the mean abundance",
          sub = "each dot is a metabolite")
```

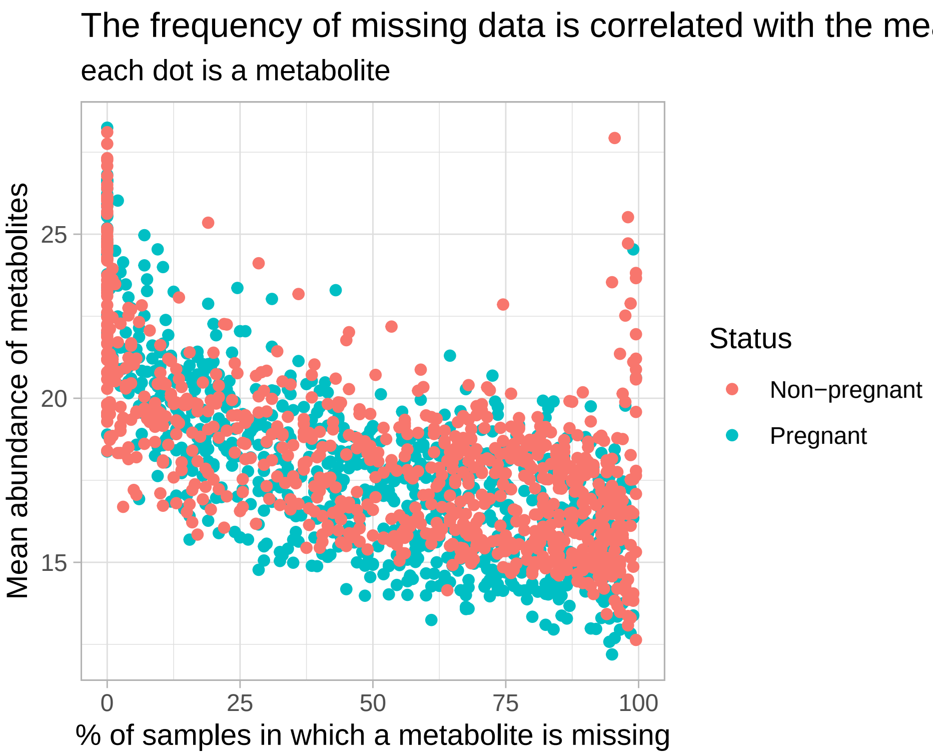

Figure 10: The frequency of missing data is higher for metabolites with lower abundances.

```
# histogram of the number of missing metabolite per sample

# ggplot(
#   rbind(
#     tibble(Status = "Pregnant", mean = colMeans(MB_P_t, na.rm = TRUE), f_missing = colMeans(is.na(MB_P_t)))
#     tibble(Status = "Non-pregnant", mean = colMeans(MB_NP_t, na.rm = TRUE), f_missing = colMeans(is.na(MB_N
#   ),
#   aes(x = 100* f_missing, y = mean, col = Status)
# ) +
#   geom_point() +
#   xlab("% of missing metabolites in sample") +
#   ylab("Mean abundance of metabolites in sample")
```

###### Imputation

Metabolon communicates that missing data indicate concentration below LLOQ. However, based on how metabolite concentrations are measured, there are good reasons to believe that technical artifacts might also generate missing data, including when concentrations were not low. For example, missing data could be due to mis-alignment of the m/z peaks or mis-calibration of the background noise level. Without access to the raw data, it is difficult to diagnose potential issues. However, we can make a few reasonable assumptions and impute for a subset of the data.

Since there are a lot of missing data, we only impute data for metabolites that are missing less than 50% of the time. And since there are a lot of samples which still have a high proportion of missing metabolites after filtering out those that are missing frequently, we only impute for samples that have less than 60% of missing metabolites.

```
MB_P_t_for_imputation <- MB_P_t[(rowMeans(is.na(MB_P_t)) < 0.5 ) %>% as.vector() %>% which(), ]
MB_P_t_for_imputation <- MB_P_t_for_imputation[, colMeans(is.na(MB_P_t_for_imputation)) < 0.6]
MB_NP_t_for_imputation <- MB_NP_t[(rowMeans(is.na(MB_NP_t)) < 0.5 ) %>% as.vector() %>% which(), ]
MB_NP_t_for_imputation <- MB_NP_t_for_imputation[, colMeans(is.na(MB_NP_t_for_imputation)) < 0.6]

dim(MB_P_t_for_imputation)
```

```
## [1] 390 179
```

```
dim(MB_NP_t_for_imputation)
```

```
## [1] 336 199
```

To impute metabolites, we make the following assumptions: - when the number of missing metabolites in a sample is low, this indicates a high-quality sample (the alignment of the peaks and the calibration of the background noise levels went fine). In that case, a missing value likely reflect a low concentration of that metabolite. It would thus makes sense to impute that missing metabolite to a fraction (e.g. 90%) of the lowest values measured for that metabolite. - on the other side of the spectrum, when the number of missing metabolites in a sample is high, this likely indicate that something went "wrong" in the peaks detection or background calibration. In that case, a missing value should rather be considered as a technical failure and the values of these metabolites would be better imputed with a KNN imputation.

Consequently, missing metabolites concentration are imputed as a mixture between a low concentration and a KNN imputed value based on the quality of the sample.

To support this imputation strategy, we show the time-series of the concentrations of *lactate* in subjects with missing values of *lactate*. *Lactate* is a metabolite which is rarely missing and its concentration is correlated with the proportion of *Lactobacillus.*

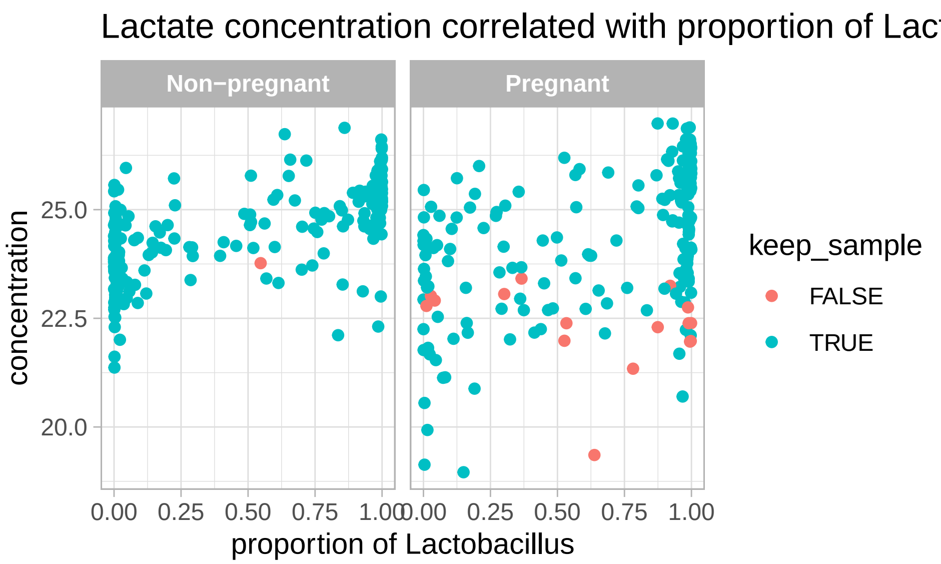

Figure 11: Correlation between lactate concentrations and proportion of Lactobacillus

Now, if we look at the time-series of subjects with missing data points for *lactate*, we see that *lactate* is often missing when the sample quality is low, not always when the proportion of *Lactobacillus* is low. In fact, all samples in which *lactate* is missing are excluded for imputation.

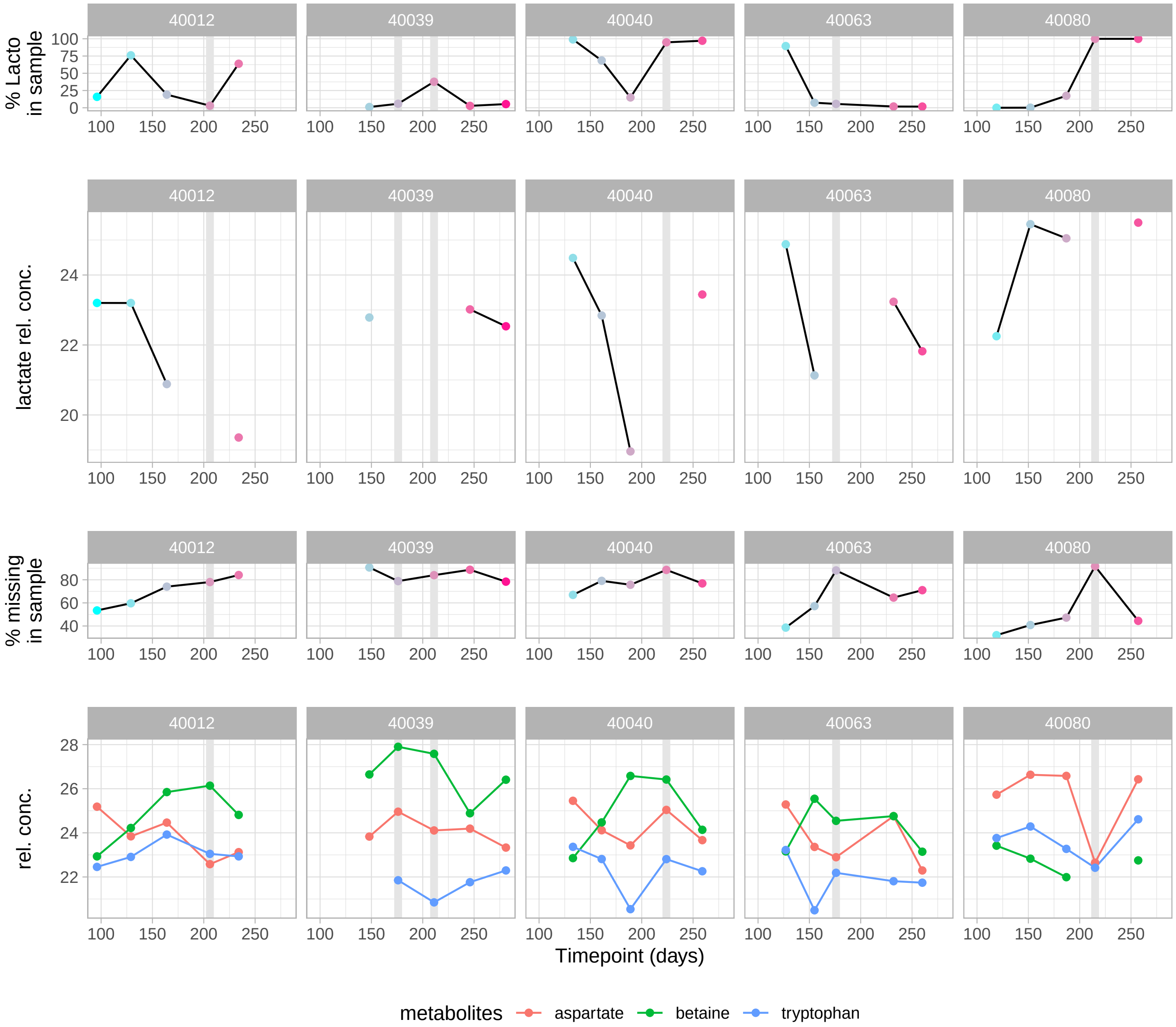

Figure 12: Time-series of subjects with missing data for lactate.

Finally, before imputing, we also check that the sample quality is not correlated with the proportion of *Lactobacillus* (our main biological interest)

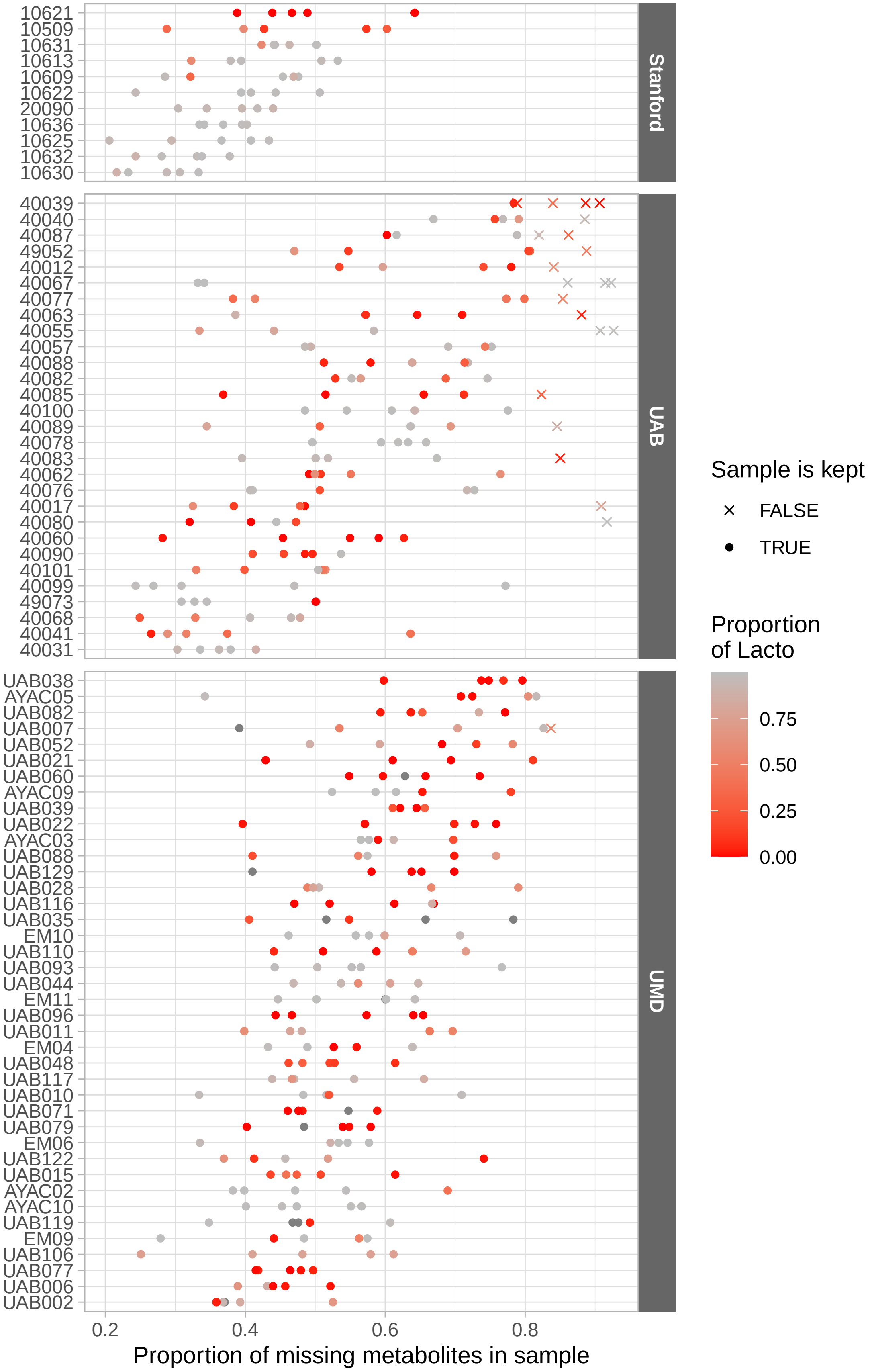

Figure 13: Metabolites samples quality in each cohort.

It looks like samples from Stanford are overall of better quality and samples from UAB has most missing metabolites. It also looks like the sample quality depends on individuals (some participants have overall better sample quality). However, within each cohort, there is no gradient based on the proportion of *Lactobacillus*.

Overall, these observations support (or do not oppose) the proposed imputation strategy.

The KNN imputation goes as follow:

- The K nearest neighbors of each *sample* (and not *metabolites*) are found by computing the euclidean distance between the metabolite abundance of non-missing metabolites in both samples.
- The value of missing metabolites for a given sample is the weighted mean of the values for this metabolites in the K nearest neighbors for which that metabolite is not missing. The weight is inversely proportion to the distance between that samples and the neighbors.

The reason why values are imputed by looking at the NN of each sample and not each metabolite is because the variance in metabolite abundance is much larger across metabolites than across samples. In other words, if a given metabolite $m$ is missing in a given sample $s$, its value is more likely to be close to the mean value of non-missing metabolites $m$ in different samples than to the mean value of the non-missing metabolites (different than $m$) of that sample $s$.

```
MB_P_t_KNN_imputed <- impute_knn(data = MB_P_t_for_imputation, k = 5)
MB_NP_t_KNN_imputed <- impute_knn(data = MB_NP_t_for_imputation, k = 5)

MB_P_t_LLOQ_imputed <- impute_lloq(data = MB_P_t_for_imputation, f = 0.95)
MB_NP_t_LLOQ_imputed <- impute_lloq(data = MB_NP_t_for_imputation, f = 0.95)

P_sample_quality <-  colMeans(MB_P_t_for_imputation %>% is.na())
P_sample_quality <- 1 - P_sample_quality/max(P_sample_quality)

NP_sample_quality <-  colMeans(MB_NP_t_for_imputation %>% is.na())
NP_sample_quality <-  1 - NP_sample_quality/max(NP_sample_quality)

MB_P_t_imputed <-
  t(t(MB_P_t_KNN_imputed) * (1-P_sample_quality) +
      t(MB_P_t_LLOQ_imputed) * P_sample_quality )

MB_NP_t_imputed <-
  t(t(MB_NP_t_KNN_imputed) * (1-NP_sample_quality) +
      t(MB_NP_t_LLOQ_imputed) * NP_sample_quality )
```

```
P_sample_map <-
  data.frame(
    assay = "MB_P_t_imputed",
    primary = MB_P_t_imputed %>% colnames(),
    colname = MB_P_t_imputed %>% colnames()
  )

mae = c(mae, MB_P_t_imputed = MB_P_t_imputed,
        sampleMap = P_sample_map)

NP_sample_map <-
  data.frame(
    assay = "MB_NP_t_imputed",
    primary = MB_NP_t_imputed %>% colnames(),
    colname = MB_NP_t_imputed %>% colnames()
  )

mae <- c(mae, MB_NP_t_imputed = MB_NP_t_imputed,
        sampleMap = NP_sample_map)
```

##### 2.5.6 Immunology data: transformation and imputation

Here, we will transform, then impute missing data in the immunology assay.

###### Transformation

```
I <- assay(mae, "I")
range(I, na.rm = TRUE)
```

```
## [1] 1.084010e-03 4.294172e+05
```

```
I_t <- log10(I) # asinh(I) is not as good

hist(I_t, breaks = 100, main = "", xlab = "")
```

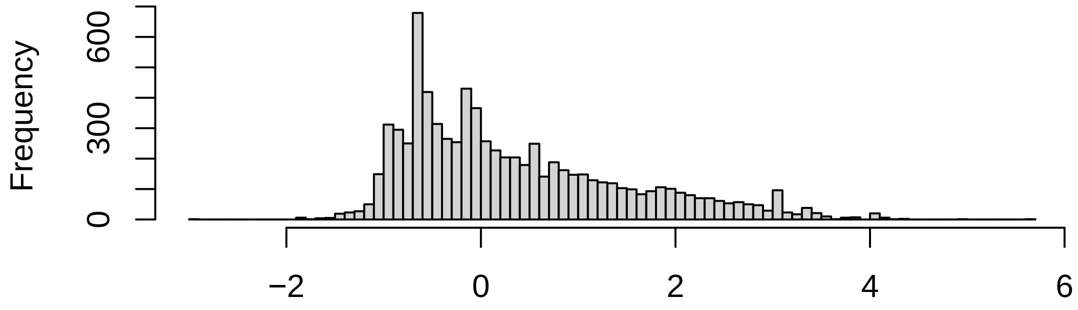

Figure 14: Histogram of transformed cytokine abundances.

```
mae <- c(mae, I_t = I_t, mapFrom = match("I", experiments(mae) %>% names()))
```

###### Cytokines distributions

```
tmp <- I_t %>% t()
o <- order(colMeans(tmp, na.rm = TRUE))
boxplot(tmp[,o], pch = 16, cex = 0.5)
```

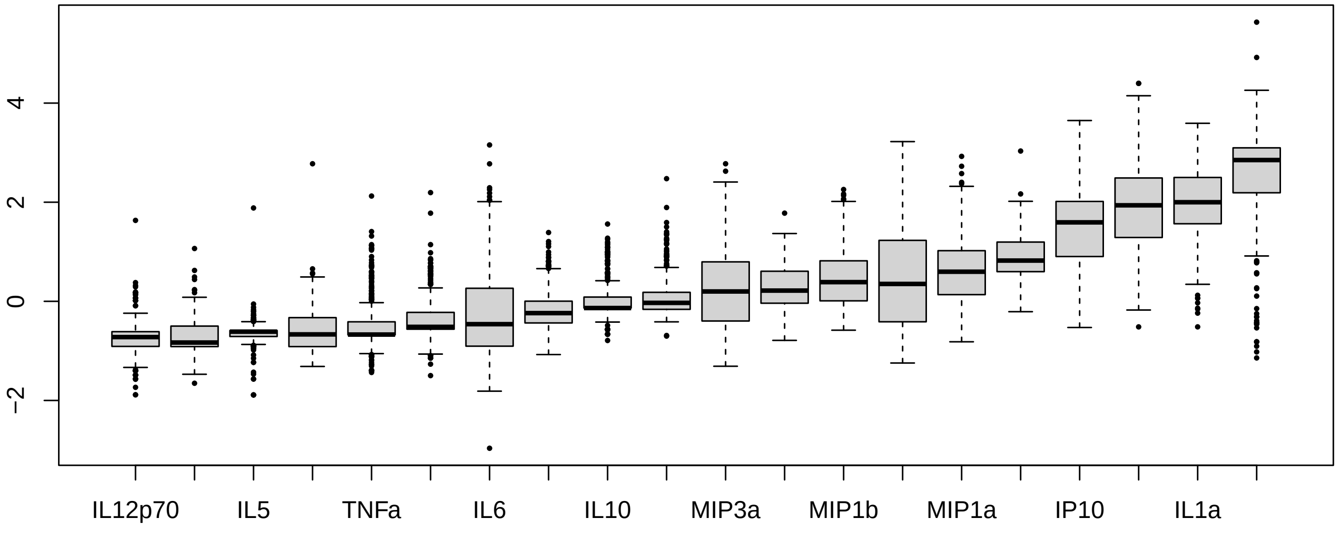

Figure 15: Cytokines distributions

###### Imputation

There are only a few missing data in the immunology data:

```
n_missing_per_sample <- apply(I_t, 2, function(x) sum(is.na(x)))
table(n_missing_per_sample) %>%
```

```r
  as.data.frame() %>%
  set_colnames(c("n_missing_per_sample", "Freq")) %>%
  dplyr::rename(
    `# of missing cytokine` =  n_missing_per_sample,
    `# of sample` = Freq) %>%
  kable(., format = "latex", booktab = TRUE)
```

| # of missing cytokine | # of sample |
|---|---|
| 0 | 381 |
| 1 | 7 |
| 3 | 1 |
| 5 | 1 |

```r
n_missing_per_cytokine <- apply(I_t, 1, function(x) sum(is.na(x)))
table(n_missing_per_cytokine) %>%
  as.data.frame() %>%
  set_colnames(c("n_missing_per_cytokine", "Freq")) %>%
  dplyr::rename(
    `# of missing samples` =  n_missing_per_cytokine,
    `# of cytokines` = Freq) %>%
  kable(., format = "latex", booktab = TRUE)
```

| # of missing samples | # of cytokines |
|---|---|
| 0 | 10 |
| 1 | 6 |
| 2 | 3 |
| 3 | 1 |

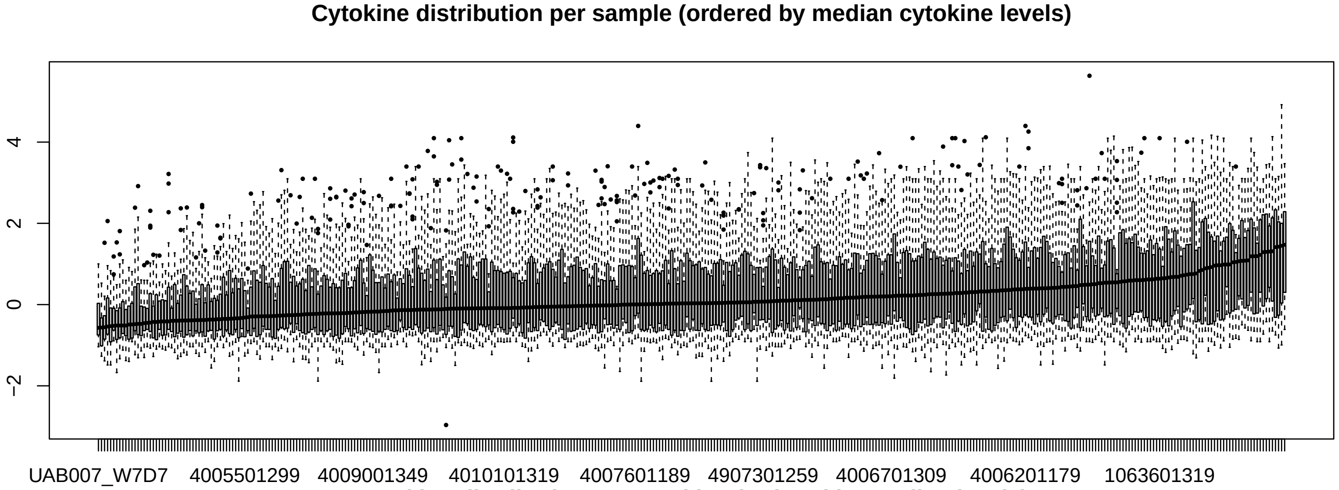

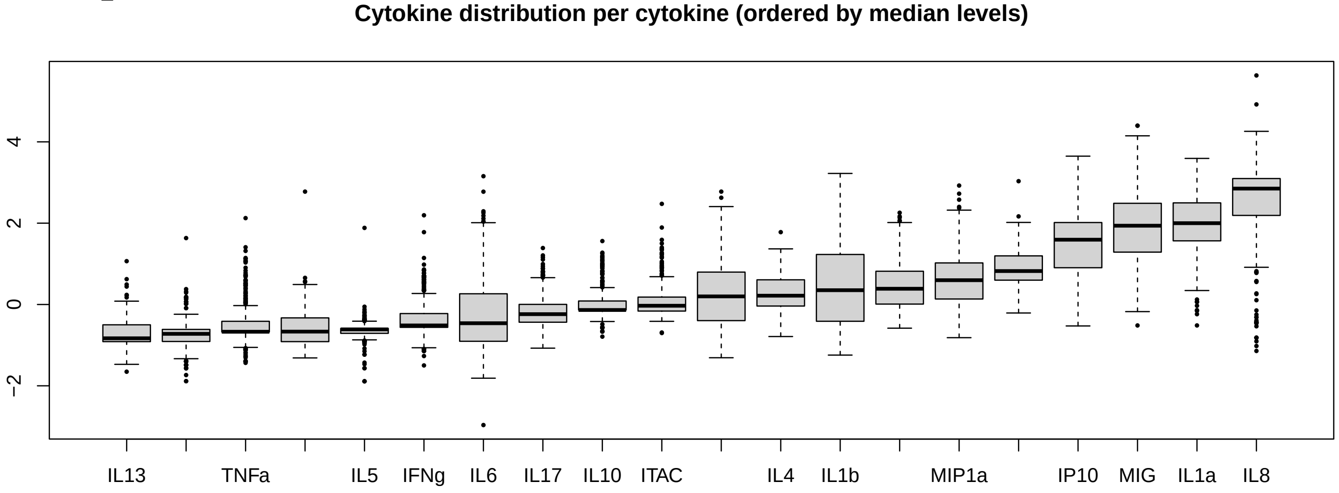

From these distributions, it looks like we are in a case similar to the metabolites although the between-sample variance is larger for the cytokines than for the metabolites.

Consequently, we use the same knn imputation approach to impute the missing data.

```
I_t_imputed <- impute_knn(data = I_t, k = 5)
```

```
mae <- c(mae, I_t_imputed = I_t_imputed,
         mapFrom = match("I", experiments(mae) %>% names()))
```

##### 2.5.7 Save MAE with transformed assays

```
saveRDS(mae, file = "../results/mae_4_transformed_data.Rds")
```

### 3 Data augmentation

#### 3.1 CST and subCST (Valencia) assignment

```
subCSTs <- identify_subCSTs(mae, assay_name = "VM16S_combined")
subCSTs <-
  tibble(SampleID = mae$SampleID) %>%
  left_join(subCSTs, by = "SampleID") %>%
  mutate(
    CST = str_remove(sub_CST, "-.*$")
  )

mae$CST <- subCSTs$CST
mae$subCST <- subCSTs$sub_CST
```

```
saveRDS(mae, file = "../results/mae_6_with_CST.Rds")
```

Table 3: Distribution of CSTs across reproductive status groups

| Status | CST I | CST II | CST III | CST IV | CST V |
|---|---|---|---|---|---|
| Non-pregnant | 204 (13%) | 162 (11%) | 261 (17%) | 857 (56%) | 50 (3%) |
| Pregnant | 612 (28%) | 100 (5%) | 901 (41%) | 464 (21%) | 102 (5%) |

#### 3.2 Menstrual cycles identification from bleeding reports of non-pregnant subjects

##### 3.2.1 Data

Bleeding intensity on a scale from 0 (none) to 3 (heavy) was reported by non-pregnant subject at each sample collection. This data is available in the sample information of the mae object.

We load the mae object and display the bleeding data for each subject.

```
mae <- readRDS("../results/mae_6_with_CST.Rds")
```

```
si <- mae@colData %>%  as.data.frame()

plot_bleeding_patterns_all_subjects(
  df = si %>% filter(Cohort == "UMD") %>% select(Subject, Timepoint_days, Bleeding))
```

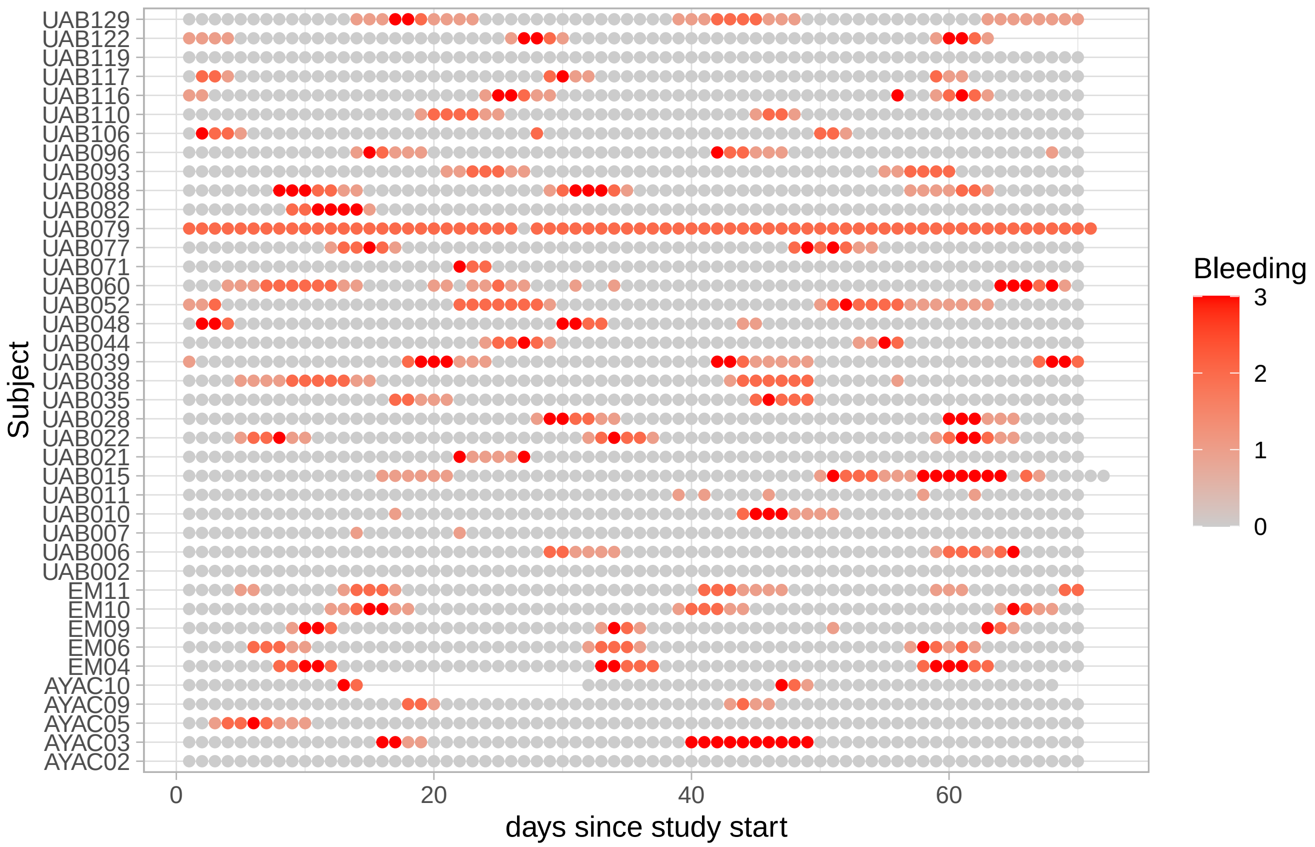

Figure 16: Bleeding intensities reported by non-pregnant subjects.

##### 3.2.2 Menstrual cycle HSMM

To identify menstrual cycle phases, we model the menstrual cycle with hidden semi-Markov model and decode the bleeding time-series to identify menses and then count forward and backward from the first day of the menses.

**Model specification**

To model the menstrual cycle, we define a 3-state model. The first state is the menses, the 2nd state is the cycle and the 3rd state is the pre-menstrual phase.

Bleeding is very likely and likely medium-heavy in the menses state, rather unlikely in the cycle state and spotting is possible in the pre-menstrual phase

```
library(HiddenSemiMarkov)
```

```r
M <- max(si$Timepoint_days[si$Cohort == "UMD"], na.rm = TRUE)
# typical cycle length (28-29) - typical period length (4-5) - pre-menstrual phase (3)
cycle_sojourn <- dgamma(1:M, shape = 18.5, scale = 1.3)
cycle_sojourn[1:16] <- 0 # we prevent cycles that are too short
menses_sojourn <- dpois(1:(M-1), lambda = 3)
menses_sojourn <- c(0,menses_sojourn) # we shift by 1
menses_sojourn[8:M] <- 0
pre_menses_sojourn <- rep(0, M)
pre_menses_sojourn[3] <- 1

M_hsmm <- specify_hsmm(
  J = 3,
  state_names = c("cycle","pre-menses","menses"),
  state_colors = c("gray80","goldenrod","red"),
  init = rep(1/3,3),
  transition = matrix(c(0,1,0,0,0,1,1,0,0),3,3, byrow = TRUE),
  sojourn =
    list(type = "ksmoothed_nonparametric",
         d = cbind(cycle_sojourn, pre_menses_sojourn, menses_sojourn),
         bw = 2
    ),
  marg_em_probs =
    list(
      Bleeding = list(
        type = "non-par",
        params = list(
          values = c(0,1,2,3) %>% as.character(),
          probs = cbind(c(0.9,0.09,0.009, 0.001), c(0.7,0.29,0.009, 0.001), c(0.2,0.3,0.25, 0.25))
        )
      )
    ),
  censoring_probs = list(p = c(0.2,0.05,0.05), q = matrix(0,nrow = 1,ncol = 3))
)
```

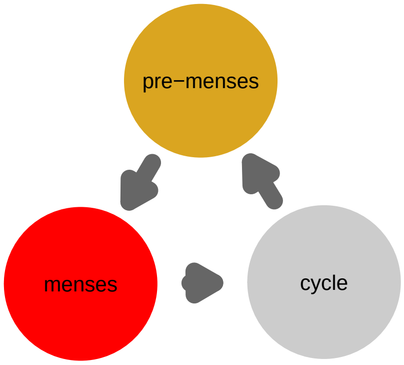

Figure 17: Graph of the hidden semi-Markov model (HSMM) used to detect menstrual cycles from bleeding records.

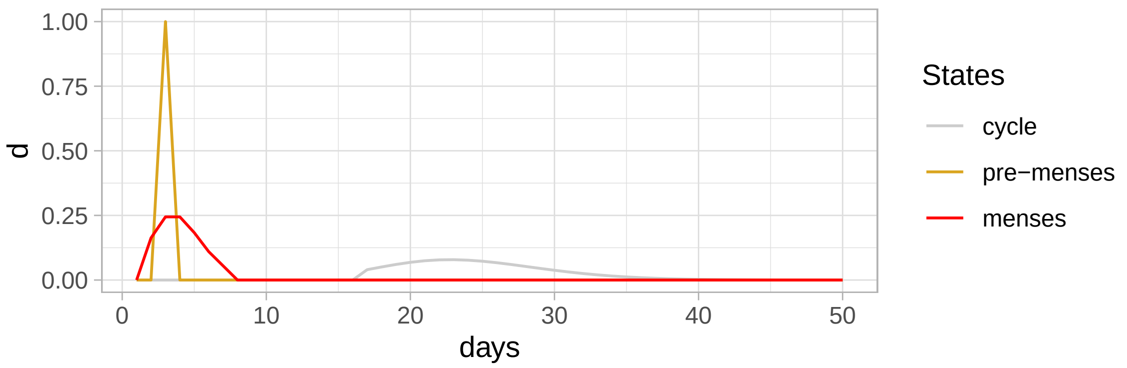

Figure 18: Specified sojourn distribution for our menstrual cycle HSMM

##### 3.2.3 Pre-processing bleeding data

To be able to use the `HiddenSemiMarkov` package functions, we need to make sure that there is data at each time-step.

```
df <-
  si %>%
  filter(!is.na(Bleeding)) %>%
  filter(Cohort == "UMD") %>%  # we select only the cohort of non-pregnant women
  # we also need to ensure that menstrual cycles can be identified by
  # removing subject that never reported any bleeding or
  # that reported constant bleeding

  group_by(Subject) %>%
  mutate(has_some_bleeding = ( sum(Bleeding) > 0),
         is_constantly_bleeding = mean(Bleeding > 0) > 0.5
  ) %>%
  ungroup() %>%
  filter(has_some_bleeding, !is_constantly_bleeding) %>%
  select(Subject, Timepoint_days, Bleeding)

X <- prepare_bleeding_data_for_HSMM(df)
```

##### 3.2.4 Fitting the model to Subjects' time series

```
fit_output <- fit_hsmm(model = M_hsmm, X = X, rel_tol = 1/100000, n_iter = 100, N0_sojourn = 20)
plot_hsmm_fit_status(fit_output)
```

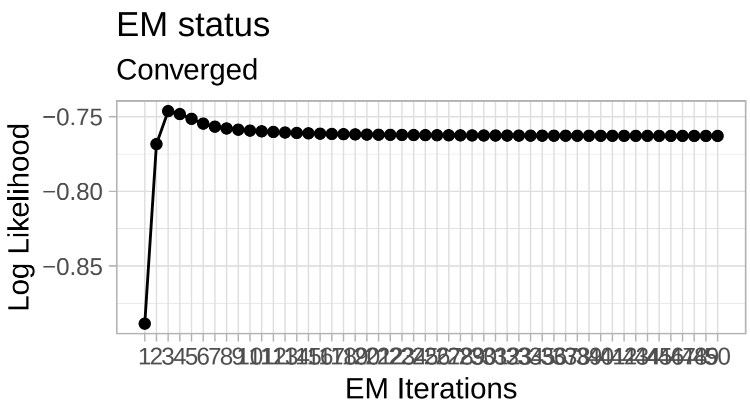

Figure 19: Fitting the HSMM: likelihood over the EM steps.

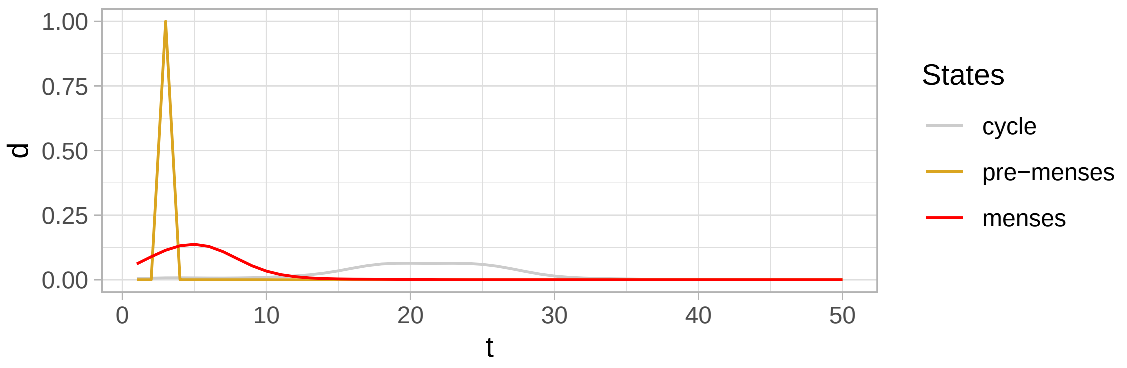

Figure 20: Sojourn distributions of the fitted HSMM.

##### 3.2.5 Decoding Subjects' time series to identify menses

```
Y <- decode_subjects_time_series(X = X, model = M_hsmm)
Y_fitted <- decode_subjects_time_series(X = X, model = fit_output$model)
```

##### 3.2.6 Determining cycle number, forward and backward cycledays and cycle length

```
C <- compute_cycleday(Y = Y)
C_fitted <- compute_cycleday(Y = Y_fitted)
```

##### 3.2.7 Choosing specified vs fitted model results

```
plot_bleeding_patterns_all_subjects(df) +
  geom_point(data = C %>% filter(is_menses), aes(x = t, y = seq_id), col = "blue", shape = 1) +
  ggtitle("Identified menses with specified model")
```

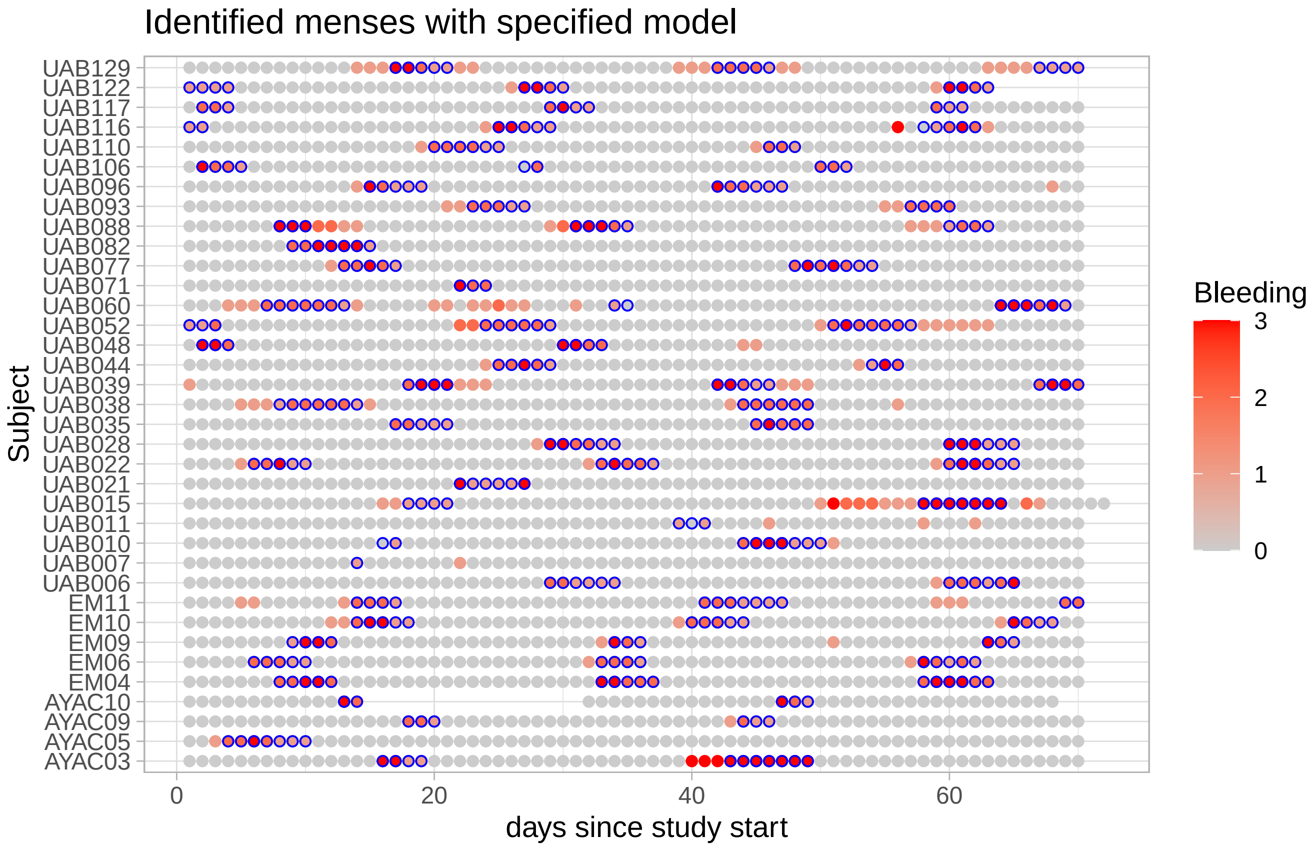

```r
plot_bleeding_patterns_all_subjects(df) +
  geom_point(data = C_fitted %>% filter(is_menses), aes(x = t, y = seq_id), col = "blue", shape = 1) +
  ggtitle("Identified menses with fitted model")
```

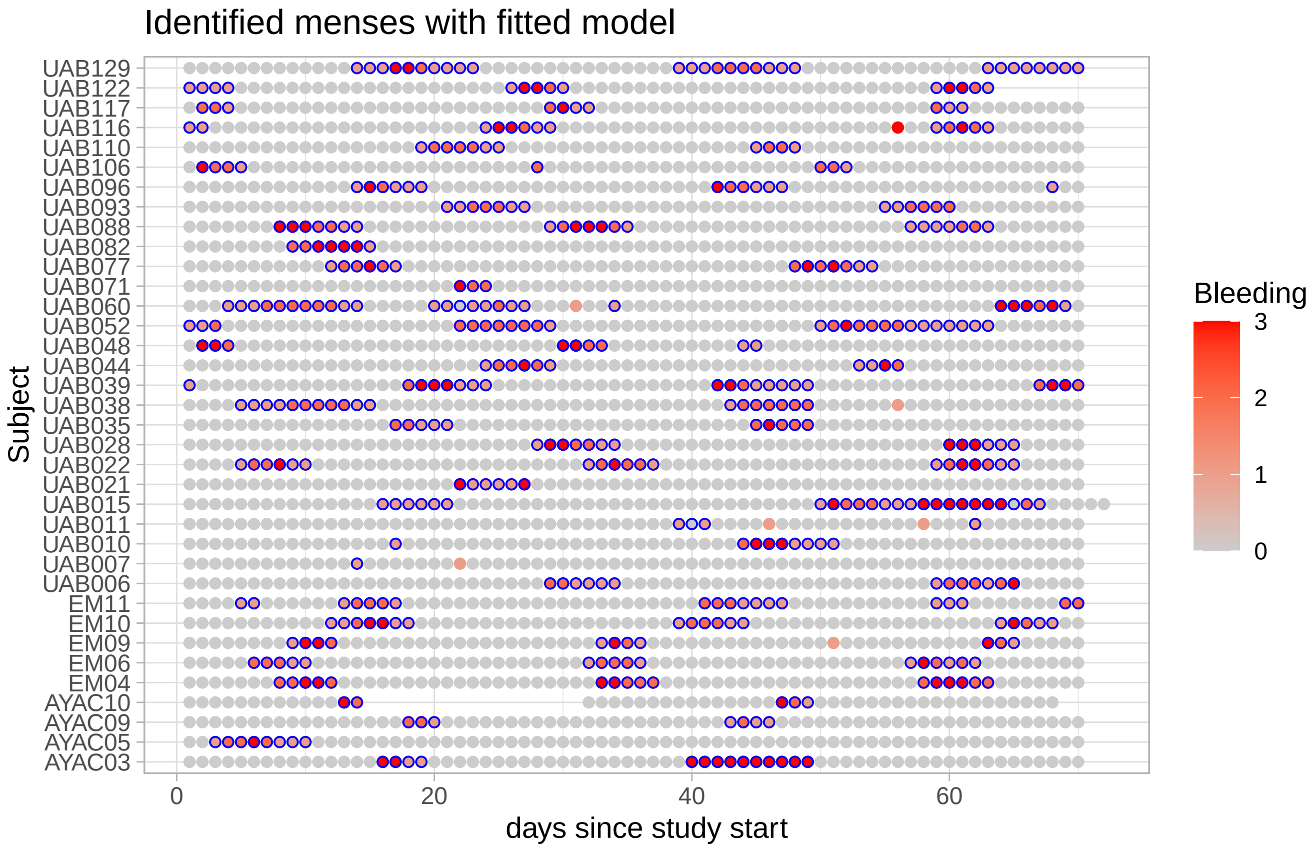

##### 3.2.8 Adding menstrual cycle info and characteristics to the MAE object

Because the decoding appears better with the specified model, we keep the menstrual cycle info from the specified model decoding. We also add a column MC_ok to flag samples for which the cycleday could be reliably identified. This excludes cycles identified from Subjects who reported less than 3 days or more than 30 days of bleeding within the 10 weeks of the study, cycles with periods shorter than two days or which started or finished too late or too early within the standardized window.

```
mae <- add_MC_data_to_mae(mae, si, C)
```

```
new_colData <- colData(mae) %>% as.data.frame()

g_cycles <-
  ggplot(new_colData %>% filter(Status == "Non-pregnant"),
       aes(x = Timepoint_days, y = Subject)) +
  geom_point(aes(col = Bleeding)) +
  scale_color_gradient(low = "gray80", high = "red3") +
  geom_point(data = new_colData %>% filter(Status == "Non-pregnant", is_menses),
             col = "blue", shape = 16, size = 0.2) +
  geom_point(data = new_colData %>% filter(Status == "Non-pregnant", !MC_ok),
             shape = 4, col = "gray20")

save(g_cycles, file = "../results/suppl_figs/cycles.Rdata")

g_cycles
```

Table 4: Number of subjects with 0, 1, 2 or 3 full standardized menstrual cycles identified during the 10-week data collection period.

| has ASV data | 0 M.C. | 1 M.C. | 2 M.C. | 3 M.C. |
|---|---|---|---|---|
| no | 6 | 1 | 3 | 5 |
| yes | 4 | 5 | 13 | 8 |

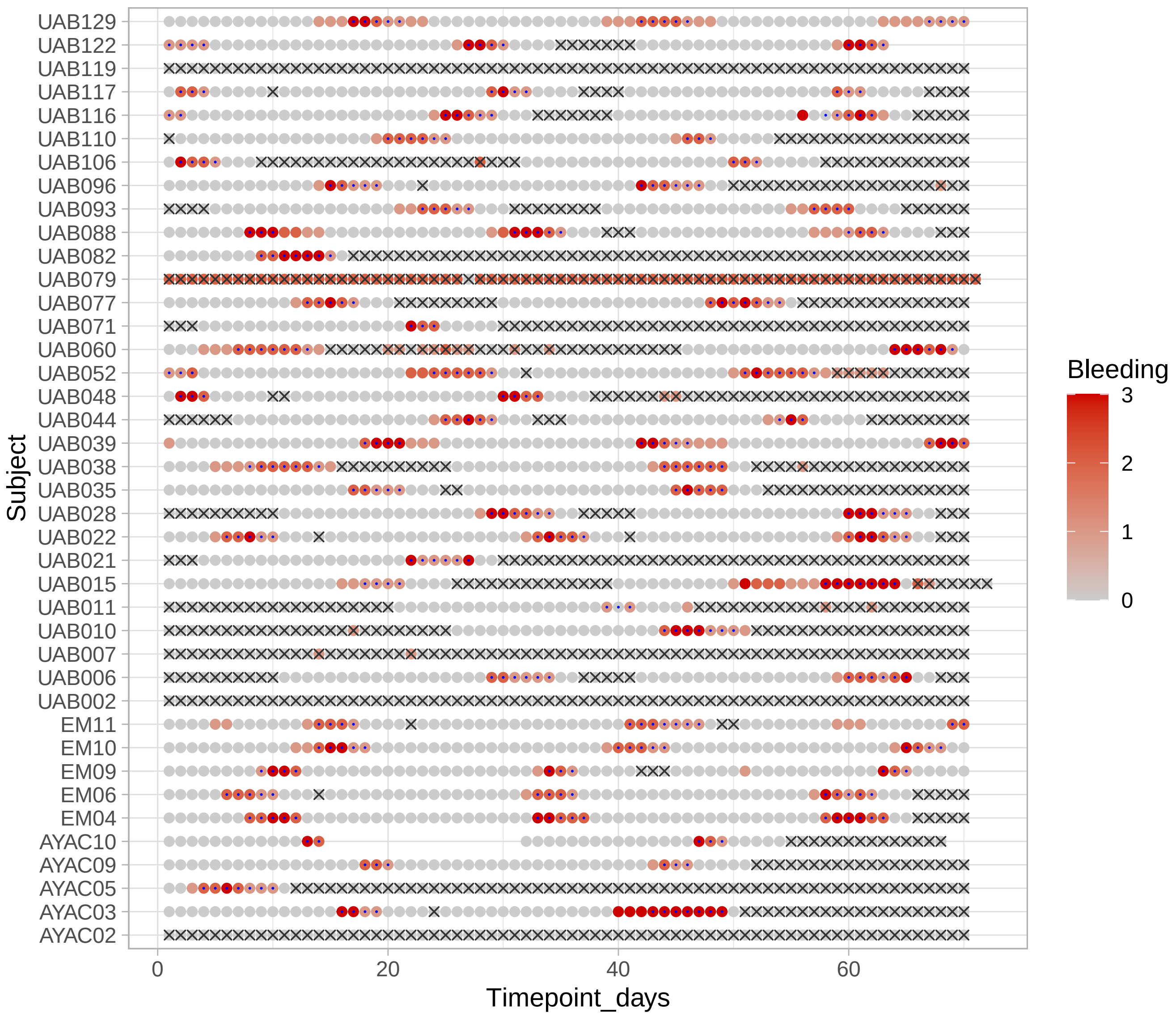

Figure 21: Identified and included standardized menstrual cycles from bleeding data.

#### 3.3 PH and Nugent score throughout the menstrual cycle

```
si <-
  colData(mae) %>%
  as.data.frame() %>%
  dplyr::filter(Status == "Non-pregnant") %>%
  mutate(dominant =
           ifelse((prop_Lacto >= 0.5),"Lactobacillus","non-Lactobacillus") %>%
           factor(., levels = c("non-Lactobacillus","Lactobacillus")))
```

**PH throughout the menstrual cycle**

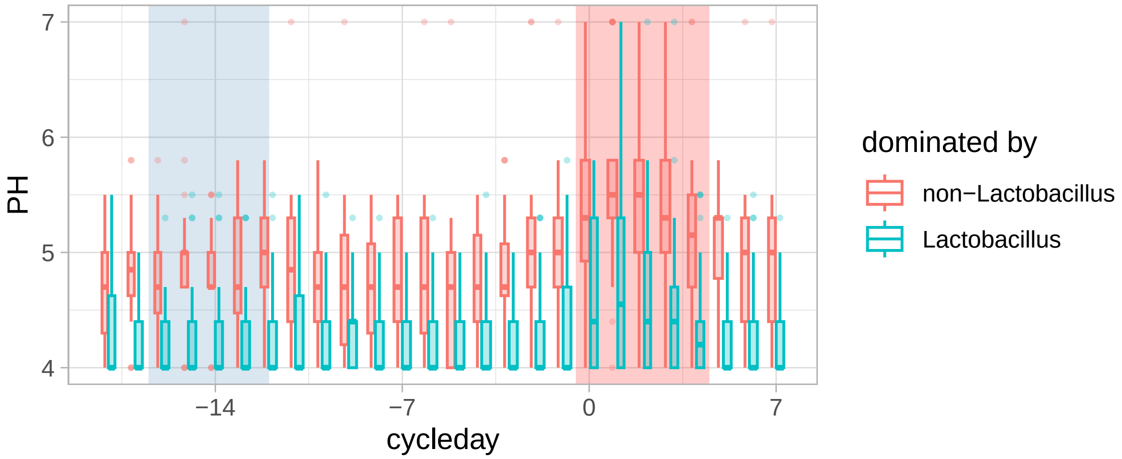

Figure 22: pH throughout the menstrual cycle (boxplot)

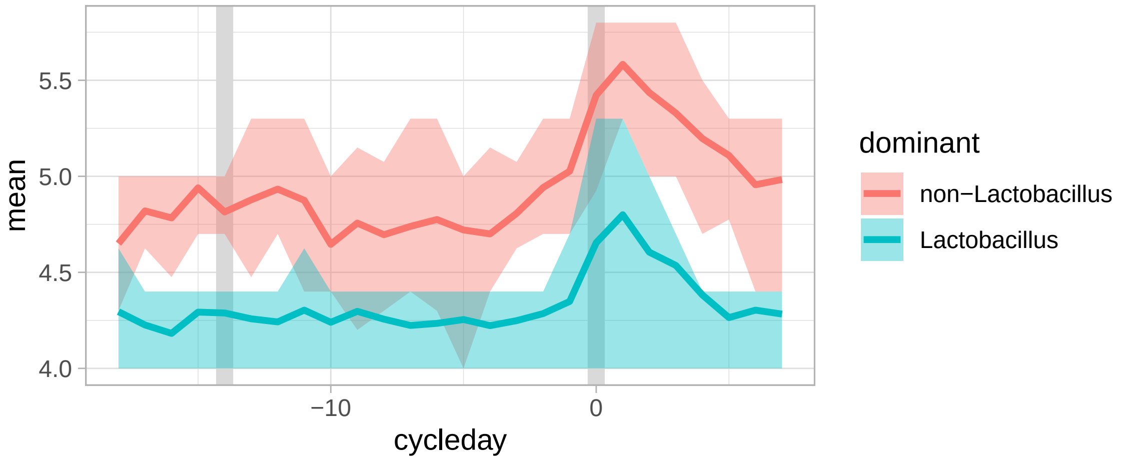

Figure 23: pH throughout the menstrual cycle (ribbon)

```
si %>%
  filter(!is.na(dominant), !is.na(is_menses)) %>%
  group_by(dominant) %>%
  summarize(
    mean_pH = mean(PH, na.rm = TRUE),
    median_pH = median(PH, na.rm = TRUE),
    pc_05_pH = quantile(PH, p = 0.05, na.rm = TRUE),
    pc_95_pH = quantile(PH, p = 0.95, na.rm = TRUE),
    .groups = "drop"
  ) %>%
  kableExtra::kable(
```

Table 5: pH values in samples dominated by Lactobacillus and non-Lactobacillus.

| dominant | mean_pH | median_pH | pc_05_pH | pc_95_pH |
|---|---|---|---|---|
| non-Lactobacillus | 4.967722 | 5 | 4 | 5.8 |
| Lactobacillus | 4.356168 | 4 | 4 | 5.3 |

Table 6: pH values in samples dominated by Lactobacillus and non-Lactobacillus throughout the cycle and during menses

| dominant | is_menses | mean_pH | median_pH | pc_05_pH | pc_95_pH |
|---|---|---|---|---|---|
| non-Lactobacillus | FALSE | 4.859949 | 5.0 | 4.0 | 5.5 |
| non-Lactobacillus | TRUE | 5.415263 | 5.3 | 4.4 | 7.0 |
| Lactobacillus | FALSE | 4.329785 | 4.0 | 4.0 | 5.3 |
| Lactobacillus | TRUE | 4.633333 | 4.4 | 4.0 | 5.8 |

```
    ., format = "latex", booktab = TRUE, linesep = "",
    caption = "pH values in samples dominated by Lactobacillus and non-Lactobacillus.")
```

```
si %>%
  filter(!is.na(dominant), !is.na(is_menses)) %>%
  group_by(dominant, is_menses) %>%
  summarize(
    mean_pH = mean(PH, na.rm = TRUE),
    median_pH = median(PH, na.rm = TRUE),
    pc_05_pH = quantile(PH, p = 0.05, na.rm = TRUE),
    pc_95_pH = quantile(PH, p = 0.95, na.rm = TRUE),
    .groups = "drop"
  ) %>%
  kableExtra::kable(
    ., format = "latex", booktab = TRUE, linesep = "",
    caption = "pH values in samples dominated by Lactobacillus
    and non-Lactobacillus throughout the cycle and during menses")
```

```
si %>%
  filter(!is.na(dominant), !is.na(is_menses)) %>%
  mutate(
    cycle_phase =
      case_when(
        is_menses ~ "menses",
        cycleday < -14 ~ "follicular",
        cycleday %in% -14:-12 ~ "ovulation",
        cycleday < -7 ~ "early-luteal",
        cycleday > 0 ~ "follicular",
        TRUE ~ "late-luteal"
      ),
    cycle_phase =
      cycle_phase %>%
      factor(., levels = c("follicular", "ovulation", "early-luteal", "late-luteal", "menses"))
  ) %>%
  group_by(dominant, cycle_phase) %>%
  summarize(
    mean_pH = mean(PH, na.rm = TRUE),
    median_pH = median(PH, na.rm = TRUE),
    pc_05_pH = quantile(PH, p = 0.05, na.rm = TRUE),
    pc_95_pH = quantile(PH, p = 0.95, na.rm = TRUE),
    .groups = "drop"
```

Table 7: pH values in samples dominated by Lactobacillus and non-Lactobacillus during each phase of the cycle

| dominant | cycle_phase | mean_pH | median_pH | pc_05_pH | pc_95_pH |
|---|---|---|---|---|---|
| non-Lactobacillus | follicular | 4.895395 | 5.0 | 4.0 | 5.635 |
| non-Lactobacillus | ovulation | 4.877612 | 4.7 | 4.0 | 5.500 |
| non-Lactobacillus | early-luteal | 4.743158 | 4.7 | 4.0 | 5.500 |
| non-Lactobacillus | late-luteal | 4.869474 | 5.0 | 4.0 | 5.500 |
| non-Lactobacillus | menses | 5.415263 | 5.3 | 4.4 | 7.000 |
| Lactobacillus | follicular | 4.276018 | 4.0 | 4.0 | 5.300 |
| Lactobacillus | ovulation | 4.261364 | 4.0 | 4.0 | 5.300 |
| Lactobacillus | early-luteal | 4.274193 | 4.0 | 4.0 | 5.000 |
| Lactobacillus | late-luteal | 4.375368 | 4.4 | 4.0 | 5.300 |
| Lactobacillus | menses | 4.633333 | 4.4 | 4.0 | 5.800 |

```
) %>%
kableExtra::kable(
  ., format = "latex", booktab = TRUE, linesep = "",
  caption = "pH values in samples dominated by Lactobacillus
  and non−Lactobacillus during each phase of the cycle")
```

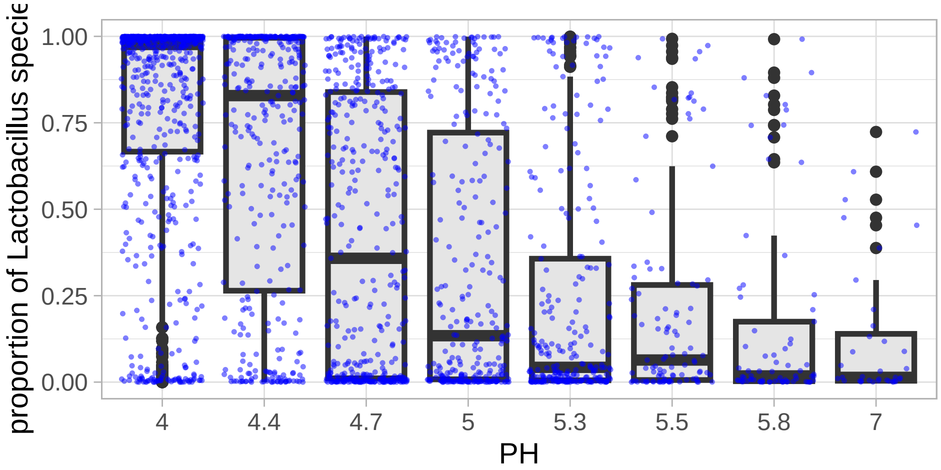

Figure 24: pH and proportion of Lactobacillus

**Nugent score**

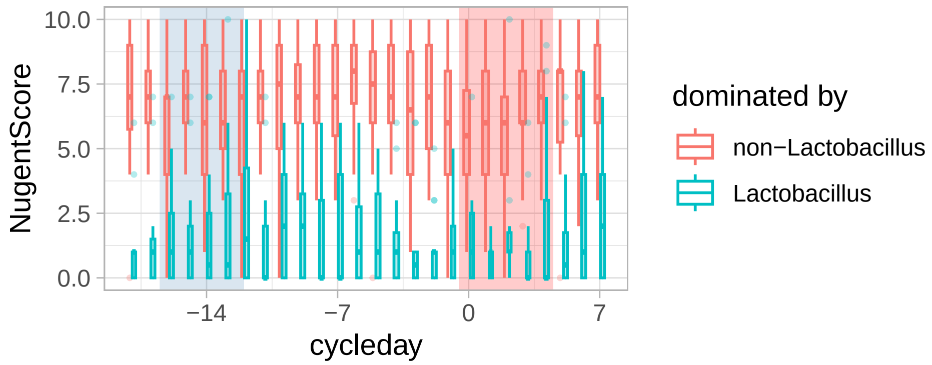

Figure 25: Nugent score throughout the menstrual cycle

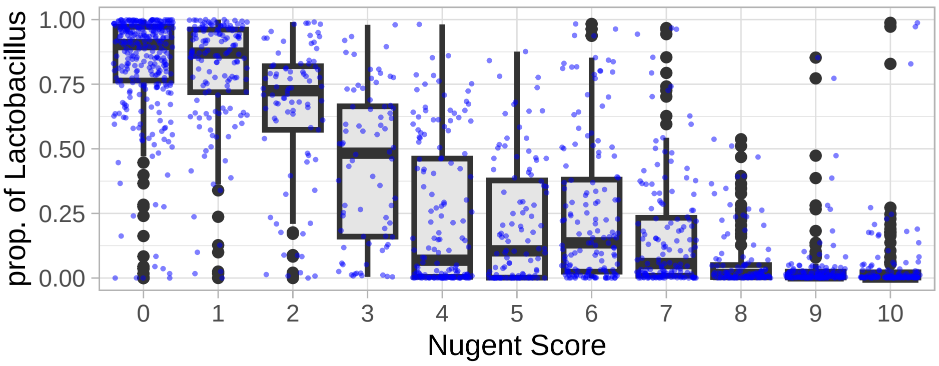

Figure 26: Nugent score and proportion of Lactobacillus

Table 8: Number of sample per reproductive state

| Reproductive status | Nb of samples |
|---|---|
| Pregnant | 2175 |
| Post-partum | 4 |
| Non-pregnant, menses | 368 |
| Non-pregnant, follicular | 630 |
| Non-pregnant, peri-ovulatory | 308 |
| Non-pregnant, luteal | 785 |
| Non-pregnant, undefined | 686 |

#### 3.4 Reproductive status at each sample

In this section, we define several additional categories of reproductive status in addition to the Status "Pregnant"/"Non-Pregnant".

In particular, samples from the pregnant cohort that were collected after delivery get the status "Post-partum". Samples from the non-pregnant cohort are classified according to their menstrual phase (menses, luteal, peri-ovulatory, follicular) or, if no menstrual cycles could be identified or if the menstrual phase could not be defined, they are labelled as "Non-pregnant - undefined".

```
mae <- add_reproductive_status(mae)
saveRDS(mae, file = "../results/mae_7_with_MC.Rds")
```

#### 3.5 Preterm births

```
mae <- readRDS(file = "../results/mae_7_with_MC.Rds")
```

```
si <- colData(mae) %>% as.data.frame()
si %>%
  filter(Status == "Pregnant") %>%
  select(Subject, Site, GestationalAgeAtDelivery_days) %>%
  distinct() %>%
  mutate(Delivery = ifelse(GestationalAgeAtDelivery_days < (37*7), "preterm", "term")) %>%
  group_by(Site, Delivery) %>%
  summarize(n = n()) %>%
  group_by(Site) %>%
  mutate(`%` = (100 * n/sum(n)) %>% round()) %>%
  ungroup() %>%
  kable(., format = "latex", booktab = TRUE,
        caption = "Number (and percentages) of pregnant participants
        who delivered preterm (i.e., before 37 weeks of gestation)") %>%
  kable_styling(latex_options = "HOLD_position")
```

Table 9: Number (and percentages) of pregnant participants who delivered preterm (i.e., before 37 weeks of gestation)

| Site | Delivery | n | % |
|---|---|---|---|
| Stanford | preterm | 9 | 23 |
| Stanford | term | 30 | 77 |
| UAB | preterm | 41 | 43 |
| UAB | term | 55 | 57 |

##### 3.5.1 Proximity to preterm birth score

For each sample of the pregnant subjects, we compute a score that reflect how close in time a sample is to a preterm birth.

```
mae <- add_preterm_proximity_score(mae)
saveRDS(mae, file = "../results/mae_8_with_preterm_risk.Rds")
```

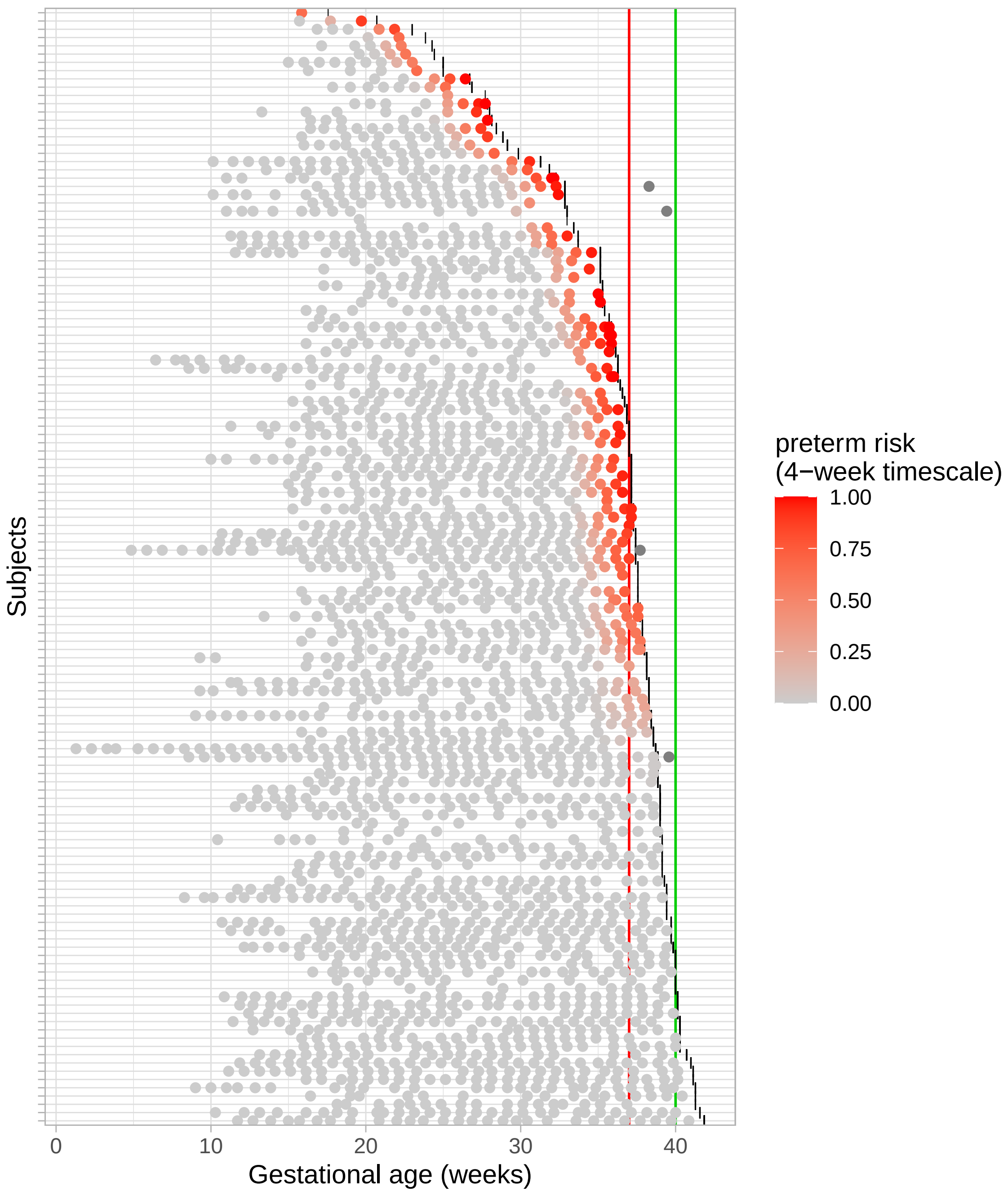

###### MAE for analyses

Data augmentation is done. We copy the latest version of the MAE and rename it to `mae_for_analyses` for convenience.

```
mae = readRDS(file = "../results/mae_8_with_preterm_risk.Rds")
saveRDS(mae, file = "../results/mae_for_analyses.Rds")
```

### 4 Topic analysis

```
LDA_models_and_alignments_dir <- "../results/LDA_models_and_alignments/"
if (!dir.exists(LDA_models_and_alignments_dir))
  dir.create(LDA_models_and_alignments_dir)
```

#### 4.1 Fitting the combined count data with LDA models

LDA models at various resolution (i.e., for K = 2 to K = 25) are fitted to the count data of the pregnant and non-pregnant participants (S2017 and UMD studies).

```
mae <- readRDS("../results/mae_for_analyses.Rds")

count_assay <- "VM16S_combined"
counts <- assay(mae, count_assay) %>% t()
```

```
df <-
  colData(mae) %>%
  as.data.frame() %>%
  select(SampleID, Subject, Status) %>%
  filter(SampleID %in% rownames(counts))

df %>%
  group_by(Status) %>%
  summarize(
    n_samples = n(),
    n_subjects = length(unique(Subject))
    ) %>%
  kable(
    ., format = "latex", booktab = TRUE, linesep = "",
    caption = "Number of samples per reproductive status on which topic models were fitted"
  ) %>%
  kable_styling(latex_options = "HOLD_position")
```

Table 10: Number of samples per reproductive status on which topic models were fitted

| Status | n_samples | n_subjects |
|---|---|---|
| Non-pregnant | 1534 | 30 |
| Pregnant | 2179 | 135 |

```
library(alto)
lda_models <-
  fit_lda_models(
    data = counts,
    dir = str_c(LDA_models_and_alignments_dir, count_assay,"/"),
    max_k = 25
  )
```

We re-label the topics so that their labels match those of the sub-CSTs whose composition is the closest. We use the Bray-Curtis dissimilarity to compare compositions.

```
library(ValenciaR) # devtools::install_github("lasy/ValenciaR")

lda_models_with_name_file <-
  str_c(
```

```
    LDA_models_and_alignments_dir, count_assay,
    "/lda_models_",ncol(lda_models$k1$gamma),
    "_", ncol(lda_models[[length(lda_models)]]$gamma),
    "_named_topics.Rdata"
  )

if (!file.exists(lda_models_with_name_file)) {

  lda_models_with_name <-
    ValenciaR::label_topics(
      lda_models,
      tax_table = rowData(mae[[count_assay]]) %>% as.data.frame() %>% mutate(Domain = Kingdom),
      distance = "BC", max_distance = 1
    )

  save(lda_models_with_name, file = lda_models_with_name_file)
} else {
  load(lda_models_with_name_file)
}
```

#### 4.2 Topic alignments

To identify which K offers the optimal description of the data, we rely on the alignment method. > TODO: cite alto paper

Because diagnosis scores vary with the maximal resolution of the alignment, we perform the alignment for several maximal resolutions (i.e., $K_{max} \in [15, 25]$).

```
alignment_dir <- str_c(LDA_models_and_alignments_dir, count_assay,"/alignments/")

align_topics_for_various_max_res(
  lda_models = lda_models_with_name,
  alignment_dir = alignment_dir,
  n_res = 11
  )
```

```
n_paths <- retrieve_n_paths_from_multiple_aligments(alignment_dir)
topics <- retrieve_topics_from_multiple_alignments(alignment_dir)
plot_alignment_diagnoses_from_multiple_alignments(n_paths, topics)
```

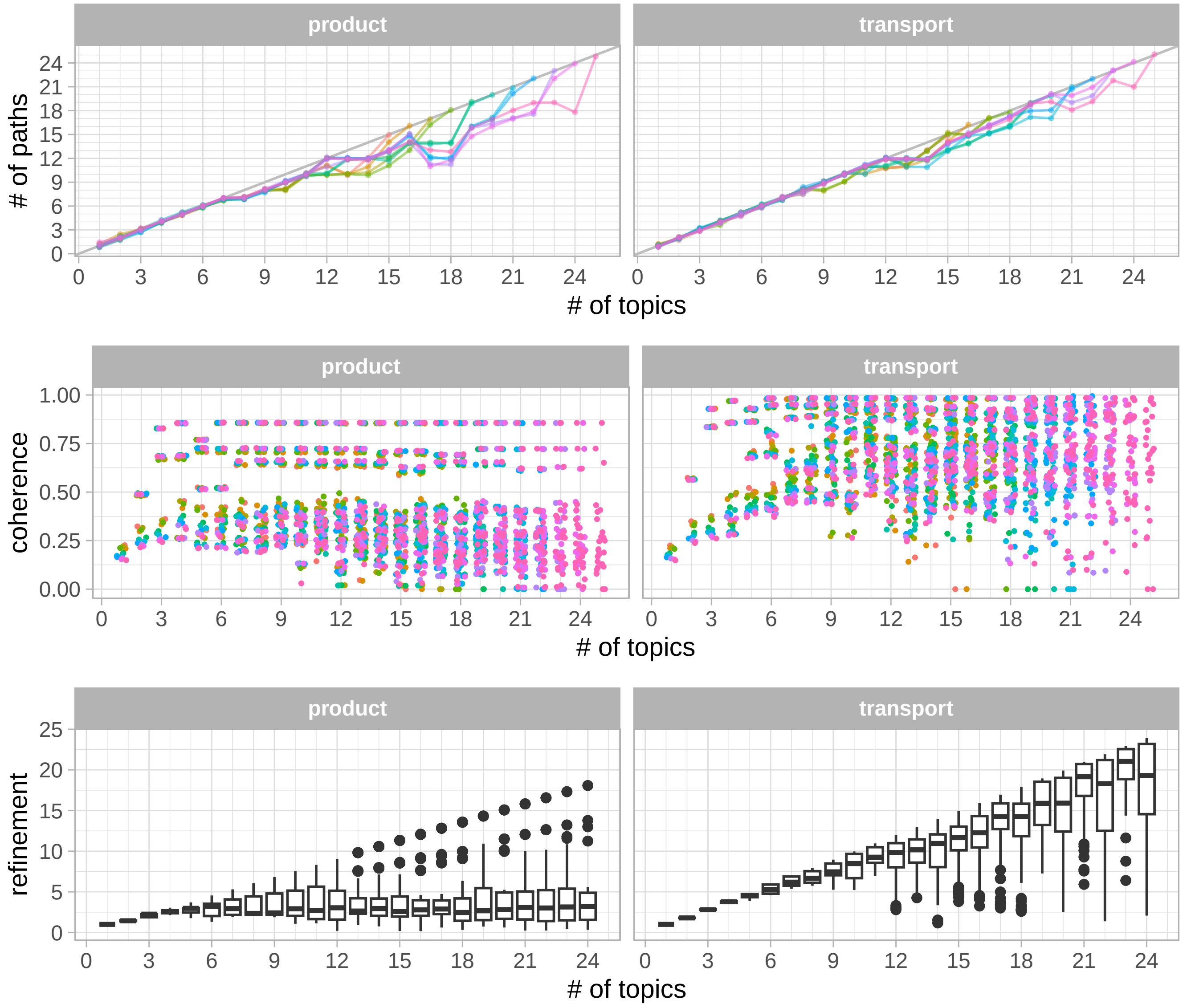

Figure 27: Alignment diagnostic scores. Colors denote the various alignments (several alignments were performed for increasing max K).

From the diagnostics scores from these various alignments, we identify the optimal $K$. We also identify a "coarser" resolution (a $K$ smaller than the optimal $K$) at which the refinement scores distributions are high.

```
best_K <- identify_optimal_K(n_paths = n_paths, topics = topics)

coarse_K <- identify_coarse_K(topics = topics, max_K = best_K - 4)

save(coarse_K, best_K, file = "../results/coarse_and_best_K.Rdata")
```

We visualize the alignments until the optimal K.

```
alignments_file <- str_c(alignment_dir, "alignments_best_K_", best_K,".Rdata")

if (!file.exists(alignments_file)) {
  transport_best_K <-
    alto::align_topics(models = lda_models_with_name[1:best_K], method = "transport")
  product_best_K <-
    alto::align_topics(models = lda_models_with_name[1:best_K], method = "product")
  save(transport_best_K, product_best_K, file = alignments_file)
} else {
```

```r
  load(file = alignments_file)
}

save(transport_best_K, file = "../results/transport_alignment_best_K.Rdata")
```

```r
path_to_topics <-
  transport_best_K@topics %>%
  filter(m == levels(m) %>% rev() %>% magrittr::extract(1)) %>%
  select(path, k_label)

g_transport <-
  alto::plot_alignment(
    transport_best_K,
    add_leaves = TRUE, min_feature_prop = 0.05, leaves_text_size = 6,
  ) +
  scale_color_manual(values = get_topic_colors(path_to_topics$k_label)) +
  scale_fill_manual(values = get_topic_colors(path_to_topics$k_label)) +
  expand_limits(x = 18)

path_to_topics <-
  product_best_K@topics %>%
  filter(m == levels(m) %>% rev() %>% magrittr::extract(1)) %>%
  select(path, k_label)

g_product <-
  alto::plot_alignment(
    product_best_K,
    add_leaves = TRUE, min_feature_prop = 0.05, leaves_text_size = 6,
    ) +
  scale_color_manual(values = get_topic_colors(path_to_topics$k_label)) +
  scale_fill_manual(values = get_topic_colors(path_to_topics$k_label)) +
  expand_limits(x = 18)

ggarrange(
  g_transport + ggtitle("Transport alignement"),
  g_product + ggtitle("Product alignement"),
  nrow = 1
)
```

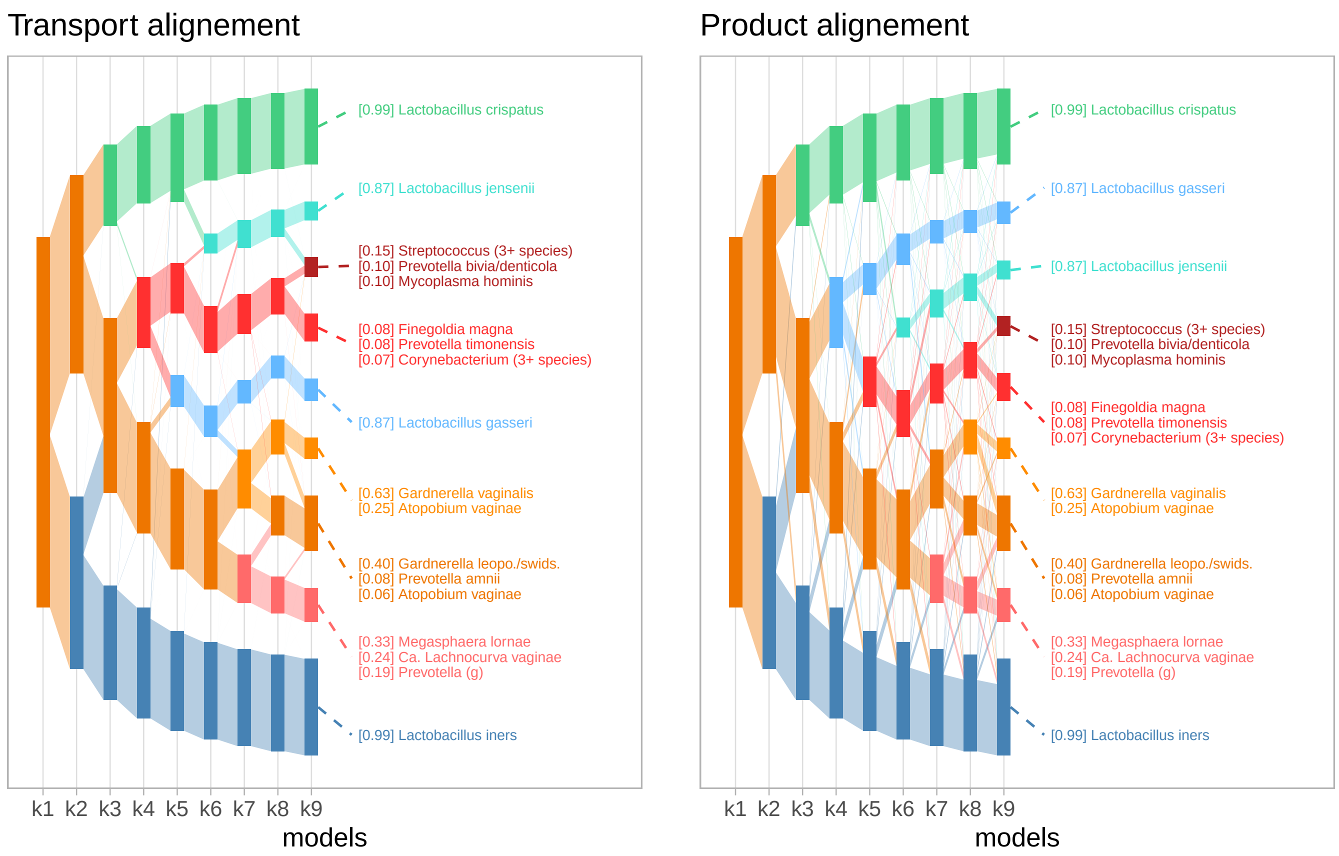

Figure 28: Topic alignments. Please note that topics might have different colors in both aligment.

We display the topic composition for $K = K^\star$ and $K = K^c$ (coarser view at high refinement score).

```
transport_best_K@topics <-
  transport_best_K@topics %>%
  mutate(
    k_label = k_label %>% factor(., levels = levels(k_label) %>% sort)
  )

path_to_topics <-
  transport_best_K@topics %>%
  filter(m == str_c("k", best_K)) %>%
  select(path, k_label) %>%
  arrange(path)

g_composition <-
  alto::plot_beta(
    transport_best_K,
    models = c(coarse_K, best_K),
    x_axis = "label",
    color_by = "path",
    threshold = 0.005
  ) +
  scale_color_manual(values = get_topic_colors(path_to_topics$k_label)) +
  guides(col = "none") +
  theme(strip.background = element_rect(color = "black", fill = "gray80"))
g_composition
```

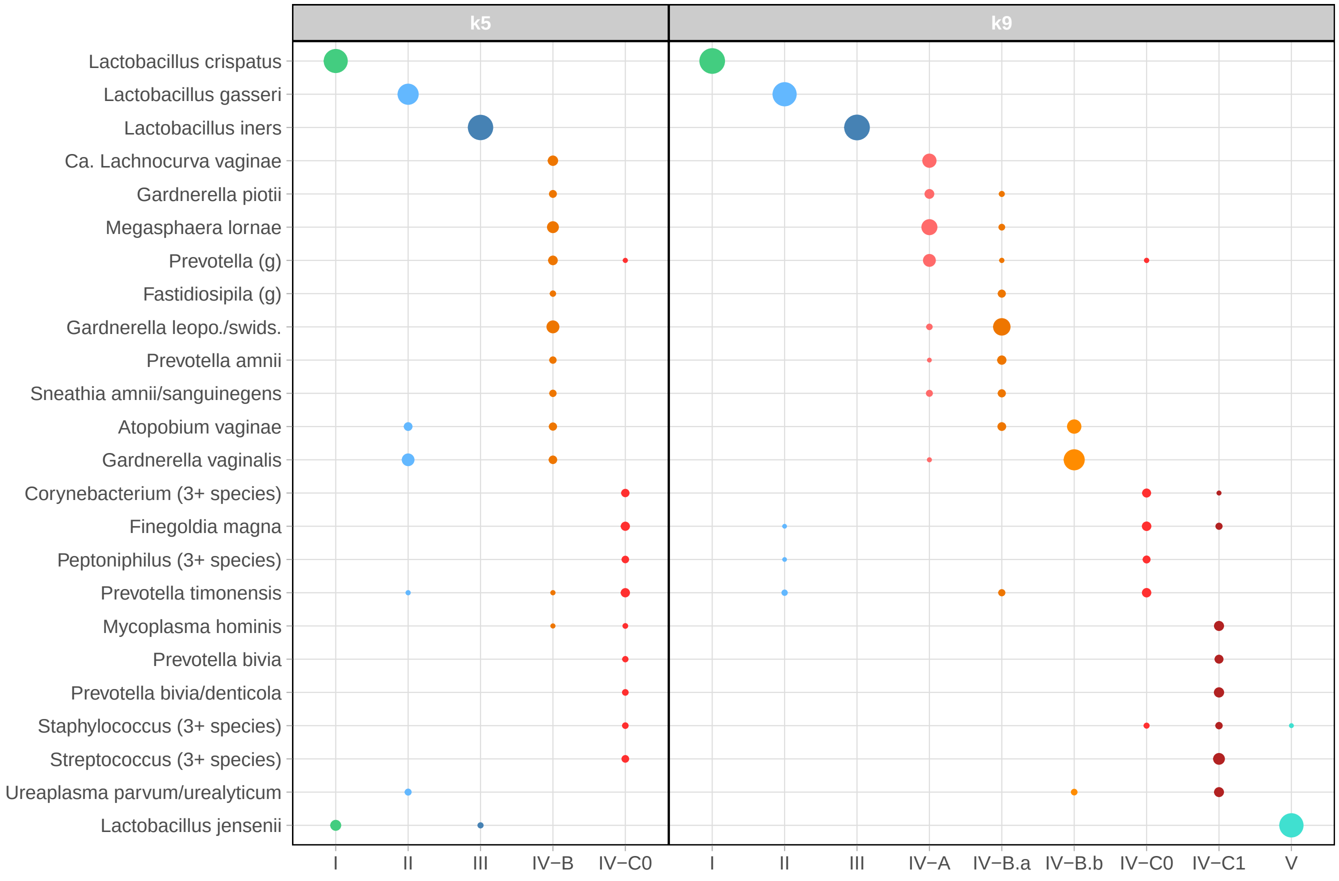

Figure 29: Topic composition for the optimal K and a coarser K

```
# g_composition <-
#    alto::plot_beta(
#     transport_best_K,
#     models = c(coarse_K, best_K),
#     x_axis = "label",
#     color_by = "path",
#     threshold = 0.005
#   ) +
#   scale_color_manual()
```

We also display topic alignments for higher max resolution than the optimal K so that we can also display topic coherence and refinement scores.

```
load(str_c(alignment_dir, "alignments_K_", 15,".Rdata"))

g_branches =
  plot_alignment(transport, add_leaves = TRUE, min_feature_prop = 0.02, leaves_text_size = 5) +
  theme_minimal() + expand_limits(x = 18)
g_coherence = plot(transport, color_by = "coherence") + theme_minimal()
g_refinement = plot(transport, color_by = "refinement") + theme_minimal()

ggpubr::ggarrange(
  plotlist = list(g_branches, g_coherence, g_refinement),
  nrow = 1, widths = c(1.4, 1, 1),
  labels = "auto"
)
```

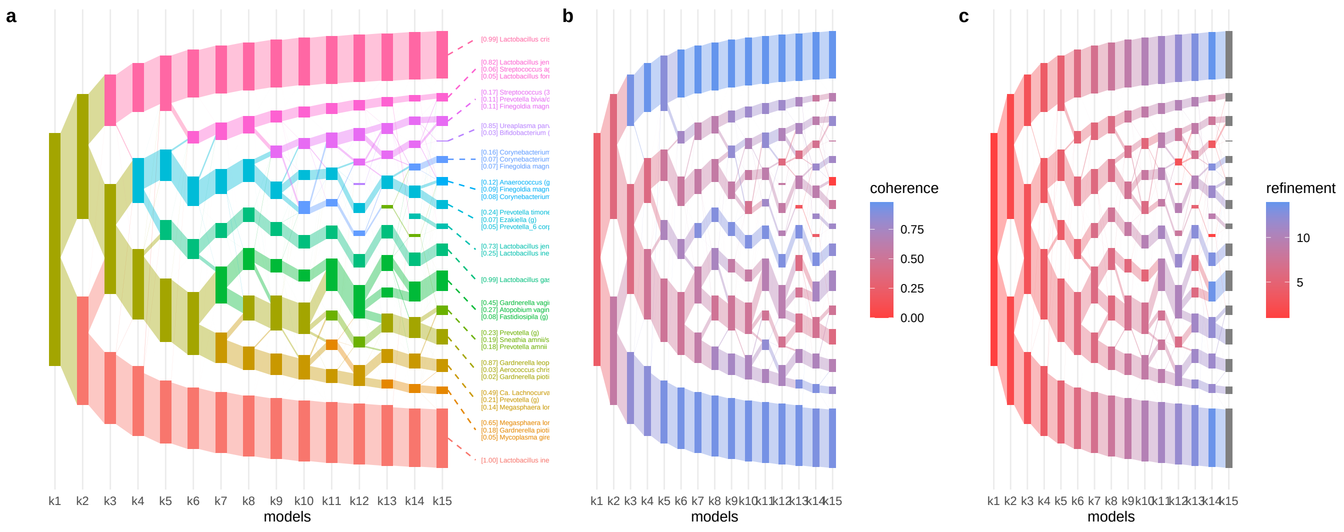

Figure 30: Topic alignment colored by paths (a), topic coherence (b), or topic refinement (c).

```r
df <-
  topics %>%
  filter(
    alignment == "product",
    K == min(K),
    m %in% str_c("k",c(coarse_K, best_K))
  ) %>%
  select(m, k, k_label, path, coherence, refinement) %>%
  pivot_longer(
    cols = c(coherence, refinement),
    names_to = "diagnosis",
    values_to = "score"
  )

path_to_topics <-
  df %>%
  filter(m == str_c("k",best_K)) %>%
  select(path, k_label) %>%
  distinct()
```

```r
g_coherence_refinement <-
  ggplot(df, aes(x = k_label, y = score, fill = path)) +
  geom_bar(stat = "identity") +
  facet_grid(diagnosis ~ m, scales = "free", space = "free_x") +
  guides(fill = "none") +
  scale_fill_manual(values = get_topic_colors(path_to_topics$k_label))

g_coherence_refinement
```

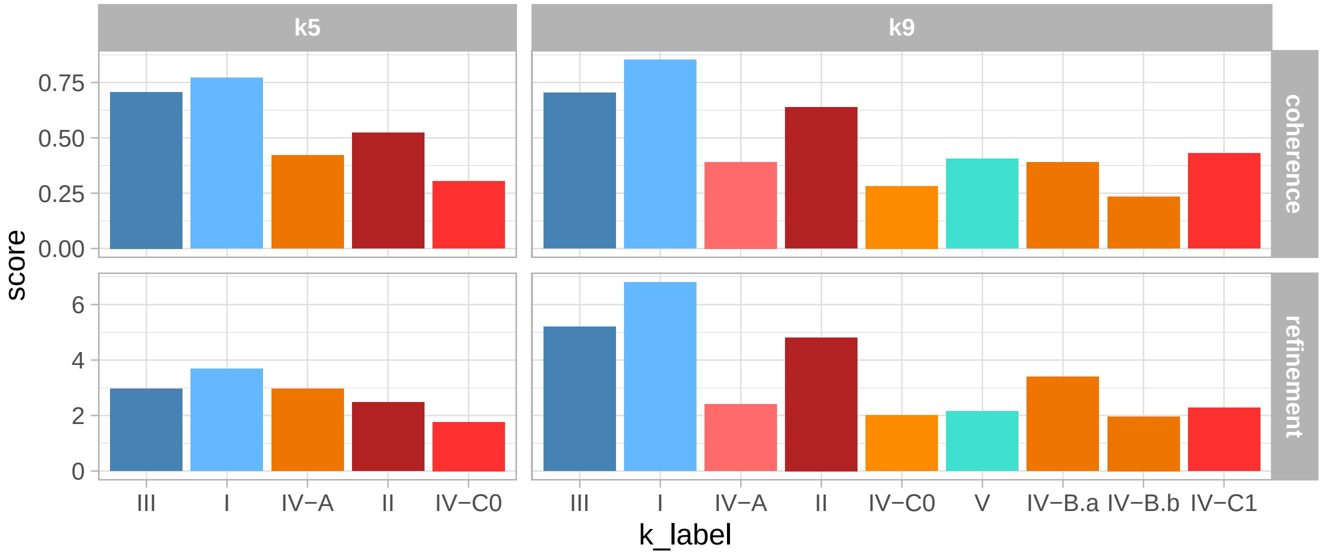

Figure 31: Coherence and refinement scores of topics at two resolutions.

```
gammas <-
  get_gammas_from_alignment(
    alignment = transport_best_K,
    m = best_K
  )

betas <-
  get_betas_from_alignment(
    alignment = transport_best_K,
    m = best_K
  )

save(gammas, file = "../results/gammas.Rdata")
```

#### 4.3 Comparison with sub-CSTs

##### 4.3.1 Valencia composition

```
valencia_centroids_mat <- get_valencia_centroids_mat()
valencia_centroids <-
  valencia_centroids_mat %>%
  as.data.frame() %>%
  mutate(subCST = rownames(valencia_centroids_mat)) %>%
  pivot_longer(., cols = -subCST, names_to = "taxa", values_to = "prop")

valencia_centroids <-
  valencia_centroids %>%
  arrange(taxa, -prop) %>%
  group_by(taxa) %>%
  mutate(main_subCST = subCST[1]) %>%
  ungroup() %>%
  arrange(main_subCST, -prop) %>%
  mutate(taxa = taxa %>% factor(., levels = unique(taxa)))

ggplot(valencia_centroids %>% filter(prop > 0.01),
       aes(x = subCST, y = taxa %>% fct_rev(), size = prop)) +
  geom_point(aes(col = subCST)) +
  scale_color_manual(values = subcst_colors) +
  ylab("")
```

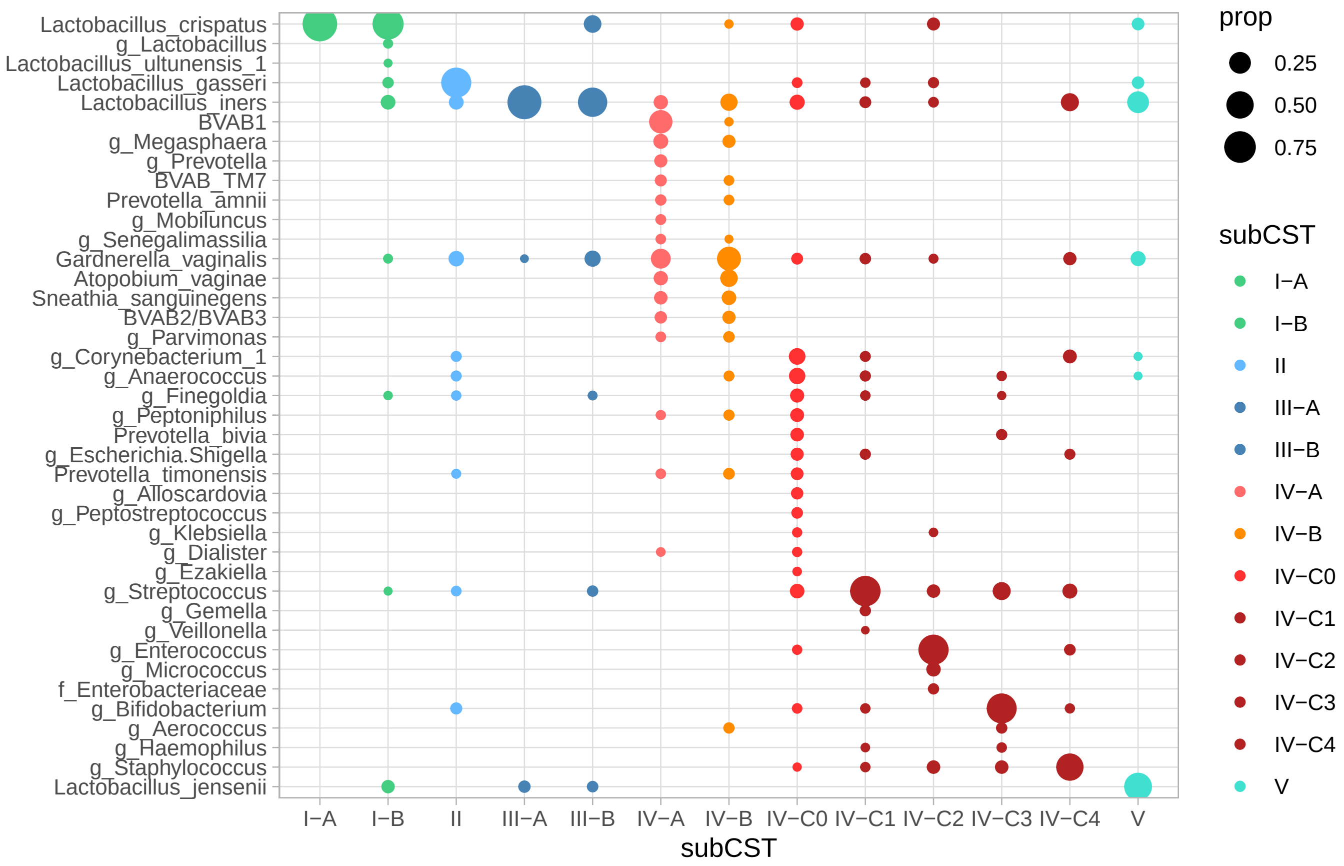

Figure 32: Composition of Valencia centroids. Adapted from France et al, 2020.

Valencia centroid compositions are less sparse than topic compositions (A lot of overlap between IV-A and IV-B)

##### 4.3.2 Composition comparison

To compare topic and sub-CST composition, we first need to harmonize taxonomic assignments.

```
converted_beta <-
  ValenciaR::convert_to_Valencia_taxonomy(
    lda_models_with_name[[best_K]]$beta %>% exp(),
    tax_table = rowData(mae[[count_assay]]) %>% as.data.frame() %>% mutate(Domain = Kingdom)
  )

valencia_taxa <- ValenciaR::get_Valencia_clusters()  %>% colnames()

taxonomic_dictionary <-
  converted_beta$conversion_table %>%
  as_tibble() %>%
  select(-tax_id, -Domain) %>%
  full_join(
    .,
    tibble(valencia_taxa_label = valencia_taxa),
    by = join_by(valencia_taxa_label)
    ) %>%
  mutate(valencia_taxa_label = valencia_taxa_label %>% factor(., levels = valencia_taxa)) %>%
  arrange(valencia_taxa_label)

write_csv(taxonomic_dictionary, "../results/valencia_taxonomic_dictionary.csv")
```

Then, we compute the Bray-Curtis dissimilarity between the topic and the sub-CST centroids.

```
distance_topics_subCSTs <-
  ValenciaR::assign_to_Valencia_clusters(
    input = converted_beta$converted_input, distance = "BC"
    )$distances

save(distance_topics_subCSTs, file = "../results/distance_topics_subCSTs.Rdata")

g_subCST_comparison <-
  plot_distance_topic_subCSTs(distance_topics_subCSTs, "BC")
g_subCST_comparison
```

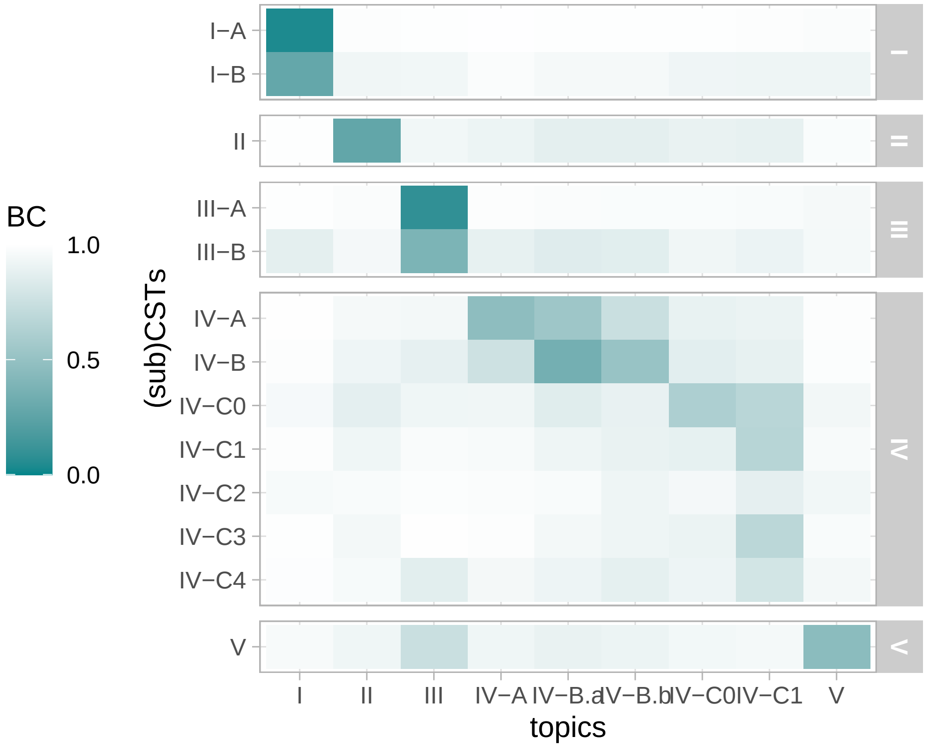

Figure 33: Bray-Curtis dissimilarity between Valencia centroids and topics.

```
distance_topics_subCSTs <-
  compute_distance_topics_subCSTs(
    beta_mat = lda_models_with_name[[best_K]]$beta %>% exp(),
    valencia_centroids_mat =  valencia_centroids_mat,
    tax_table = rowData(mae[[count_assay]]) %>% as.data.frame()
    )

save(distance_topics_subCSTs, file = "../results/distance_topics_subCSTs.Rdata")

g_subCST_comparison <- plot_distance_topic_subCSTs(distance_topics_subCSTs)
g_subCST_comparison
```

##### 4.3.3 Sample composition accuracy

We next evaluate if topics or sub-CST provide a more accurate representation of the sample composition.

To do that, we compare the Bray-Curtis dissimilarity between the actual sample compositions and those predicted by sub-CSTs or by topics. For sub-CSTs, the predicted sample composition is simply the composition of the sub-CST centroids. For topics, the predicted sample composition is the matrix product between the gamma and beta matrices.

```
sample_composition_accuracy <-
  get_sample_composition_prediction_dissimilarity(
    mae, count_assay, best_lda_model = lda_models_with_name[[best_K]]
    )
```

```
df_wide <-
  sample_composition_accuracy$sample_composition_accuracy %>%
  pivot_wider(
    id_cols = c(SampleID, Status, subCST),
    names_from = method, values_from = BC_dissimilarity
  ) %>%
  mutate(
```

```
    diff = subCSTs - topics
  ) %>%
  left_join(colData(mae) %>% as.data.frame() %>% select(SampleID, Subject), by = "SampleID")

# Because the sample size is very large (2179 for Pregnant participants and 1534 for non-pregnant ones)
# we can use a z-test for testing that the mean difference is greater than 0.
# at this sample size, z-tests and t-tests are approximately the same because the uncertainty on the standard
ttest_P <- t.test(x = df_wide$diff[df_wide$Status == "Pregnant"], alternative = "greater")
ttest_P
```

```
##
##  One Sample t-test
##
## data:  df_wide$diff[df_wide$Status == "Pregnant"]
## t = 51.112, df = 2178, p-value < 2.2e-16
## alternative hypothesis: true mean is greater than 0
## 95 percent confidence interval:
##  0.1133572       Inf
## sample estimates:
## mean of x
## 0.1171282
```

```
ttest_NP <- t.test(x = df_wide$diff[df_wide$Status == "Non-pregnant"], alternative = "greater")
ttest_NP
```

```
##
##  One Sample t-test
##
## data:  df_wide$diff[df_wide$Status == "Non-pregnant"]
## t = 4.2462, df = 1533, p-value = 1.152e-05
## alternative hypothesis: true mean is greater than 0
## 95 percent confidence interval:
##  0.01275611        Inf
## sample estimates:
##  mean of x
## 0.02082978
```

```
tests <-
  tibble(
    Status = c("Pregnant", "Non-pregnant"),
    p_val = c(ttest_P$p.value, ttest_NP$p.value)
  )

j_NP <- which((df_wide$Status == "Non-pregnant") &  !str_detect(df_wide$subCST, "C[1-4]"))
t.test(x = df_wide$diff[j_NP], alternative = "greater")
```

```
##
##  One Sample t-test
##
## data:  df_wide$diff[j_NP]
## t = 23.294, df = 1373, p-value < 2.2e-16
## alternative hypothesis: true mean is greater than 0
## 95 percent confidence interval:
##  0.06435692        Inf
## sample estimates:
##  mean of x
## 0.06925021
```

```
# we could also use non-parametric tests
# wilcox.test(x = df_wide$diff[df_wide$Status == "Pregnant"], alternative = "greater")
```

```
# wilcox.test(x = df_wide$diff[df_wide$Status == "Non-pregnant"], alternative = "greater")
# wilcox.test(x = df_wide$diff[j_NP], alternative = "greater")
```

There are 10% of sub-CST IV-C1-4 samples in non-pregnant participants.

```
plot_sample_composition_accuracy(
  sample_composition_accuracy$sample_composition_accuracy,
  tests = tests
  )
```

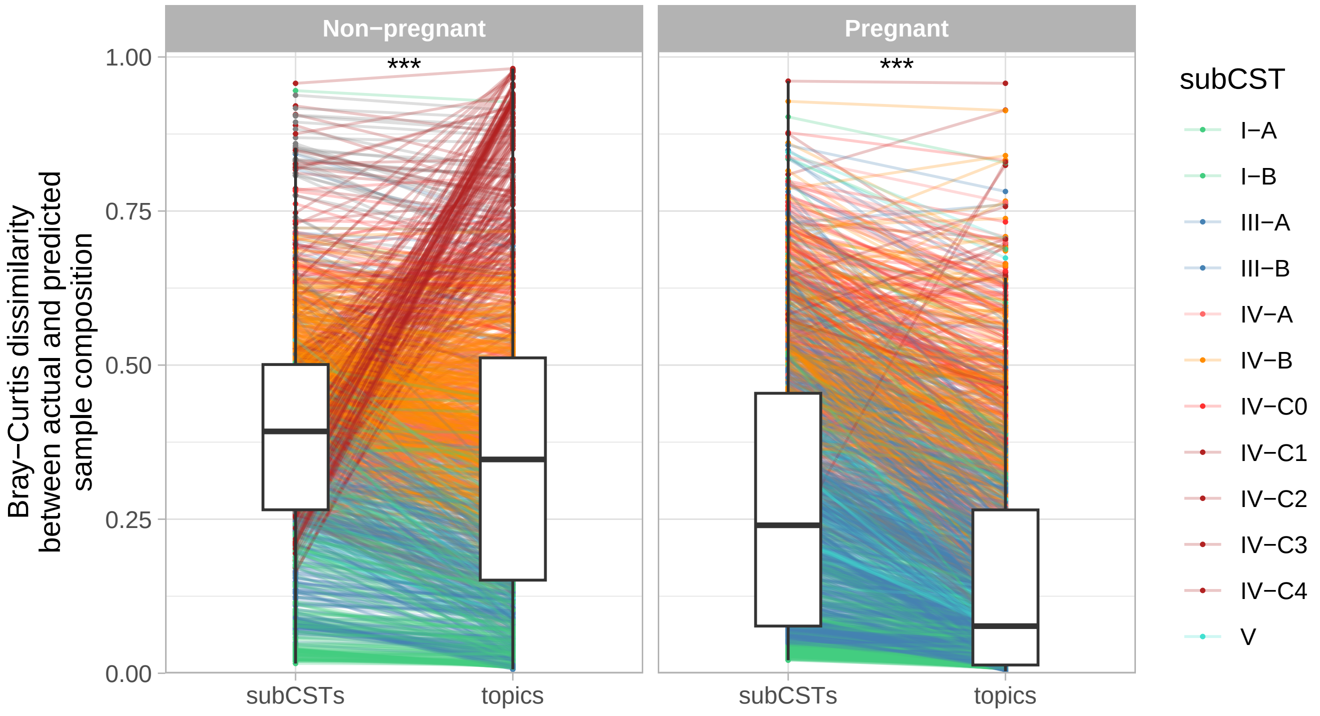

Figure 34: Bray-Curtis dissimilarity between the actual sample compositions and those predicted by topic mixed membership or sub-CST membership.

```
save(sample_composition_accuracy, tests, file = "../results/sample_composition_accuracy.Rdata")
```

**Does it depend on stability?**

To test that, we define a local stability score based on the Bray-Curtis dissimilarity between consecutive samples, then determine if samples belong to a "stable" vs "transition" state. We then evaluate the sample composition descriptive accuracy of topics and sub-CST in these two states.

```
local_stability <- compute_sample_stability(mae, count_assay, BC_threshold = 0.25)

tmp <-
  sample_composition_accuracy$sample_composition_accuracy %>%
  left_join(local_stability, by = join_by(SampleID, Status, subCST))
```

```
tmp_wide <-
  tmp %>%
  filter(!is.na(state)) %>%
  pivot_wider(
    id_cols = c(SampleID, Status, subCST, state),
    names_from = method,
    values_from = BC_dissimilarity
  ) %>%
  mutate(diff = subCSTs - topics)
```

```
tests_for_each_group <-
  tmp_wide %>%
  group_by(Status, state) %>%
  summarize(
    mean_diff = mean(diff),
    p_val = t.test(x = subCSTs, y = topics, paired = TRUE, alternative = "greater")$p.value
  )

ggplot(tmp %>% filter(!is.na(state)), # , !str_detect(subCST, "C[1-4]")
       aes(x = method, y = BC_dissimilarity)) +
  geom_line(aes(group = SampleID, col = subCST), alpha = 0.25, size = 0.3) +
  geom_point(aes(col = subCST), size = 0.3) +
  geom_boxplot(width = 0.5, outlier.shape = NA) +
  facet_grid(Status ~ state) +
  scale_color_manual(values = subcst_colors)
```

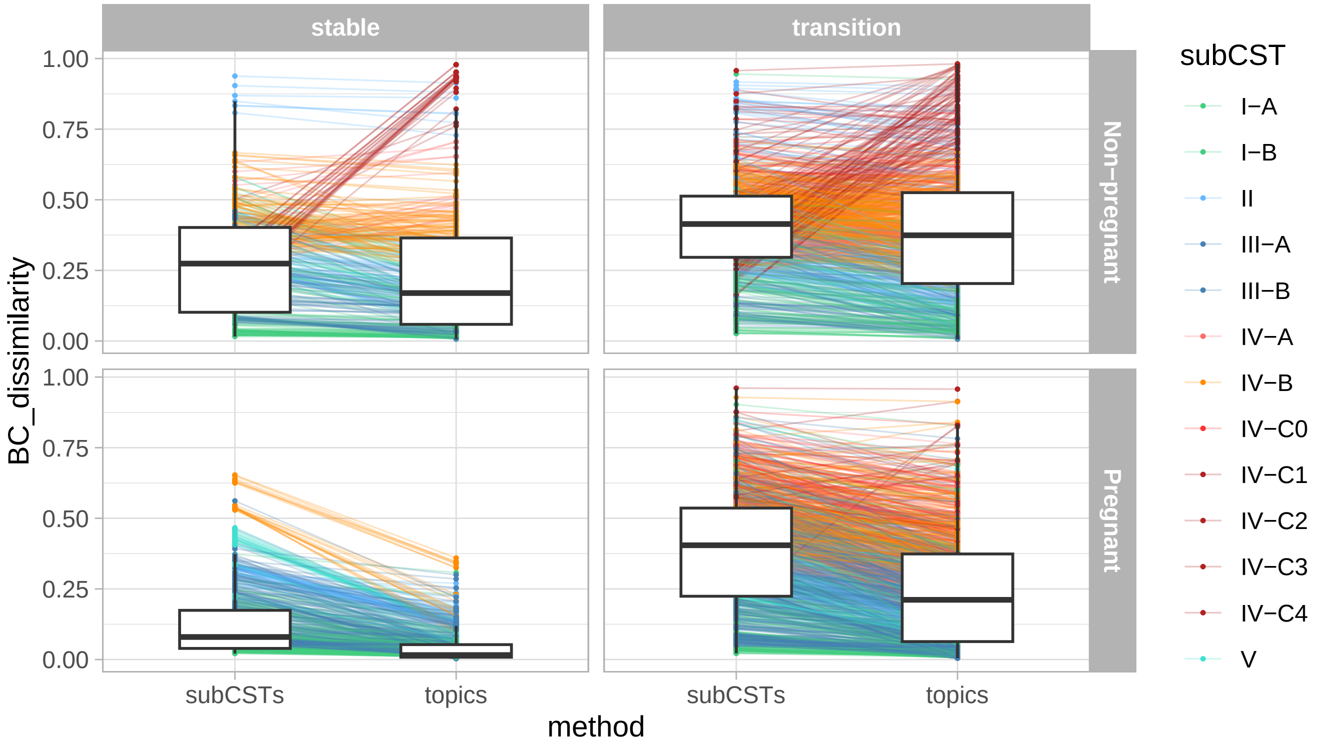

```
library(ggbeeswarm)

g_local_stability_and_topics_vs_clusters <-
  ggplot(tmp_wide, aes(x = state, y = diff)) +
  # geom_jitter(aes(col = subCST), size = 0.3, width = 0.25) +
  geom_beeswarm(aes(col = subCST), size = 0.3, cex = 0.5) +
  geom_boxplot(outlier.shape = NA, alpha = 0.5) + # , linewidth = 2
  facet_grid(. ~ Status) +
  scale_color_manual(values = subcst_colors) +
  ylab("BC(subCSTs) - BC(topics)") +
  xlab("")

save(
  g_local_stability_and_topics_vs_clusters,
  file = "../results/suppl_figs/local_stability_and_topics_vs_clusters.Rdata"
)
```

```
g_local_stability_and_topics_vs_clusters
```

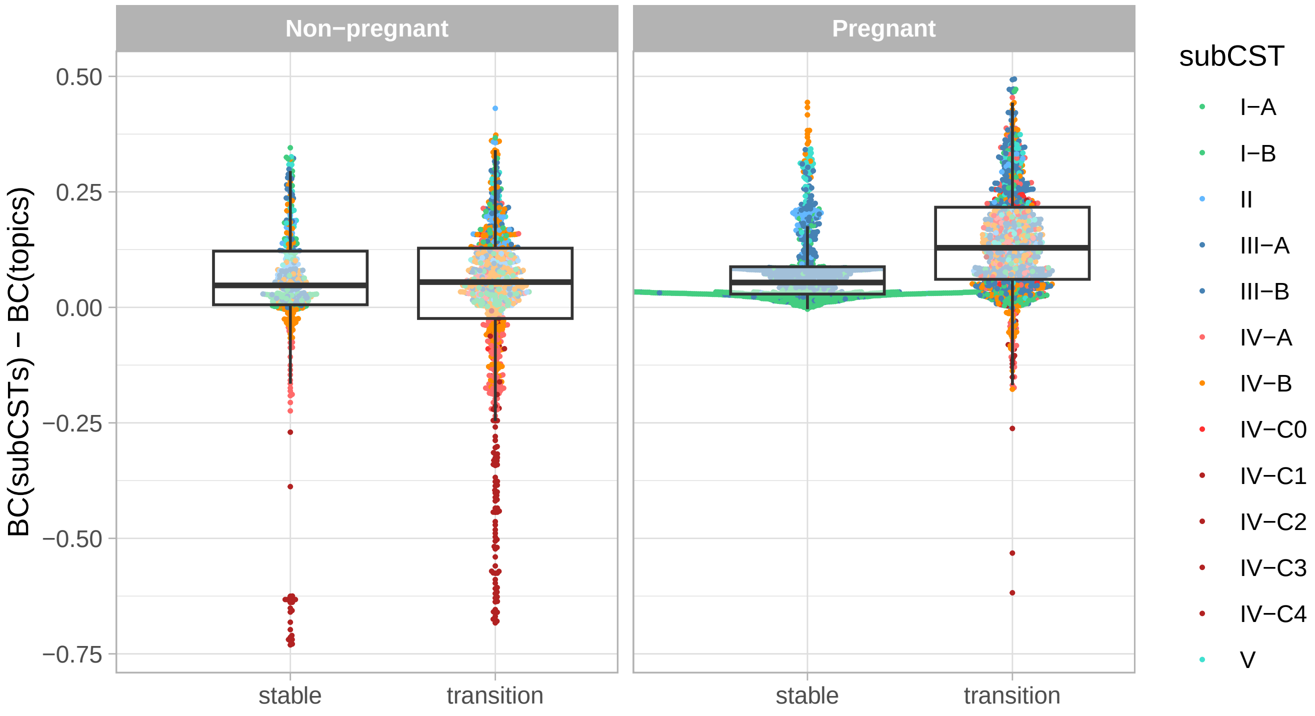

Figure 35: Distribution of the differences between the Bray-Curtis dissimilarity between actual sample compositon and that predicted by sub-CST and topics for samples in stable microbiota episodes or samples belonging to transitions or unstable microbiotas.

```
t_NP <-
  t.test(
  x = tmp_wide$diff[(tmp_wide$Status == "Non-pregnant") & (tmp_wide$state == "stable")],
  y = tmp_wide$diff[(tmp_wide$Status == "Non-pregnant") & (tmp_wide$state != "stable")],
  alternative = "less"
  )

t_P <-
t.test(
  x = tmp_wide$diff[(tmp_wide$Status == "Pregnant") & (tmp_wide$state == "stable")],
  y = tmp_wide$diff[(tmp_wide$Status == "Pregnant") & (tmp_wide$state != "stable")],
  alternative = "less"
  )

t_BC_25 <-
bind_rows(
  tibble(Status = "Non-pregnant",
         `mean(stable)` = t_NP$estimate[1] %>% round(., 4),
         `mean(transitions)` = t_NP$estimate[2] %>% round(., 4),
         `diff. means` = diff(t_NP$estimate),
         `p-value` = get_sign_levels(t_NP$p.value)),
  tibble(Status = "Pregnant",
         `mean(stable)` = t_P$estimate[1] %>% round(., 4),
         `mean(transitions)` = t_P$estimate[2] %>% round(., 4),
         `diff. means` = diff(t_P$estimate),
         `p-value` = get_sign_levels(t_P$p.value))
)
```

```r
t_BC_25 %>%  kable(., format = "latex", booktab = TRUE, caption = "Average differences between the Bray-curti
  kableExtra::kable_styling(latex_option = "HOLD_position")
```

Table 11: Average differences between the Bray-curtis dissimilarity (2nd and 3rd columns) between actual and predicted sample composition when sample composition is predicted with topic or cluster membership(s) in samples belonging to stable microbiotas vs transition states from pregnant or non-pregnant participants (1st column). The 5th column provides the p-value from a one-sided t-test.

| Status | mean(stable) | mean(transitions) | diff. means | p-value |
|---|---|---|---|---|
| Non-pregnant | 0.0300 | 0.0231 | -0.0068986 | > 0.1 |
| Pregnant | 0.0832 | 0.1422 | 0.0589620 | ≤ 0.001 |

In pregnant individuals, samples in stable microbiotas are almost equally well described by topics vs CSTs, while those in transitions/unstable microbiotas are much better described by topics. In non-pregnant individuals, it does not matter if the sample belongs to a stable vs unstable microbiota.

**Sensitiviy analysis for various thresholds for stability**

```r
local_stability_015 <- compute_sample_stability(mae, count_assay, BC_threshold = 0.15)

tmp_wide_015 <-
  sample_composition_accuracy$sample_composition_accuracy %>%
  left_join(local_stability_015, by = join_by(SampleID, Status, subCST)) %>%
  filter(!is.na(state)) %>%
  pivot_wider(
    id_cols = c(SampleID, Status, subCST, state),
    names_from = method,
    values_from = BC_dissimilarity
  ) %>%
  mutate(diff = subCSTs - topics)

t_NP_015 <-
  t.test(
  x = tmp_wide_015$diff[(tmp_wide_015$Status == "Non-pregnant") & (tmp_wide_015$state == "stable")],
  y = tmp_wide_015$diff[(tmp_wide_015$Status == "Non-pregnant") & (tmp_wide_015$state != "stable")],
  alternative = "less"
  )

t_P_015 <-
t.test(
  x = tmp_wide_015$diff[(tmp_wide_015$Status == "Pregnant") & (tmp_wide_015$state == "stable")],
  y = tmp_wide_015$diff[(tmp_wide_015$Status == "Pregnant") & (tmp_wide_015$state != "stable")],
  alternative = "less"
  )

local_stability_035 <- compute_sample_stability(mae, count_assay, BC_threshold = 0.35)

tmp_wide_035 <-
  sample_composition_accuracy$sample_composition_accuracy %>%
  left_join(local_stability_035, by = join_by(SampleID, Status, subCST)) %>%
  filter(!is.na(state)) %>%
  pivot_wider(
    id_cols = c(SampleID, Status, subCST, state),
    names_from = method,
    values_from = BC_dissimilarity
```

```r
  ) %>%
  mutate(diff = subCSTs - topics)

t_NP_035 <-
  t.test(
  x = tmp_wide_035$diff[(tmp_wide_035$Status == "Non-pregnant") & (tmp_wide_035$state == "stable")],
  y = tmp_wide_035$diff[(tmp_wide_035$Status == "Non-pregnant") & (tmp_wide_035$state != "stable")],
  alternative = "less"
  )

t_P_035 <-
t.test(
  x = tmp_wide_035$diff[(tmp_wide_035$Status == "Pregnant") & (tmp_wide_035$state == "stable")],
  y = tmp_wide_035$diff[(tmp_wide_035$Status == "Pregnant") & (tmp_wide_035$state != "stable")],
  alternative = "less"
  )
```

```r
t_sensitivity <-
  bind_rows(
  tibble(threshold = 0.15, Status = "Non-pregnant",
         `mean BC diff. (stable microbiotas)` = t_NP_015$estimate[1] %>% round(., 4),
         `mean BC diff. (transitions)` = t_NP_015$estimate[2] %>% round(., 4),
         `p-value` = get_sign_levels(t_NP_015$p.value)),
  tibble(threshold = 0.15, Status = "Pregnant",
         `mean BC diff. (stable microbiotas)` = t_P_015$estimate[1] %>% round(., 4),
         `mean BC diff. (transitions)` = t_P_015$estimate[2] %>% round(., 4),
         `p-value` = get_sign_levels(t_P_015$p.value)),
  t_BC_25 %>% mutate(threshold = 0.25) %>% select(-`diff. means`) %>%
    dplyr::rename(`mean BC diff. (stable microbiotas)` = `mean(stable)`,
           `mean BC diff. (transitions)` = `mean(transitions)`),
  tibble(threshold = 0.35, Status = "Non-pregnant",
         `mean BC diff. (stable microbiotas)` = t_NP_035$estimate[1] %>% round(., 4),
         `mean BC diff. (transitions)` = t_NP_035$estimate[2] %>% round(., 4),
         `p-value` = get_sign_levels(t_NP_035$p.value)),
  tibble(threshold = 0.35, Status = "Pregnant",
         `mean BC diff. (stable microbiotas)` = t_P_035$estimate[1] %>% round(., 4),
         `mean BC diff. (transitions)` = t_P_035$estimate[2] %>% round(., 4),
         `p-value` = get_sign_levels(t_P_035$p.value))
)

save(t_sensitivity, file = "../results/suppl_figs/table_sensitivity.Rdata")

t_sensitivity %>%  kable(., format = "latex", booktab = TRUE, caption = "Average differences between the Bray
  kableExtra::kable_styling(latex_option = "HOLD_position")
```

Table 12: Average differences between the Bray-curtis dissimilarity (3rd and 4th columns) between actual and predicted sample composition when sample composition is predicted with topic or cluster membership(s) for various thresholds (1st column) differentiating between stable microbiotas and transition states in samples from pregnant or non-pregnant participants (2nd column). The 5th column provides the p-value from a one-sided t-test.

| threshold | Status | mean BC diff. (stable microbiotas) | mean BC diff. (transitions) | p-value |
|---|---|---|---|---|
| 0.15 | Non-pregnant | 0.0453 | 0.0225 | > 0.1 |
| 0.15 | Pregnant | 0.0638 | 0.1421 | ≤ 0.001 |
| 0.25 | Non-pregnant | 0.0300 | 0.0231 | > 0.1 |
| 0.25 | Pregnant | 0.0832 | 0.1422 | ≤ 0.001 |
| 0.35 | Non-pregnant | 0.0436 | 0.0042 | > 0.1 |
| 0.35 | Pregnant | 0.0999 | 0.1361 | ≤ 0.001 |

##### 4.3.4 Predicting the risk of losing Lactobacillus dominance at the next sample

Next, we test whether describing samples in terms of topic membership better predict the risk of losing Lactobacillus dominance at the next sample, than sub-CST would.

To do that, we train two logistic regression models to predict whether the next sample is still Lactobacillus-dominated or not. One model's input variable is the subCST category, the other model's input variables are the topic partial memberships. Because we predict rare events, we use weights in our logistic regression models such that transitions have higher weights.

Then, we evaluate the predictions from both methods on a test set and compute the F1 score (harmonic mean between precision and recall), a metric suited to evaluate the prediction performances for rare events.

We repeat this on 10 independent random splits for our training and test sets and report the distribution of the F1 scores for each method.

```
load(file = "../results/gammas.Rdata")
```

```
input_data <-
  prepare_input_data_for_transition_predictions(mae, gammas) %>%
  filter(cat == "L", !str_detect(subCST, "IV-"))

set.seed(1)
res <- purrr::map_dfr(
  .x = 1:10,
  .f = train_test_model_transition_predictions,
  input_data = input_data
)
```

```
## 1 |  2 |     3 |     4 |     5 |     6 |     7 |     8 |     9 |     10 |
```

```
F1 <- compute_F1_score(res)
F1_long <- get_F1_long(F1)

save(res, F1, file = "../results/data_for_figures/risk_predictions.RData")
```

```
g_pred_metrics <-
  ggplot(F1_long, aes(x = Predictor, y = value, fill = Predictor)) +
  geom_point(size = 0.5) +
  geom_line(aes(group = i), alpha = 0.25) +
  geom_boxplot(alpha = 0.8) +
  xlab("") +
  guides(fill = "none") +
  facet_wrap(metric ~ ., scales = "free")

save(g_pred_metrics, file = "../results/suppl_figs/pred_metrics.Rdata")
```

```
g_pred_metrics
```

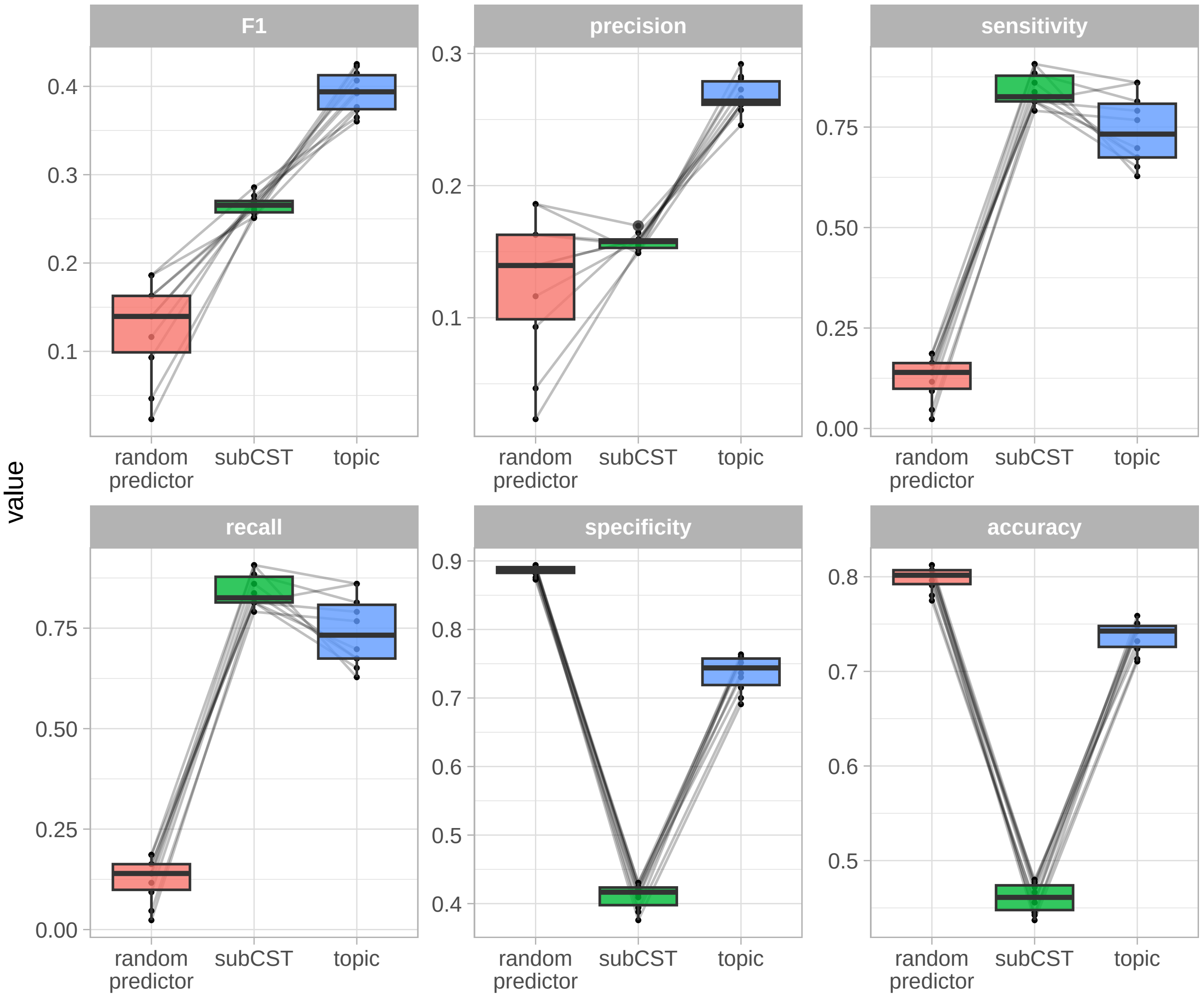

Figure 36: Prediction metrics (panels) for each method (x-axis)

```
wilcox_test_metric(F1, metric = "F1") %>%
  kable(booktab = TRUE, format = "latex", caption = "Median F1 scores between sub-CST-based and topic-based p
  kableExtra::kable_styling(latex_option = "HOLD_position")
```

Table 13: Median F1 scores between sub-CST-based and topic-based predictions

| median(subCST) | median(topic) | p-value |
|---:|---:|---:|
| 0.2654079 | 0.3938185 | 0.0019531 |

```
wilcox_test_metric(F1, metric = "precision") %>%
  kable(booktab = TRUE, format = "latex", caption = "Median precision between sub-CST-based and topic-based p
  kableExtra::kable_styling(latex_option = "HOLD_position")
```

Table 14: Median precision between sub-CST-based and topic-based predictions

| median(subCST) | median(topic) | p-value |
|---|---|---|
| 0.157642 | 0.2639346 | 0.0019531 |

```
g_F1 <-
  ggplot(F1 %>% filter(predictor != "random predictor"),
         aes(x = predictor, y = F1, fill = predictor)) +
  geom_line(aes(group = i), linewidth = 0.5, alpha = 0.25) +
  geom_boxplot(alpha = 0.9) +
  ylab("F1 score (harmonic mean of precision and sensitivity)") +
  scale_fill_manual(values = c("darkolivegreen3","turquoise4")) +
  guides(fill = "none")
```

```
F1_subCST <- get_F1_by_subCST(res, input_data)

ggplot(F1_subCST %>% filter(predictor != "random predictor"),
       aes(x = predictor, y = F1, fill = predictor)) +
  geom_line(aes(group = i), size = 0.5, alpha = 0.25) +
  geom_boxplot(alpha = 0.9) +
  facet_grid(. ~ subCST) +
  scale_fill_manual(values = c("darkolivegreen3","turquoise4")) +
  guides(fill = "none")
```

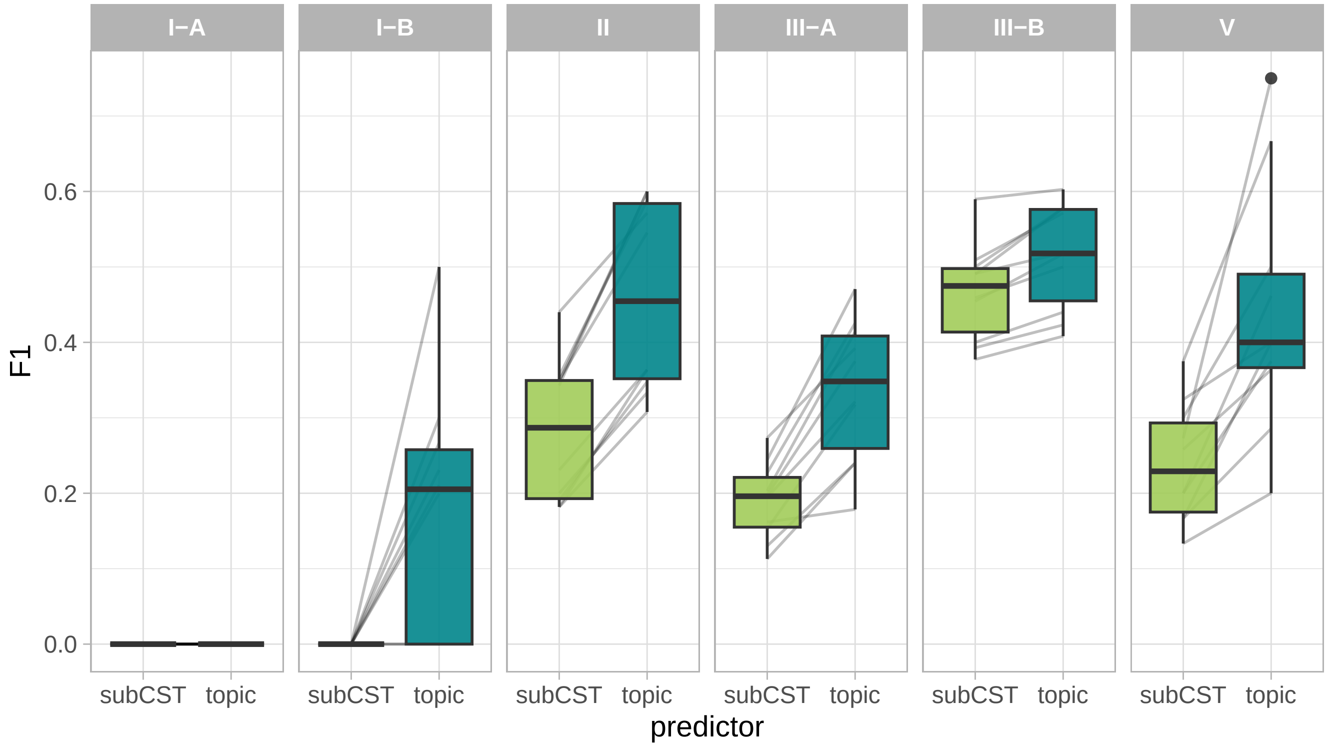

Figure 37: F1 score per sub-CST for each method.

We can also compare the predicted probabilities of losing *Lactobacillus* dominance for the samples which are *Lactobacillus* dominated.

To test whether these probabilities significantly differ, we use non-parametric tests: the paired sign test for testing differences between the probabilities predicted by subCST vs topics, and the Wilcox (Mann-Whitney) test for testing differences in predicted probabilities by topics in samples that actually lose their Lactobacillus dominance or not.

```r
input_data <- prepare_input_data_for_transition_predictions(mae, gammas) %>% filter(cat == "L")

cst_model <- glm(next_cat ~ subCST + 0, data = input_data,
                 weights = 1 + 10*(input_data$next_cat != input_data$cat),
                 family = "binomial")

topic_model <- glm(next_cat ~ . + 0,
                   data = input_data %>% select(next_cat, starts_with("k_")),
                   weights = 1 + 10*(input_data$next_cat != input_data$cat),
                   family = "binomial")

res <-
  input_data %>%
  mutate(
    predicted_risk_cst = cst_model$fitted.values,
    predicted_risk_topics = topic_model$fitted.values,
  ) %>%
  filter(cat == "L")

res_long <-
  bind_rows(
    input_data %>%
      mutate(predicted_risk = cst_model$fitted.values, predictor = "subCSTs"),
    input_data %>%
      mutate(predicted_risk = topic_model$fitted.values, predictor = "Topics")
  ) %>%
  filter(cat == "L")

tests <- test_significance_transitions(res)
```

```r
plot_predicted_risk(res_long, tests)
```

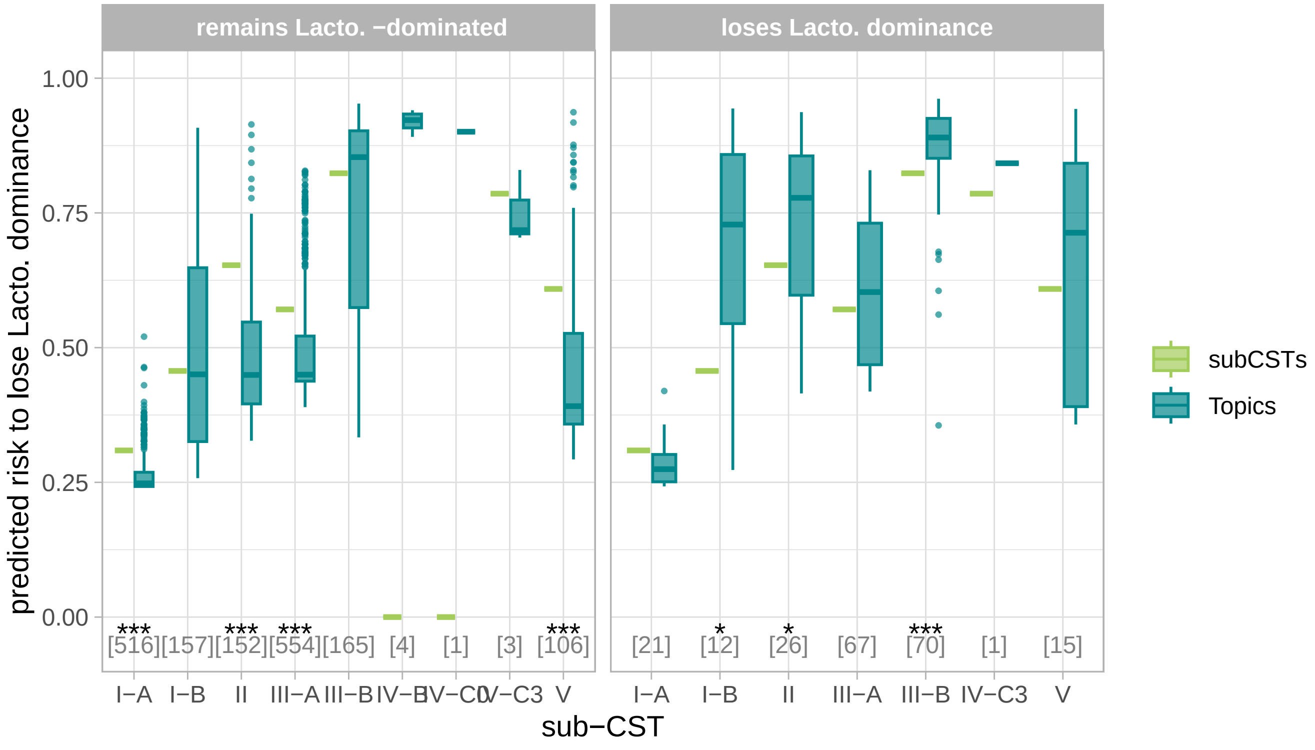

Figure 38: Predicted risk of losing lactobacillus dominance per method and CST.

Table 15: p-value and adjusted p-value of the Wilcox test comparing the median predicted risk of losing Lactobacillus dominance in samples that actually do or do not lose Lactobacillus dominance.

| sub-CST | p-value Wilcox test | adjusted p-value Wilcox test |
|---|---:|---:|
| I-A | 0.0001406 | 0.0005168 |
| I-B | 0.0031376 | 0.0076870 |
| II | 0.0000000 | 0.0000000 |
| III-A | 0.0000000 | 0.0000000 |
| III-B | 0.0000117 | 0.0000575 |
| V | 0.0019233 | 0.0056545 |

We observe that for each sub-CSTs, topic predict significantly lower probability of losing Lactobacillus dominance when dominance is not lost or higher probability of losing Lactobacillus dominance when it is lost.

Further, for each sub-CST, there is a significant difference between the predicted risk for samples followed by a loss of Lactobacillus dominance or not.

```
tests$wilcox %>%
  set_colnames(c("sub-CST", "p-value Wilcox test", "adjusted p-value Wilcox test")) %>%
  kable(., format = "latex", booktab = TRUE, caption = "p-value and adjusted p-value of the Wilcox test compa
```

#### 4.4 Associations with demographics, cohorts, and reproductive state

##### 4.4.1 Association with race, study site location, and pregnancy Status

In this section, we test whether demographic variables (Race and Study site location) or pregnancy status is associated with differential topic composition. To do so, we use a Dirichlet regression since topics are compositional data and use the Race, Site, and Status as input variable.

```
si <-
  colData(mae) %>%
  as.data.frame() %>%
  as_tibble() %>%
  select(Subject, SampleID, Race, Site, Status) %>%
    filter(!(Site %in% c("AYAC","EM"))) %>%
  mutate(
    Site =
      case_when(Site == "Stanford" ~ "SU", TRUE ~ Site) %>%
      factor(., levels = c("UAB","SU")),
    Race =
      ifelse(Race %in% c("Asian","Hispanic/Latino"), "Other", Race) %>%
      factor(.,levels =c("Other","Black","White")),
    Status = Status %>% factor(., levels = c("Non-pregnant","Pregnant"))
  ) %>%
  dplyr::rename(d = SampleID)
```

Because not all participants have the same number of samples, we aggregate topic composition at the participant-level and use the mean over all their time-points.

```
g_race_composition <-
  plot_topic_composition_per_group_and_subject(
    gammas = gammas,
    groups = colData(mae) %>% as.data.frame() %>%
      select(Subject, SampleID, Race) %>%
      mutate(Race = ifelse(Race %in% c("Asian","Hispanic/Latino"), "Other", Race)) %>% # there are not enough
      dplyr::rename(d = SampleID,
                    group = Race) %>%
      mutate(group_name = "Race",
             group = group %>% factor(., levels = c("Other","Black","White")))
      ) +
  scale_fill_manual(values = get_topic_colors(gammas$k %>% levels()))

g_race_composition
```

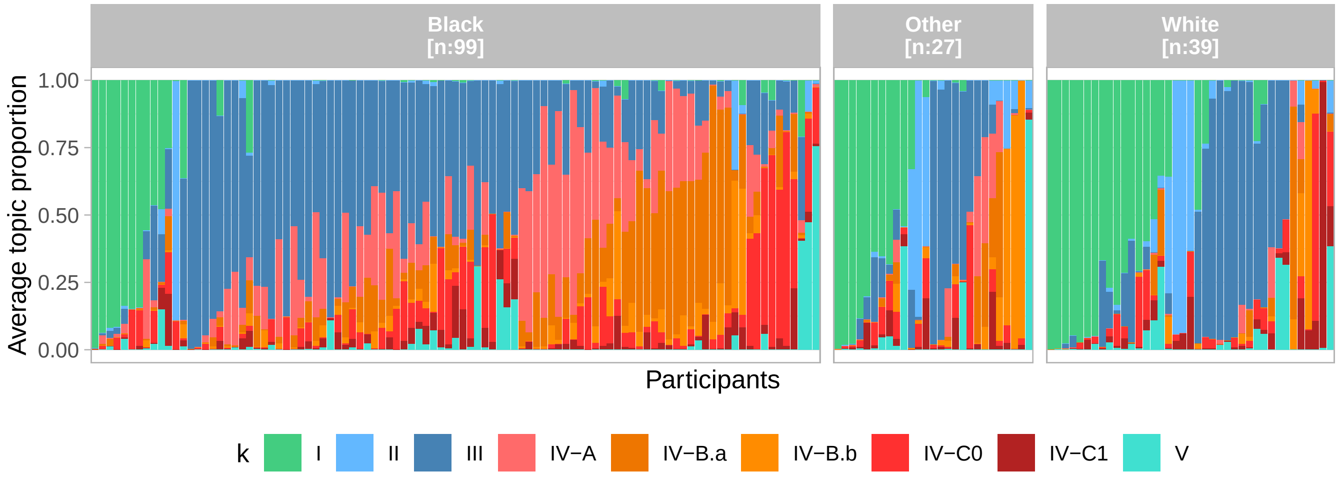

Figure 39: Topic composition per racial groups.

```
g_cohort_composition <-
  plot_topic_composition_per_group_and_subject(
    gammas = gammas,
    colData(mae) %>% as.data.frame() %>%
      select(Subject, SampleID, Status, Site) %>%
      filter(!(Site %in% c("AYAC","EM"))) %>%
      mutate(
        d = SampleID,
        site = case_when(Site == "Stanford" ~ "SU", TRUE ~ Site),
        group = str_c(Status, " (", site,")") %>%
          factor(., levels = c("Non-pregnant (UAB)","Pregnant (UAB)", "Pregnant (SU)")),
        group_name = "Cohort") %>%
      select(Subject, d, group, group_name)
      ) +
  scale_fill_manual(values = get_topic_colors(gammas$k %>% levels()))

g_cohort_composition
```

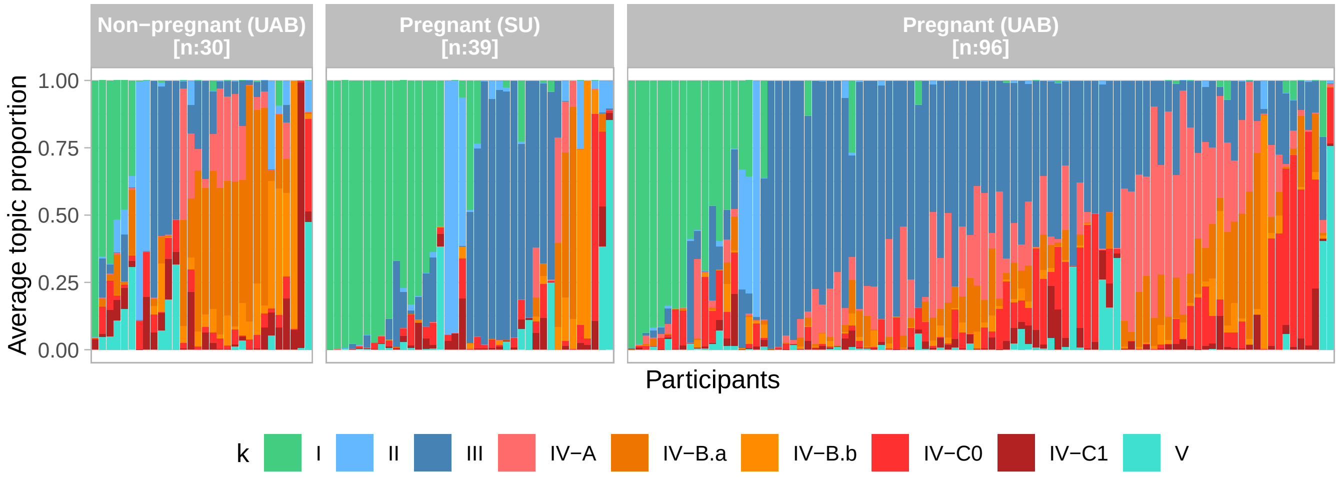

Figure 40: Topic composition per cohort (location and pregnancy status).

From these two figures, it looks like race, location, and pregnancy status might be associated with average topic composition. So, we formally test that with a Dirichlet regression:

```
subject_level_gammas <-
  gammas %>%
  left_join(si, by = "d") %>%
  group_by(Subject, Race, Site, Status, k) %>%
  summarize(prop = mean(g), .groups = "drop")

subject_level_gammas_wide <-
  subject_level_gammas %>%
  pivot_wider(
    id_cols = c(Subject, Race, Site, Status),
    names_from = k, values_from = prop,
    names_prefix = "prop_"
  ) %>%
  ungroup()

library(DirichletReg)

y <- subject_level_gammas_wide %>% select(starts_with("prop_")) %>% as.matrix()
```

```r
x <- subject_level_gammas_wide %>% select(Race, Site, Status) %>% as.data.frame()
data <- x
data$y <- DR_data(y)

dirichlet_model <- DirichReg(y ~ Status, data = data)
dirichlet_model <- DirichReg(y ~ Race, data = data)
dirichlet_model <- DirichReg(y ~ Site, data = data)

dirichlet_model <- DirichReg(y ~ Race + Site + Status, data = data)

dirichlet_model %>% summary()
```

```
## Call:
## DirichReg(formula = y ~ Race + Site + Status, data = data)
##
## Standardized Residuals:
##                  Min       1Q   Median       3Q     Max
## prop_I       -0.8032  -0.5361  -0.5321   1.1932  7.9105
## prop_II      -0.5713  -0.4840  -0.4380  -0.4157  7.5322
## prop_III     -1.4817  -0.6878  -0.0943   1.2140  3.9173
## prop_IV-A    -0.8287  -0.6151  -0.3992   0.4642  4.1813
## prop_IV-B.a  -1.0761  -0.5911  -0.3910   0.0952  5.1388
## prop_IV-B.b  -0.6945  -0.5552  -0.4705  -0.3583  5.2182
## prop_IV-C0   -0.7691  -0.6681  -0.5444  -0.1473  4.7713
## prop_IV-C1   -0.7387  -0.5860  -0.5440  -0.3424  3.8107
## prop_V       -0.5466  -0.4961  -0.4923  -0.3988  6.4213
##
## ------------------------------------------------------------------
## Beta-Coefficients for variable no. 1: prop_I
##                Estimate Std. Error z value Pr(>|z|)
## (Intercept)    -1.46168    0.28648  -5.102 3.36e-07 ***
## RaceBlack      -0.26342    0.25679  -1.026    0.305
## RaceWhite       0.12678    0.25207   0.503    0.615
## SiteSU          0.02972    0.24783   0.120    0.905
## StatusPregnant  0.15615    0.21287   0.734    0.463
## ------------------------------------------------------------------
## Beta-Coefficients for variable no. 2: prop_II
##                Estimate Std. Error z value Pr(>|z|)
## (Intercept)    -1.70715    0.27848  -6.130 8.77e-10 ***
## RaceBlack      -0.16461    0.26003  -0.633    0.527
## RaceWhite       0.10699    0.25018   0.428    0.669
## SiteSU         -0.08927    0.25474  -0.350    0.726
## StatusPregnant -0.08625    0.21253  -0.406    0.685
## ------------------------------------------------------------------
## Beta-Coefficients for variable no. 3: prop_III
##                 Estimate Std. Error z value Pr(>|z|)
## (Intercept)    -1.528291   0.281579  -5.428 5.71e-08 ***
## RaceBlack       0.611719   0.257229   2.378 0.017402 *
## RaceWhite      -0.004043   0.251739  -0.016 0.987188
## SiteSU         -0.883136   0.263496  -3.352 0.000803 ***
## StatusPregnant  1.029248   0.212917   4.834 1.34e-06 ***
## ------------------------------------------------------------------
## Beta-Coefficients for variable no. 4: prop_IV-A
##                Estimate Std. Error z value Pr(>|z|)
## (Intercept)     -1.6882     0.2832  -5.962 2.49e-09 ***
## RaceBlack        0.5192     0.2562   2.026 0.042746 *
## RaceWhite       -0.2380     0.2508  -0.949 0.342748
## SiteSU          -0.9243     0.2434  -3.798 0.000146 ***
```

```
## StatusPregnant   0.3801     0.2074   1.833 0.066770 .
## ------------------------------------------------------------------
## Beta-Coefficients for variable no. 5: prop_IV-B.a
##                Estimate Std. Error z value Pr(>|z|)
## (Intercept)     -1.0829     0.2760  -3.923 8.75e-05 ***
## RaceBlack        0.6904     0.2568   2.689  0.00717 **
## RaceWhite       -0.3829     0.2517  -1.521  0.12819
## SiteSU          -0.3630     0.2495  -1.455  0.14570
## StatusPregnant  -0.5951     0.2087  -2.852  0.00435 **
## ------------------------------------------------------------------
## Beta-Coefficients for variable no. 6: prop_IV-B.b
##                Estimate Std. Error z value Pr(>|z|)
## (Intercept)    -1.22901    0.27040  -4.545 5.49e-06 ***
## RaceBlack       0.09652    0.27109   0.356   0.7218
## RaceWhite      -0.43146    0.25078  -1.720   0.0853 .
## SiteSU         -0.35627    0.26466  -1.346   0.1782
## StatusPregnant -0.28238    0.22062  -1.280   0.2006
## ------------------------------------------------------------------
## Beta-Coefficients for variable no. 7: prop_IV-C0
##                Estimate Std. Error z value Pr(>|z|)
## (Intercept)    -0.99797    0.27566  -3.620 0.000294 ***
## RaceBlack       0.07146    0.24598   0.291 0.771429
## RaceWhite      -0.07510    0.24993  -0.300 0.763814
## SiteSU         -0.28344    0.24016  -1.180 0.237909
## StatusPregnant -0.17831    0.20772  -0.858 0.390661
## ------------------------------------------------------------------
## Beta-Coefficients for variable no. 8: prop_IV-C1
##                Estimate Std. Error z value Pr(>|z|)
## (Intercept)    -1.01197    0.27830  -3.636 0.000277 ***
## RaceBlack      -0.01188    0.24912  -0.048 0.961979
## RaceWhite       0.02633    0.24887   0.106 0.915751
## SiteSU         -0.12040    0.24696  -0.488 0.625870
## StatusPregnant -0.39807    0.20484  -1.943 0.051979 .
## ------------------------------------------------------------------
## Beta-Coefficients for variable no. 9: prop_V
##                 Estimate Std. Error z value Pr(>|z|)
## (Intercept)    -1.690073   0.278904  -6.060 1.36e-09 ***
## RaceBlack      -0.034231   0.254961  -0.134    0.893
## RaceWhite       0.009452   0.251894   0.038    0.970
## SiteSU         -0.285821   0.260729  -1.096    0.273
## StatusPregnant  0.001159   0.212154   0.005    0.996
## ------------------------------------------------------------------
## Significance codes: 0 '***' 0.001 '**' 0.01 '*' 0.05 '.' 0.1 ' ' 1
##
## Log-likelihood: 4420 on 45 df (135 BFGS + 2 NR Iterations)
## AIC: -8751, BIC: -8611
## Number of Observations: 165
## Link: Log
## Parametrization: common
```

```r
save(dirichlet_model, file = "../results/topics_and_demographic_variables_associations.Rdata")
```

```r
g_dem_associations <- plot_dirichlet_model_results(dirichlet_model)
g_dem_associations
```

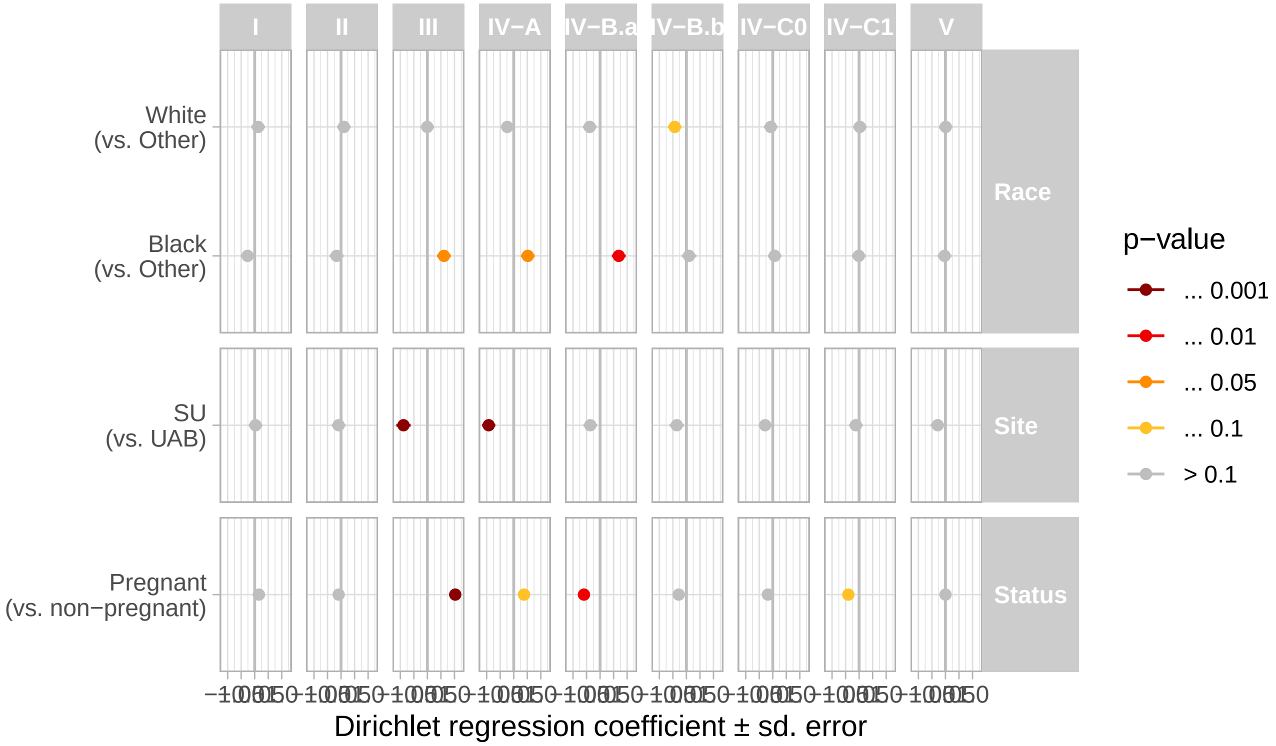

Note that the dirichlet/multinomial approach is better than performing multiple logistic regression (one for each topic) as the latter approach would miss some key differences in topic composition such as a larger proportion of topic III in black participants. That is because the few white participants with topic III have larger proportion of topic III, leading to an roughly similar average proportion of topic III in both groups.

The figures and test below show the results with this multiple logistic regression approach.

```
g_status <-
  plot_topic_distribution_per_group_and_subject(
    gammas = gammas,
    groups = colData(mae) %>%
      as.data.frame() %>%
      select(Subject, SampleID, Status) %>%
      dplyr::rename(d = SampleID,
                    group = Status) %>%
      mutate(group_name = "Status") %>%
      mutate(group = group %>% factor(., levels = c("Non−pregnant","Pregnant"))),
    add_test_significance = TRUE,
    add_mean = TRUE
  )

g_status
```

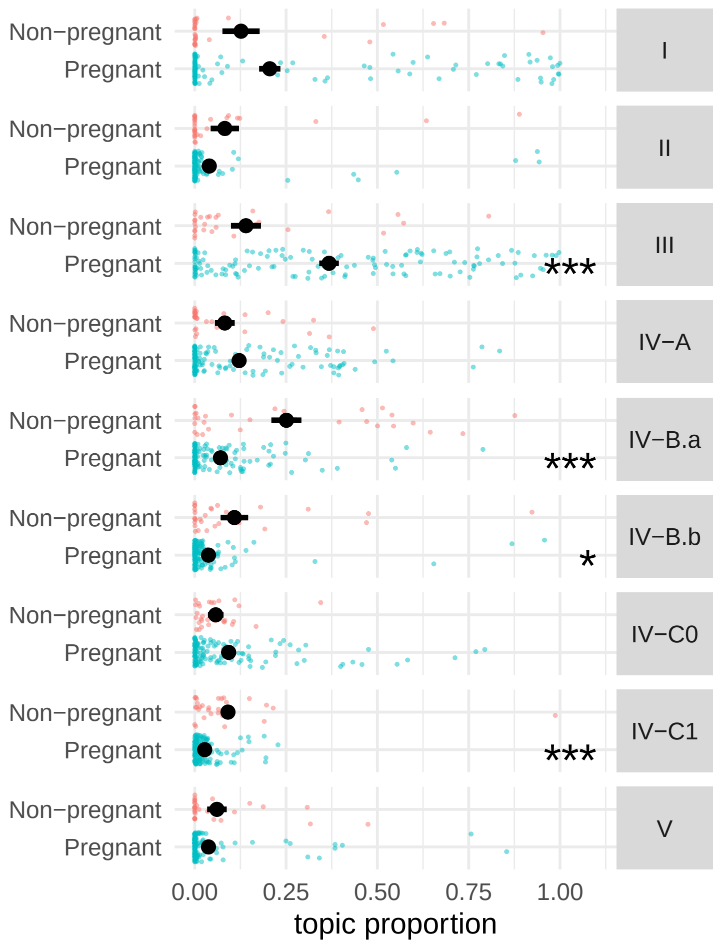

Figure 41: Topic distribution per status in all subjects.

```
g_cohorts <-
  plot_topic_distribution_per_group_and_subject(
    gammas = gammas,
    groups =
      colData(mae) %>% as.data.frame() %>%
      select(Subject, SampleID, Status, Site) %>%
      filter(!(Site %in% c("AYAC","EM"))) %>%
      mutate(
        d = SampleID,
        site = case_when(Site == "Stanford" ~ "SU", TRUE ~ Site),
        group = str_c(Status, " (", site,")") %>%
          factor(., levels = c("Non-pregnant (UAB)","Pregnant (UAB)", "Pregnant (SU)")),
        group_name = "Cohort") %>%
      select(Subject, d, group, group_name),
    add_test_significance = TRUE,
    add_mean = TRUE
  )

g_cohorts
```

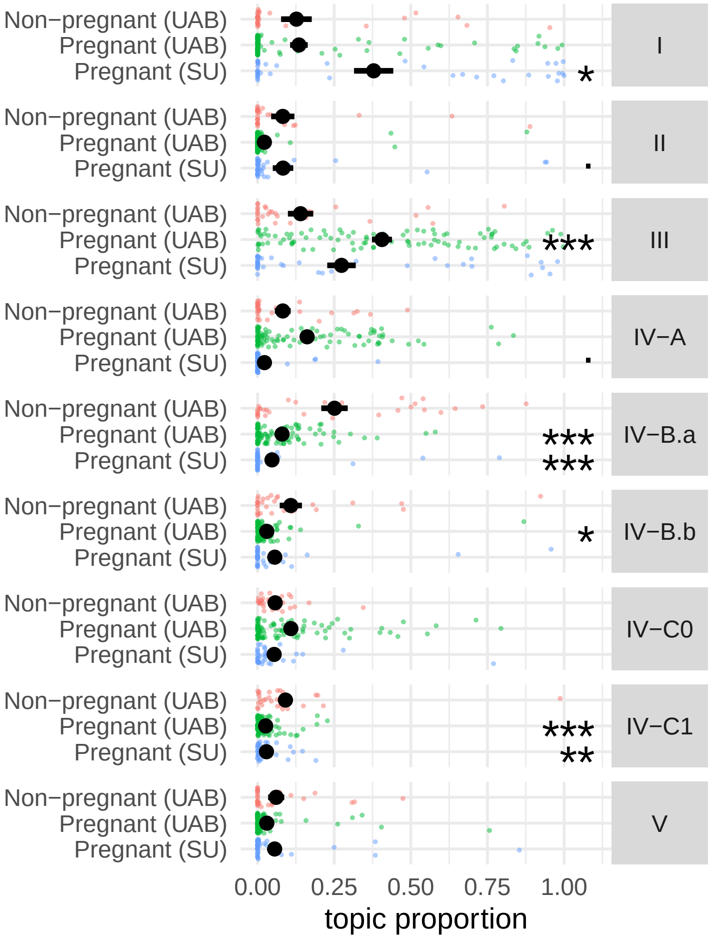

Figure 42: Topic distribution per status in all subjects.

```
g_race <-
  plot_topic_distribution_per_group_and_subject(
    gammas = gammas,
    groups = colData(mae) %>% as.data.frame() %>%
      select(Subject, SampleID, Race) %>%
      mutate(Race = ifelse(Race %in% c("Asian","Hispanic/Latino"), "Other", Race)) %>% # there are not enough
      dplyr::rename(d = SampleID,
                    group = Race) %>%
      mutate(group_name = "Race",
             group = group %>% factor(., levels = c("Other","Black","White"))),
    add_test_significance = TRUE,
    add_mean = TRUE
  )

g_race
```

Table 16: Variables with statistically significant associations with topic proportions.

| k | variable | estimate | str_error | z_value | p_value | q_value | sign_level |
|---|---|---|---|---|---|---|---|
| I | RaceBlack | -1.126328 | 0.5354594 | -2.103480 | 0.0369879 | 0.0979092 | . |
| II | RaceBlack | -1.772012 | 0.7468570 | -2.372626 | 0.0188501 | 0.0605895 | . |
| III | StatusPregnant | 1.392470 | 0.3729638 | 3.733526 | 0.0002621 | 0.0019655 | ** |
| IV-B.a | StatusPregnant | -1.630211 | 0.3214026 | -5.072180 | 0.0000011 | 0.0000162 | *** |
| IV-B.b | RaceBlack | -1.324153 | 0.5057425 | -2.618236 | 0.0096874 | 0.0396301 | * |
| IV-B.b | RaceWhite | -1.244130 | 0.5377842 | -2.313437 | 0.0219714 | 0.0659143 | . |
| IV-B.b | StatusPregnant | -1.345966 | 0.4497504 | -2.992696 | 0.0032040 | 0.0144182 | * |
| IV-C1 | StatusPregnant | -1.145439 | 0.3178900 | -3.603255 | 0.0004191 | 0.0026945 | ** |

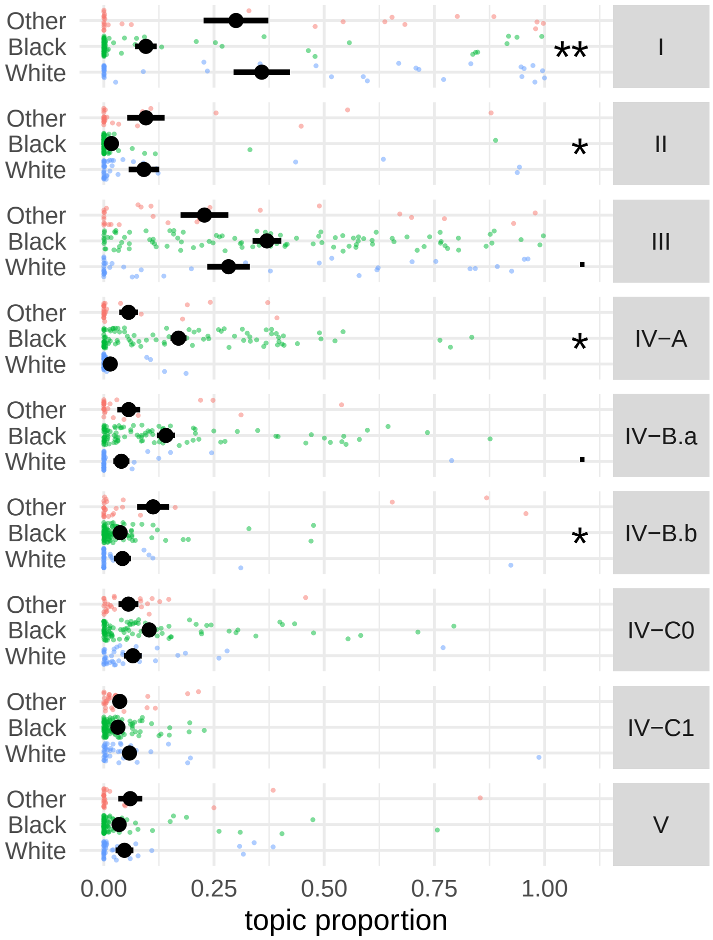

Figure 43: Topic distribution per Race in all subjects.

```r
# One approach is to fit a model for each topic

test_res <- test_for_groups_differences(gammas, si, groups = c("Race", "Site","Status"))

test_res %>%
  filter(variable != "(Intercept)", q_value <= 0.1) %>%
  kable(
    ., booktab = TRUE, format = "latex",
    caption = "Variables with statistically significant associations with topic proportions."
  )
```

Table 17: Associations between the topics and the menstrual cycle.

| k | amplitude | phase_min | phase_max | pval | sign_level |
|---|---|---|---|---|---|
| I | 0.3102697 | 3 | -16 | 0.0041534 | ** |
| II | 0.1395453 | 1 | -14 | 0.2345929 | |
| III | 0.0971368 | -14 | -13 | 0.3185800 | |
| IV-A | 0.0762947 | 7 | -8 | 0.1045411 | |
| IV-B.a | 0.0723358 | -4 | 1 | 0.5239866 | |
| IV-B.b | 0.0541842 | -4 | 2 | 0.5937547 | |
| IV-C0 | 0.1076709 | -9 | 2 | 0.0000000 | *** |
| IV-C1 | 0.1250077 | -12 | 1 | 0.0071655 | ** |
| V | 0.0657596 | 1 | -1 | 0.7011065 | |

##### 4.4.2 Topic proportion throughout the cycle

```r
mc_data <-
  colData(mae) %>%
  as.data.frame() %>%
  filter(Status != "Pregnant", !is.na(cycleday), MC_ok) %>%
  dplyr::select(SampleID, Subject, cycle_nb_m, cycleday) %>%
  mutate(d = SampleID)

df <-
  gammas %>%
  inner_join(mc_data, by = "d")

df_viz <-
   df %>%
  group_by(Subject, cycle_nb_m, k) %>%
  mutate(has_k = any(g > 0.05)) %>%
  ungroup() %>%
  filter(has_k)

save(df_viz, file = "../results/df_viz_topics_around_MC")
```

We test if any of the topics are associated with the menstrual cycle.

```r
test_for_MC_association_topic_proportion(df_viz, df = 4) %>%
  kable(
    ., booktab = TRUE, format = "latex",
    caption = "Associations between the topics and the menstrual cycle."
  )
```

```r
g_cycle <- plot_topics_distribution_around_MC(df_viz)
g_cycle
```

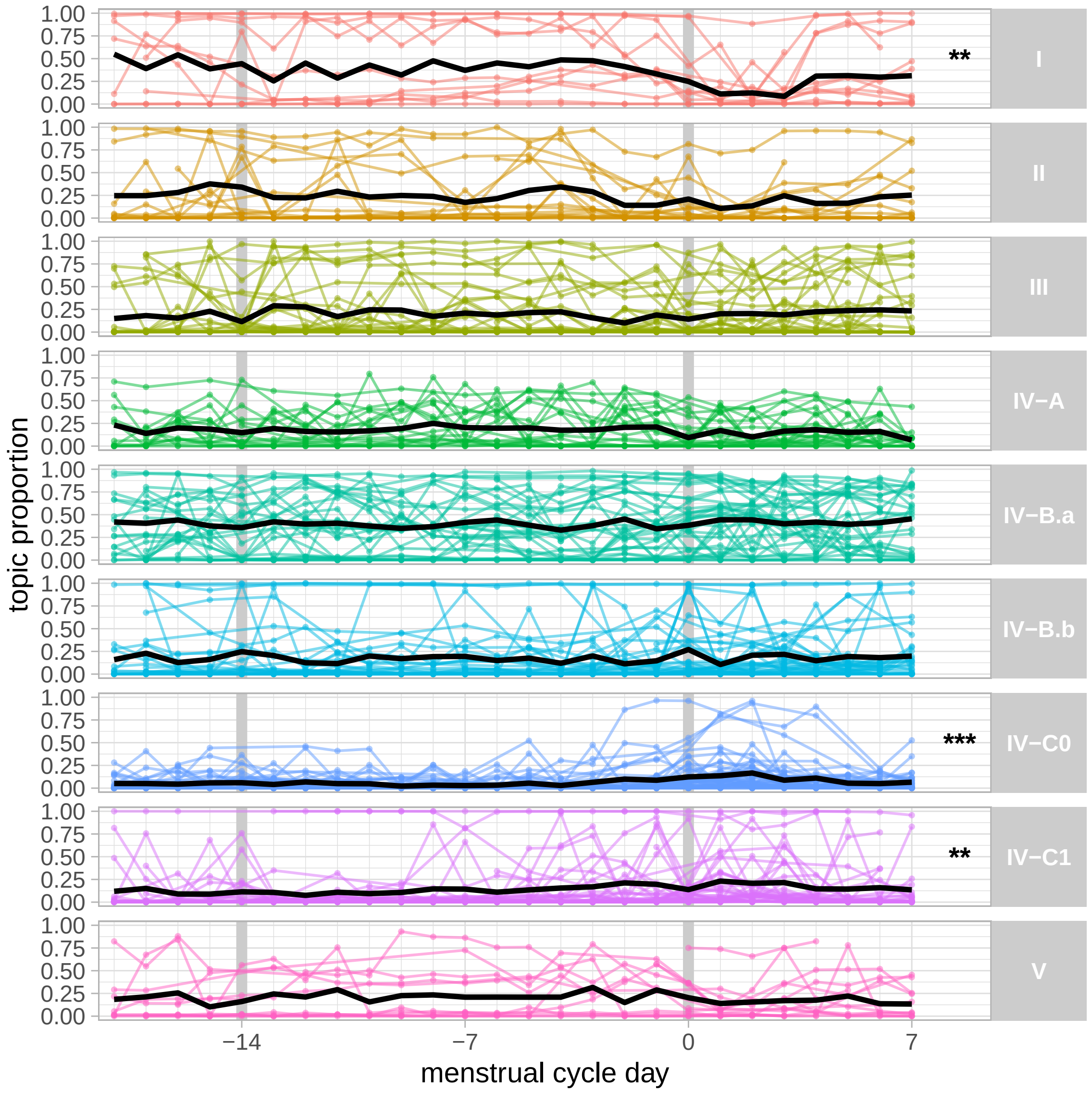

##### 4.4.3 Association topic proportions and preterm birth

```
preterm_si <-
  colData(mae) %>%
  as.data.frame() %>%
  filter(Status == "Pregnant", GestationalAge_days < GestationalAgeAtDelivery_days) %>%
  dplyr::select(SampleID, Subject, GestationalAgeAtDelivery_days) %>%
  mutate(
    preterm =
      GestationalAgeAtDelivery_days %>%
      cut(., breaks = c(-Inf, 7*37, Inf), labels = c("Preterm","Term"))
  )

g_preterm <-
  plot_topic_and_preterm_association(
    topic_and_preterm_association(
      gammas = gammas,
      preterm = preterm_si
    )
  )
```

```
g_preterm
```

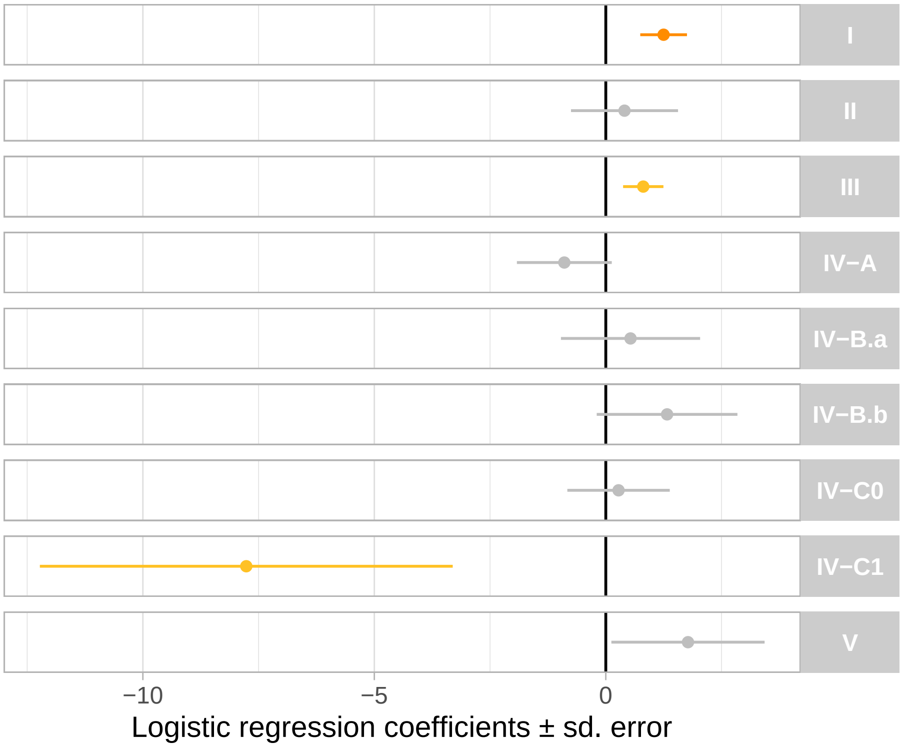

```
g_preterm_2 <-
  plot_topic_distribution_per_group_and_subject(
    gammas = gammas,
    groups =
      colData(mae) %>%
      as.data.frame() %>%
      filter(Status == "Pregnant", GestationalAge_days < GestationalAgeAtDelivery_days) %>%
      dplyr::select(SampleID, Subject, GestationalAgeAtDelivery_days) %>%
      mutate(
        preterm = GestationalAgeAtDelivery_days %>% cut(., breaks = c(-Inf, 7*37, Inf), labels = c("Preterm",
        group_name = "Delivery"
        ) %>%
      dplyr::rename(
        d = SampleID,
        group = preterm
      ),
    add_test_significance = TRUE,
    add_mean = TRUE
  )

g_preterm_2
```

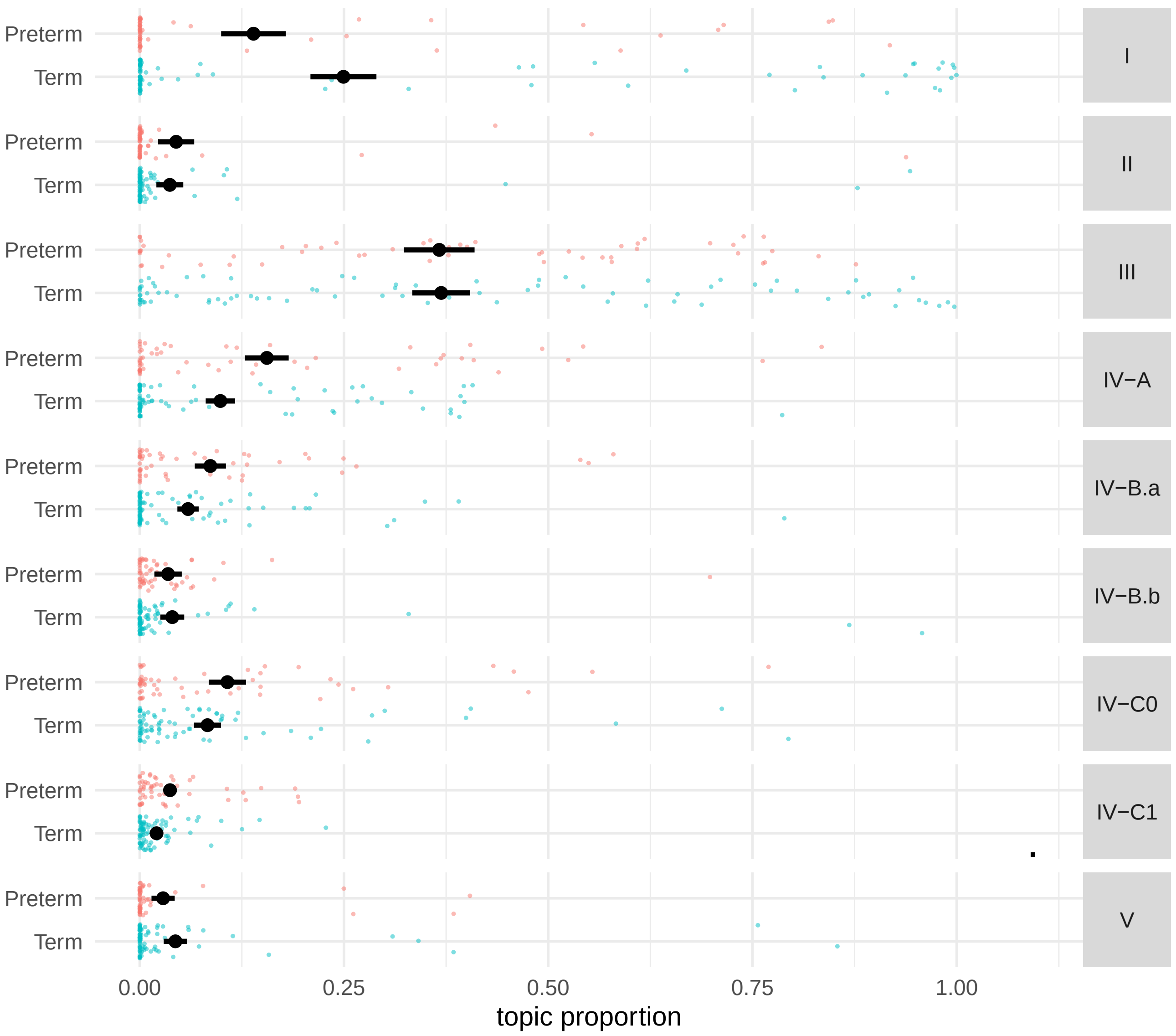

```
topic_and_short_term_preterm_risk_association <- function(gammas, preterm) {

  topic_names <-
    tibble(topic_name = gammas$k %>% levels()) %>%
    mutate(k = row_number())
  df <-
    gammas %>%
    dplyr::rename(topic_name = k) %>%
    mutate(k = topic_name %>% as.integer()) %>%
    pivot_wider(id_cols = d, names_from = k, values_from = g, names_prefix = "k_") %>%
    left_join(preterm %>% select(SampleID, preterm_risk_4w) %>% dplyr::rename(d = SampleID) %>% distinct(), b
    mutate(y = scale(preterm_risk_4w))
  formula <- str_c("y ~ ", str_c("k_", topic_names$k) %>% str_c(., collapse = " + "), " + 0")
  summ <- glm(formula, data = df, family = "gaussian") %>%  summary()
  summ$coefficients %>% as_tibble() %>%
    mutate(k = topic_names$topic_name) %>%
    dplyr::select(k, everything()) %>%
    mutate(p_val_cat = get_sign_levels(`Pr(>|t|)`))
}

short_term_preterm_assoc <-
  topic_and_short_term_preterm_risk_association(
    gammas = gammas,
```

```r
    preterm =
      colData(mae) %>%
      as.data.frame() %>%
      filter(Status == "Pregnant", GestationalAge_days < GestationalAgeAtDelivery_days) %>%
      dplyr::select(SampleID, Subject, GestationalAgeAtDelivery_days, preterm_risk_4w)
  )

g_short_preterm <-
  ggplot(short_term_preterm_assoc %>% mutate(k = k)) +
  geom_vline(xintercept = 0) +
  geom_segment(aes(x = Estimate - `Std. Error`, xend = Estimate + `Std. Error`,
                   y = k, yend = k, col = p_val_cat)) +
  geom_point(aes(x = Estimate, y = k, col = p_val_cat)) +
  scale_color_manual("p-value",
                     breaks = p_val_labels,
                     values = p_val_cols) +
  facet_grid(k ~ ., scales = "free") +
  xlab("Logistic regression coefficients ± sd. error") +
  ylab("") +
  scale_y_discrete(breaks = NULL) +
  theme(
    legend.position = "top",
    strip.text.y = element_text(angle = 0, hjust = 0.5),
    strip.background.y = element_rect(fill = "gray80", color = NA)
  ) +
  guides(col = "none")

g_short_preterm
```

### 5 Topic time-series

```
mae = readRDS("../results/mae_for_analyses.Rds")
load("../results/gammas.Rdata")
```

```
selected_subjects = c("40003", "40056")

for(s in selected_subjects){
  plot_subject_topic_time_series(
    mae = mae, gammas = gammas, subj = s,
    xlab = "Gestational age (weeks)"
  ) %>%
    print()
}
```

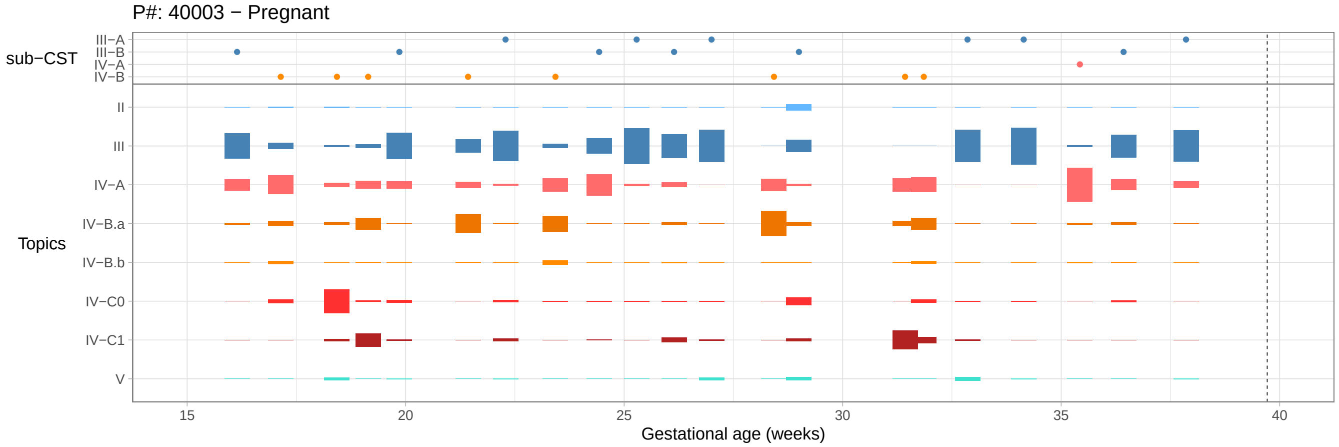

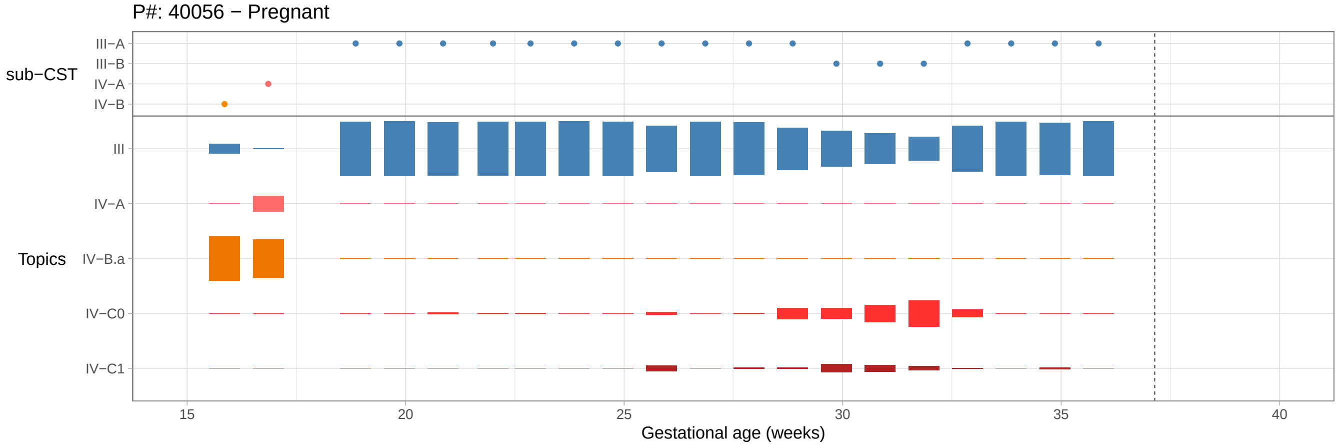

```
selected_subjects = c("UAB022", "UAB028", "UAB077") # "UAB077" "UAB044"  "UAB006" "UAB010" #,"AYAC05")

for(s in selected_subjects){
   plot_subject_topic_time_series(mae = mae, gammas = gammas, subj = s,
                       xlab = "Time since study start (weeks)",
                        xticks = seq(0,10,by = 2)) %>%
    print()
}
```

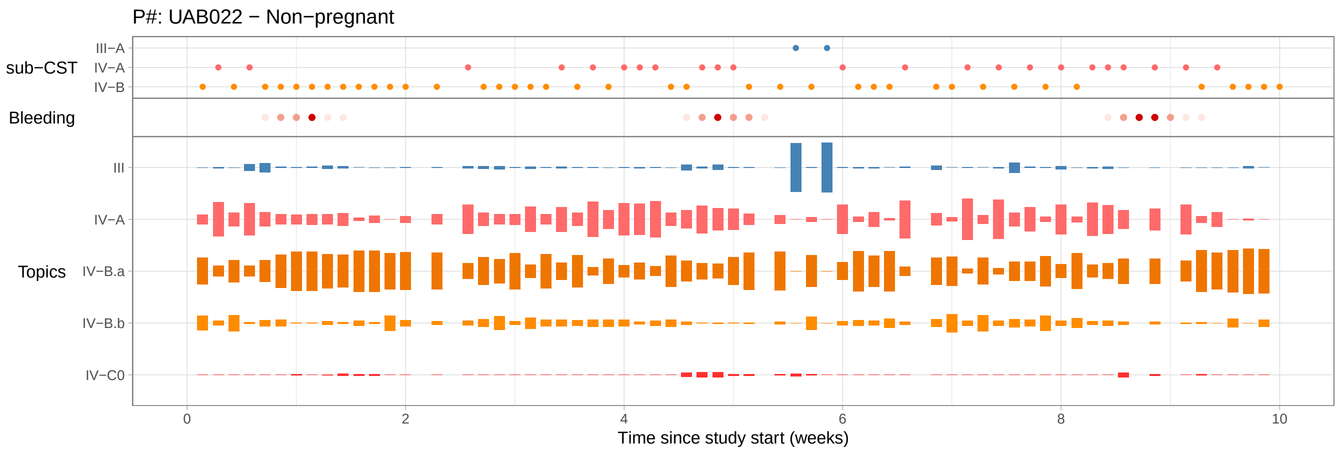
P#: UAB022 − Non−pregnant
sub−CST
III−A
IV−A
IV−B
Bleeding
Topics
III
IV−A
IV−B.a
IV−B.b
IV−C0
0
2
4
6
8
10
Time since study start (weeks)

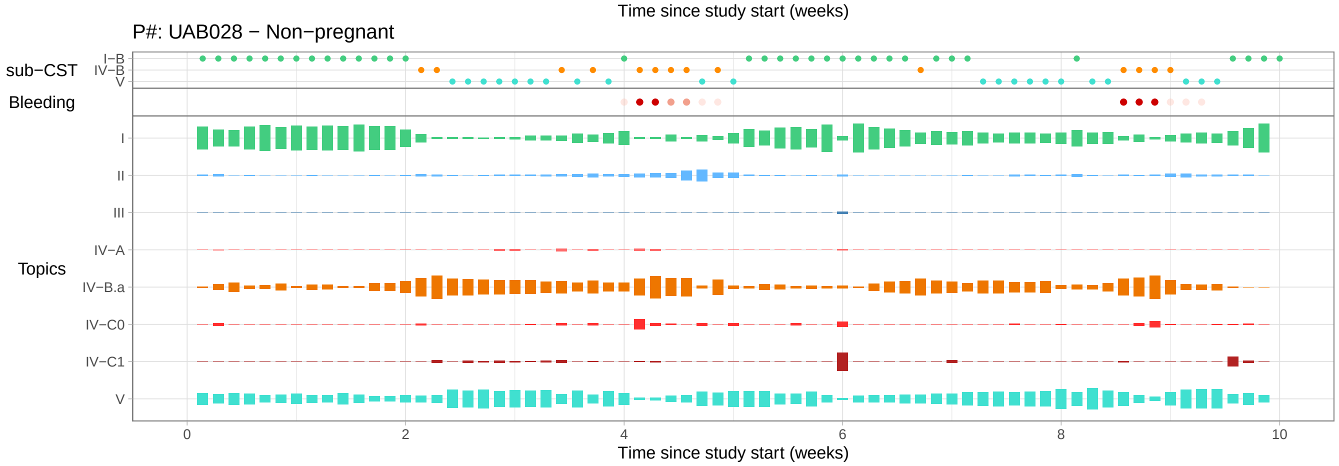
P#: UAB028 − Non−pregnant
sub−CST
I−B
IV−B
V
Bleeding
Topics
I
II
III
IV−A
IV−B.a
IV−C0
IV−C1
V
0
2
4
6
8
10
Time since study start (weeks)

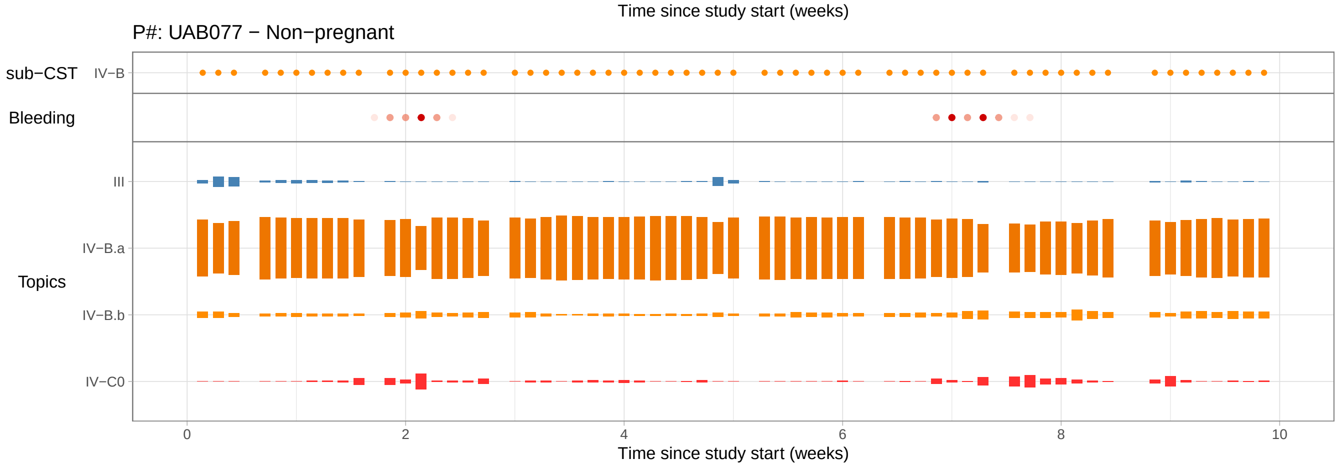
P#: UAB077 − Non−pregnant
sub−CST
IV−B
Bleeding
Topics
III
IV−B.a
IV−B.b
IV−C0
0
2
4
6
8
10
Time since study start (weeks)

### 6 Vaginal microbiome composition throughout the menstrual cycle

#### 6.1 Microbiota composition throughout the menstrual cycle.

We previously observed that three topics had decreased or increased proportions throughout the menstrual cycle. Here, we investigate if there are further associations with the menstrual cycle. Specifically, we look at the intra-individual between-cycle correlations. In other words, we are interested in understand if participants that have unstable microbiota composition have similar changes in composition from one cycle to the next.

```
mae = readRDS("../results/mae_for_analyses.Rds")
load("../results/gammas.Rdata")
```

```
si <- colData(mae) %>% as.data.frame()
number_of_cycles <-
  si %>%
  filter(SampleID %in% gammas$d) %>%
  filter(!is.na(cycleday)) %>%
  group_by(Subject, cycle_nb_m) %>%
  summarize(n_days = n(), range_cycleday = range(cycleday) %>% diff, .groups = "drop") %>%
  filter(n_days >= 10, range_cycleday >= 18) %>%
  select(Subject, cycle_nb_m) %>%
  distinct() %>%
  group_by(Subject) %>%
  summarize(n_cycles = n())
```

Number of participants with at least one menstrual cycle: 26.

Number of participants with at least two menstrual cycles: 20.

We add the cycle data (cycle number, cycleday, etc) to the topic composition data.

```
df_topics <-
  gammas %>%
  dplyr::rename(SampleID = d, feature = k, prop = g) %>%
  left_join(colData(mae) %>% as.data.frame(), by = "SampleID")
```

We filter for participants and samples that belong to two consecutive menstrual cycles.

```
df_topics_consecutive_cycles <-
  get_df_consecutive_cycles(df_topics)

df_topics_consecutive_cycles <-
  df_topics_consecutive_cycles %>%
  group_by(Subject, feature) %>%
  mutate(median_prop = median(prop), max_prop = max(prop)) %>%
  ungroup() %>%
  filter(max_prop > 0.05, median_prop > 0.01)
```

We compute the correlation between the topic composition of two consecutive menstrual cycles.

```
correlations_between_consecutive_cycles <-
  compute_correlation_between_cycles(df_topics_consecutive_cycles)

n_sign_topics <- (correlations_between_consecutive_cycles$qvalue <= 0.05) %>% sum()
n <- nrow(correlations_between_consecutive_cycles)

# n_sign_topics
# n_above_half/n
```

20 (4) of participants have a significant between-cycle correlation.

```
plot_correlations_between_consecutive_cycles(correlations_between_consecutive_cycles)
```

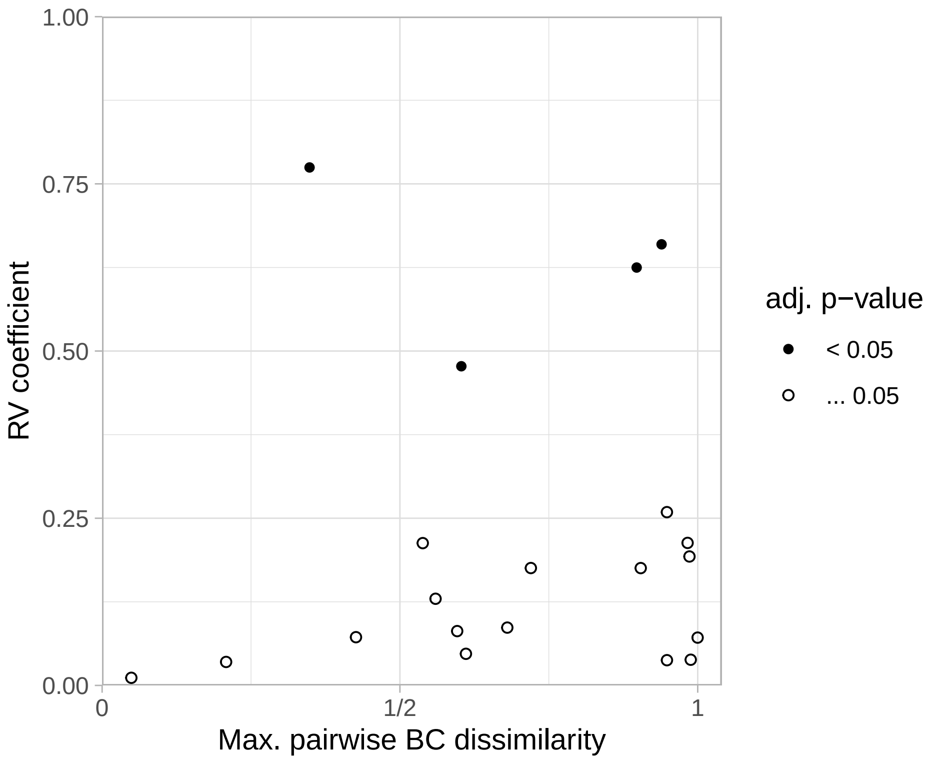

```
plot_composition_consecutive_cycles(df_topics_consecutive_cycles)
```

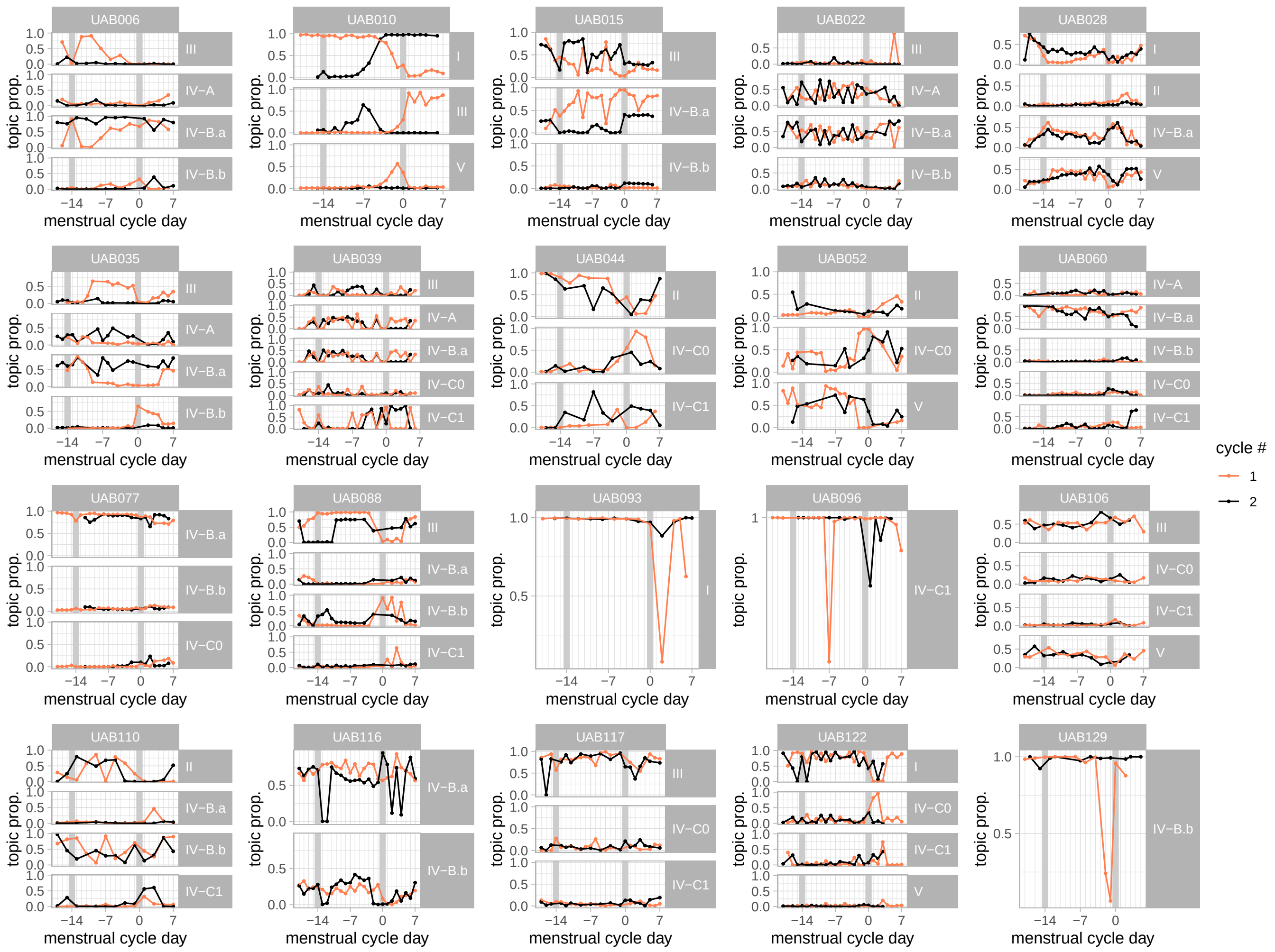

We also compute the correlationa and visualize the taxa composition of consecutive cycles:

```r
df_taxa <- get_taxa_proportion_long_format(mae, "VM16S_combined")

df_taxa_consecutive_cycles <-
  get_df_consecutive_cycles(df_taxa)

df_taxa_consecutive_cycles <-
  df_taxa_consecutive_cycles %>%
  group_by(Subject, feature) %>%
  mutate(median_prop = median(prop), max_prop = max(prop)) %>%
  ungroup() %>%
  filter(max_prop > 0.05, median_prop > 0.01)

# plot_composition_consecutive_cycles(df_taxa_consecutive_cycles)
```

```r
plot_composition_consecutive_cycles(
  df_taxa_consecutive_cycles %>%
    filter(Subject %in% c("UAB077","UAB028"))
  )
```

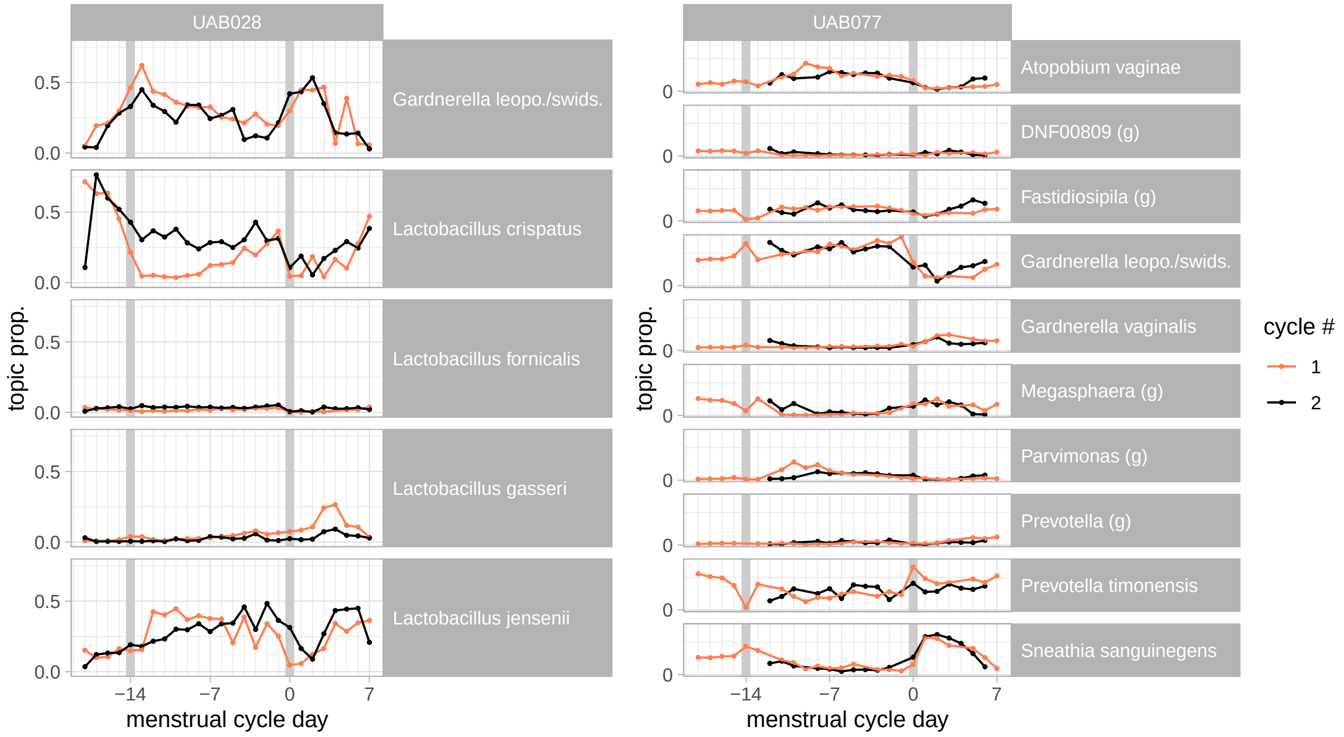

```r
correlations_between_consecutive_cycles_taxa_level <-
  compute_correlation_between_cycles(df_taxa_consecutive_cycles)

plot_correlations_between_consecutive_cycles(correlations_between_consecutive_cycles_taxa_level)
```

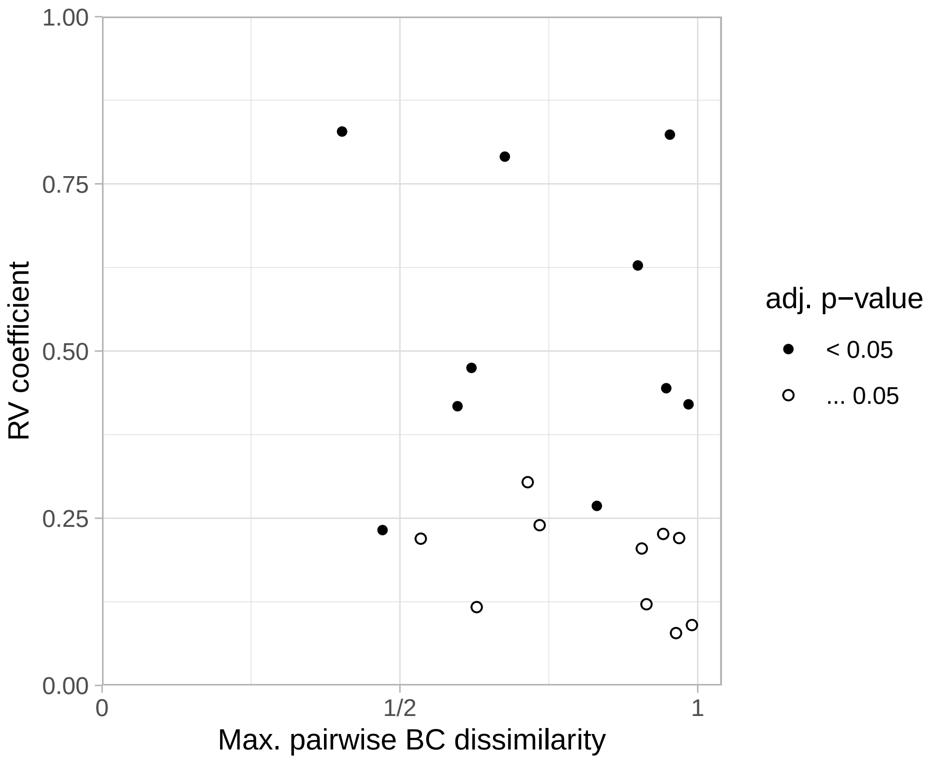

```
n_sign_taxa <- (correlations_between_consecutive_cycles_taxa_level$qvalue <= 0.05) %>% sum()

# n_sign_taxa
# n_sign_taxa/n
```

50 (10) of participants have a significant between-cycle correlation when microbiota is described by taxa proportions.

```
save(df_topics_consecutive_cycles, file = "../results/data_for_figures/df_topics_consecutive_cycles.Rdata")
save(df_taxa_consecutive_cycles, file = "../results/data_for_figures/df_taxa_consecutive_cycles.Rdata")
```

#### 6.2 Metabolites

In this section, we identify the metabolites which have differential abundance at specific phases of the menstrual cycle.

We do this by fitting the abundances to a periodic spline along the standardized menstrual cycle days.

```
MB_long <- get_assays_long_format(mae, assay_name = "MB_NP_t", imputed_assay_name = "MB_NP_t_imputed")

MB_MC_associations = compute_MC_associations(MB_long)

MB_MC_associations %>%
  arrange(association_strength) %>%
  group_by(association_strength) %>%
  summarize(n = n(), .groups = "drop") %>%
  kable(., format = "latex", booktab = TRUE,
        cap = "Number of metabolites with an association with the menstrual cycle.") %>%
  kableExtra::kable_styling(latex_options = "HOLD_position")
```

Table 18: Number of metabolites with an association with the menstrual cycle.

| association_strength | n |
|---|---|
| no association | 276 |
| association | 32 |
| strong association | 28 |

```r
metabolites_with_MC_variations <-
  MB_MC_associations %>%
  filter(association_strength != "no association") %>%
  arrange(-effect)

metabolites_with_strong_MC_variations <-
  MB_MC_associations %>%
  filter(association_strength == "strong association") %>%
  arrange(-effect)

selected_metabolites <-
  MB_MC_associations %>%
  arrange(qval) %>%
  slice_head(n = 6) %>%
  select(feature, pval, qval, effect)
```

```r
g_MB_MC <-
  plot_MC_associations_heatmap(
  MB_long %>%
    filter(feature %in% metabolites_with_MC_variations$feature)
)

save(g_MB_MC, file = "../results/suppl_figs/MB_MC.Rdata")

g_MB_MC
```

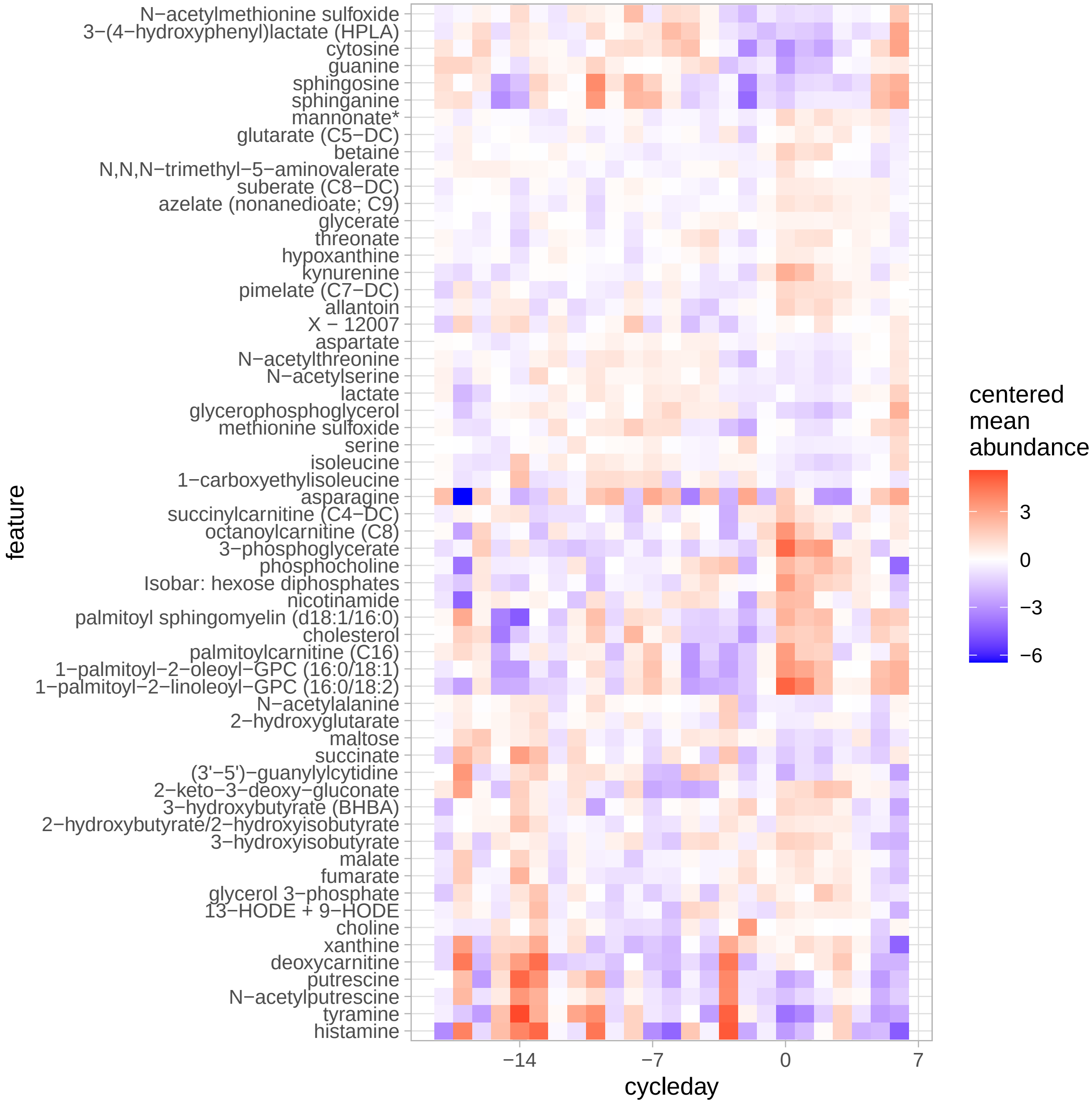

Figure 44: Metabolites associated with the menstrual cycle.

Most metabolites peak or drop around menses:

```
MB_MC_associations_long <-
  MB_MC_associations %>%
  filter(association_strength != "no association") %>%
  pivot_longer(cols = starts_with("phase"), values_to = "phase", names_to = "direction") %>%
  mutate(
    cycle_phase =
      cut(phase, breaks = c(-19, -16, -12, -3, 5, 7),
          labels = c("follicular","peri-ovulatory","luteal", "menses","follicular"))
  )

# ggplot(MB_MC_associations_long,
#        aes(x = phase, fill = direction)) +
```

```r
#   geom_histogram(binwidth = 1) +
#   scale_x_continuous(breaks = -21:10)

ggplot(
  MB_MC_associations_long,
  aes(x = cycle_phase, fill = direction)
) +
  geom_bar()
```

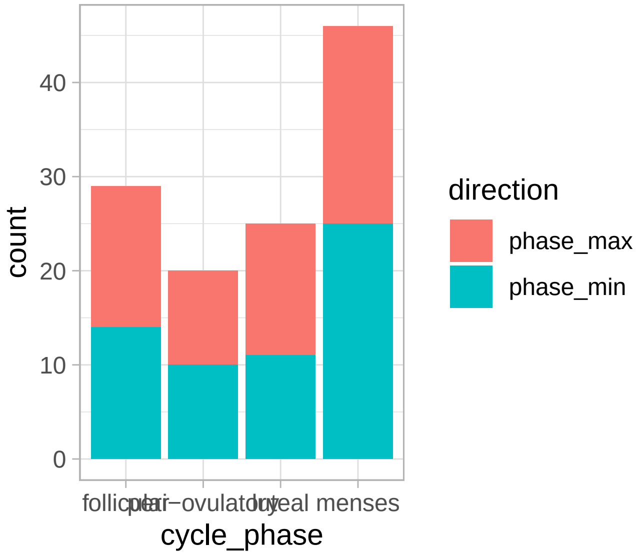

```r
plot_MC_associations(
  MB_long %>% filter(feature %in% metabolites_with_strong_MC_variations$feature),
  ncol = 4
)
```

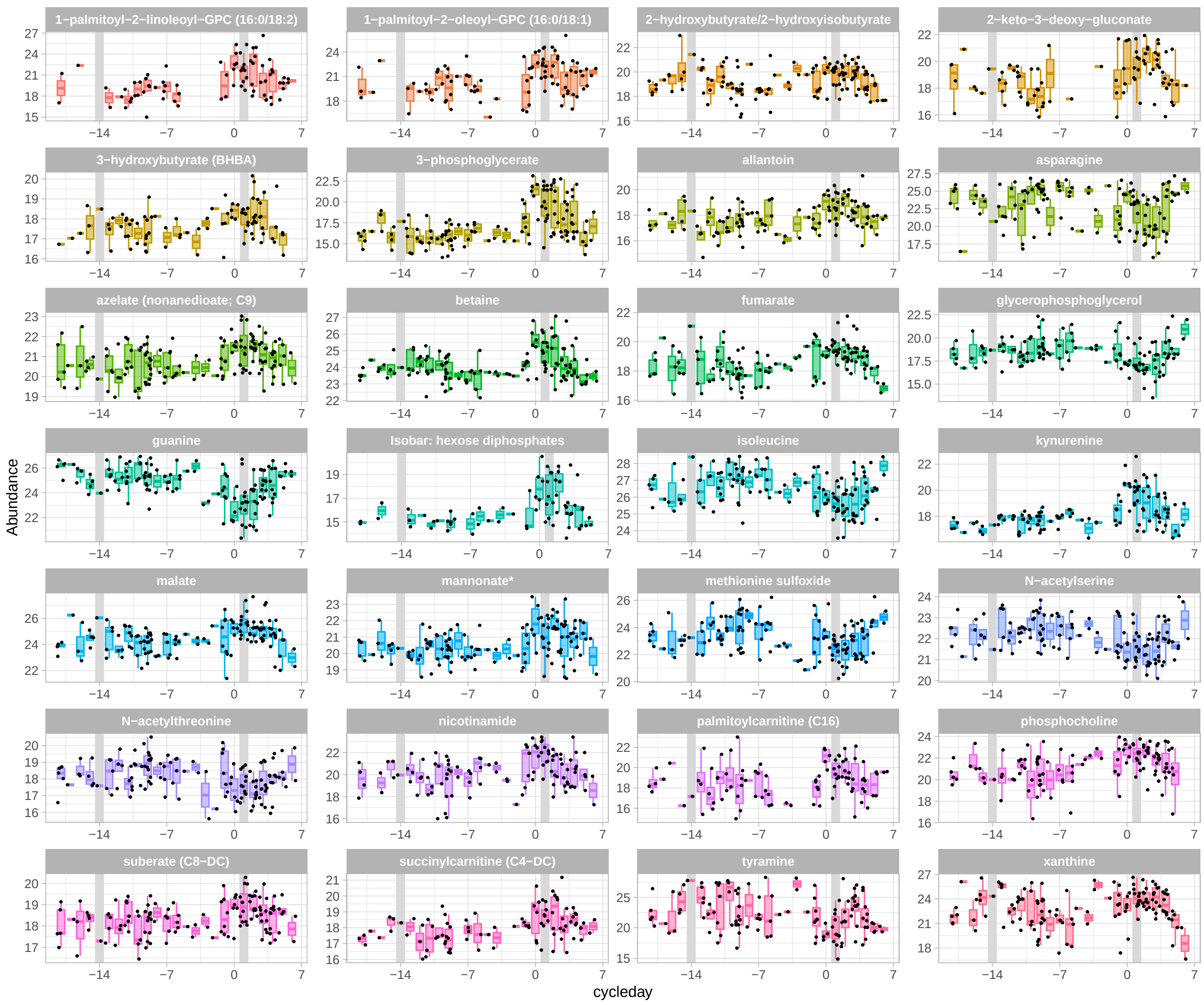

Figure 45: Metabolites with a strong association with the menstrual cycle.

```r
df = MB_long %>%
  filter(feature %in% selected_metabolites$feature)

save(df, file = str_c(fig_data_dir, "selected_MC_metabolites.Rdata"))
```

#### 6.3 Cytokines

We proceed similarly for the cytokines.

```r
I <- assay(mae, "I_t_imputed") %>% t()

I_long <- get_assays_long_format(mae, assay_name = "I_t", imputed_assay_name = "I_t_imputed")
I_MC_associations <- compute_MC_associations(I_long)
cycling_cytokines <- I_MC_associations %>% filter(association_strength != "no association")

g_I_MC <-
  plot_MC_associations(
  I_long %>%
    mutate(abundance = 10^abundance) %>%
    filter(feature %in% cycling_cytokines$feature),
```

```
  ncol = 4
) +
  scale_y_log10("concentration (pg/mL)")

save(g_I_MC, file = "../results/suppl_figs/I_MC.Rdata")

g_I_MC
```

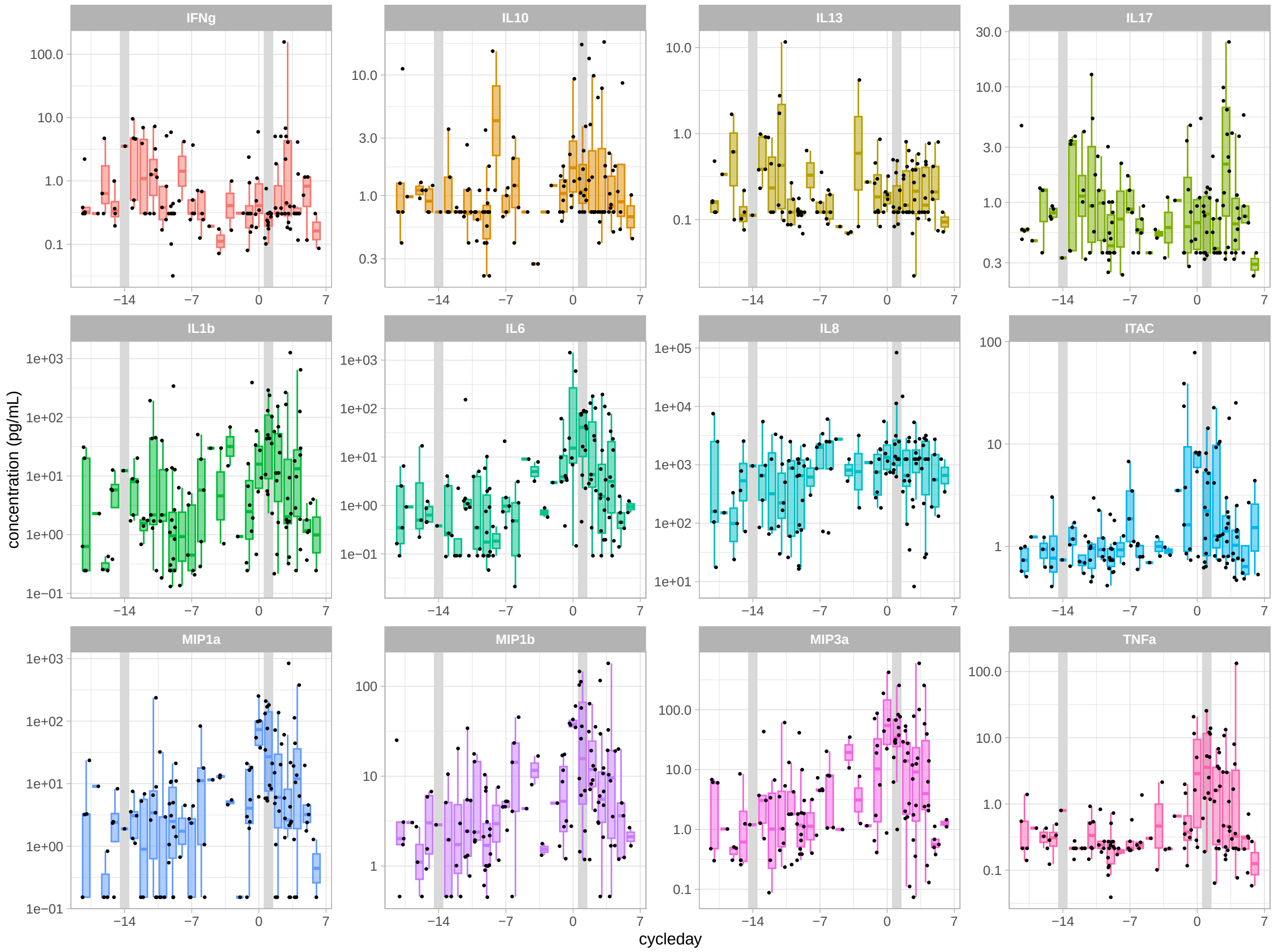

Figure 46: Cytokines whose concentration varies with the menstrual cycle.

#### Reproducibility Receipt

```r
session_info()
```

```
## - Session info ---------------------------------------------------------------
##  setting  value
##  version  R version 4.2.1 (2022-06-23)
##  os       macOS Big Sur ... 10.16
##  system   x86_64, darwin17.0
##  ui       X11
##  language (EN)
##  collate  en_US.UTF-8
##  ctype    en_US.UTF-8
##  tz       America/Los_Angeles
##  date     2022-12-19
##  pandoc   2.19.2 @ /Applications/RStudio.app/Contents/Resources/app/quarto/bin/tools/ (via rmarkdown)
##
## - Packages -------------------------------------------------------------------
##  ! package              * version   date (UTC) lib source
##  P abind                  1.4-5     2016-07-21 [?] CRAN (R 4.2.0)
##    ade4                   1.7-20    2022-11-01 [1] CRAN (R 4.2.0)
##    affy                   1.76.0    2022-11-01 [1] Bioconductor
##    affyio                 1.68.0    2022-11-01 [1] Bioconductor
##    alto                 * 0.1.0     2022-11-17 [1] local
##    annotate               1.76.0    2022-11-01 [1] Bioconductor
##    AnnotationDbi          1.60.0    2022-11-01 [1] Bioconductor
##    ape                    5.6-2     2022-03-02 [1] CRAN (R 4.2.0)
##    assertthat             0.2.1     2019-03-21 [3] CRAN (R 4.2.0)
##  P backports              1.4.1     2021-12-13 [?] CRAN (R 4.2.0)
##  P Barycenter             1.3.1     2018-05-04 [?] CRAN (R 4.2.0)
##    Biobase              * 2.58.0    2022-11-01 [1] Bioconductor
##    BiocGenerics         * 0.44.0    2022-11-01 [1] Bioconductor
##  P BiocManager            1.30.19   2022-10-25 [?] CRAN (R 4.2.0)
##    BiocParallel           1.32.4    2022-12-01 [1] Bioconductor
##    BiocStyle            * 2.26.0    2022-11-01 [1] Bioconductor
##    biomformat             1.26.0    2022-11-01 [1] Bioconductor
##    Biostrings           * 2.66.0    2022-11-01 [1] Bioconductor
##    bit                    4.0.5     2022-11-15 [1] CRAN (R 4.2.0)
##    bit64                  4.0.5     2020-08-30 [1] CRAN (R 4.2.0)
##    bitops                 1.0-7     2021-04-24 [1] CRAN (R 4.2.0)
##    blob                   1.2.3     2022-04-10 [1] CRAN (R 4.2.0)
##    bookdown               0.31      2022-12-13 [1] CRAN (R 4.2.0)
##  P broom                  1.0.1     2022-08-29 [?] CRAN (R 4.2.0)
##  P cachem                 1.0.6     2021-08-19 [?] CRAN (R 4.2.0)
##  P callr                  3.7.3     2022-11-02 [?] CRAN (R 4.2.0)
##  P car                    3.1-1     2022-10-19 [?] CRAN (R 4.2.0)
##  P carData                3.0-5     2022-01-06 [?] CRAN (R 4.2.0)
##  P cli                    3.4.1     2022-09-23 [?] CRAN (R 4.2.0)
##  P cluster                2.1.4     2022-08-22 [3] CRAN (R 4.2.0)
##    coda                   0.19-4    2020-09-30 [1] CRAN (R 4.2.0)
##  P codetools              0.2-18    2020-11-04 [3] CRAN (R 4.2.1)
##    colorspace             2.0-3     2022-02-21 [1] CRAN (R 4.2.0)
##  P cowplot                1.1.1     2020-12-30 [?] CRAN (R 4.2.0)
##  P crayon                 1.5.2     2022-09-29 [?] CRAN (R 4.2.0)
##    data.table             1.14.6    2022-11-16 [1] CRAN (R 4.2.0)
##    DBI                    1.1.3     2022-06-18 [1] CRAN (R 4.2.0)
##    DECIPHER             * 2.26.0    2022-11-01 [1] Bioconductor
##    DelayedArray           0.24.0    2022-11-01 [1] Bioconductor
```

```
##    DESeq2              * 1.38.2    2022-12-14 [1] Bioconductor
##  P devtools            * 2.4.5     2022-10-11 [?] CRAN (R 4.2.0)
##    digest                0.6.31    2022-12-11 [1] CRAN (R 4.2.0)
##    DirichletReg        * 0.7-1     2021-05-18 [3] CRAN (R 4.2.0)
##  P dplyr               * 1.0.10    2022-09-01 [?] CRAN (R 4.2.0)
##  P ellipsis              0.3.2     2021-04-29 [?] CRAN (R 4.2.0)
##    evaluate              0.19      2022-12-13 [1] CRAN (R 4.2.0)
##  P fansi                 1.0.3     2022-03-24 [?] CRAN (R 4.2.0)
##    farver                2.1.1     2022-07-06 [1] CRAN (R 4.2.0)
##  P fastmap               1.1.0     2021-01-25 [?] CRAN (R 4.2.0)
##    forcats             * 0.5.2     2022-08-19 [1] CRAN (R 4.2.0)
##    foreach               1.5.2     2022-02-02 [1] CRAN (R 4.2.0)
##    Formula             * 1.2-4     2020-10-16 [3] CRAN (R 4.2.0)
##  P fs                    1.5.2     2021-12-08 [?] CRAN (R 4.2.0)
##    geneplotter           1.76.0    2022-11-01 [1] Bioconductor
##  P generics              0.1.3     2022-07-05 [?] CRAN (R 4.2.0)
##    GenomeInfoDb        * 1.34.4    2022-12-01 [1] Bioconductor
##    GenomeInfoDbData      1.2.9     2022-12-19 [1] Bioconductor
##    GenomicRanges       * 1.50.2    2022-12-16 [1] Bioconductor
##    geomnet             * 0.3.1     2022-12-19 [1] Github (sctyner/geomnet@030537d)
##    ggnewscale          * 0.4.8     2022-10-06 [1] CRAN (R 4.2.0)
##    ggplot2             * 3.4.0     2022-11-04 [1] CRAN (R 4.2.0)
##  P ggpubr              * 0.5.0     2022-11-16 [?] CRAN (R 4.2.1)
##  P ggrepel             * 0.9.2     2022-11-06 [?] CRAN (R 4.2.0)
##  P ggsignif              0.6.4     2022-10-13 [?] CRAN (R 4.2.0)
##  P glue                  1.6.2     2022-02-24 [?] CRAN (R 4.2.0)
##  P gridExtra             2.3       2017-09-09 [?] CRAN (R 4.2.0)
##    gtable                0.3.1     2022-09-01 [1] CRAN (R 4.2.0)
##    HiddenSemiMarkov    * 0.1.0     2022-11-23 [3] Github (lasy/HiddenSemiMarkov@2a5e9b4)
##  P highr                 0.9       2021-04-16 [?] CRAN (R 4.2.0)
##    hms                   1.1.2     2022-08-19 [1] CRAN (R 4.2.0)
##    htmltools             0.5.4     2022-12-07 [1] CRAN (R 4.2.0)
##    htmlwidgets           1.6.0     2022-12-15 [1] CRAN (R 4.2.0)
##  P httpuv                1.6.6     2022-09-08 [?] CRAN (R 4.2.0)
##  P httr                  1.4.4     2022-08-17 [?] CRAN (R 4.2.0)
##    igraph                1.3.5     2022-09-22 [1] CRAN (R 4.2.0)
##    IRanges             * 2.32.0    2022-11-01 [1] Bioconductor
##    iterators             1.0.14    2022-02-05 [1] CRAN (R 4.2.0)
##    jsonlite              1.8.4     2022-12-06 [1] CRAN (R 4.2.0)
##    KEGGREST              1.38.0    2022-11-01 [1] Bioconductor
##    knitr               * 1.41      2022-11-18 [1] CRAN (R 4.2.0)
##    labeling              0.4.2     2020-10-20 [1] CRAN (R 4.2.0)
##  P later                 1.3.0     2021-08-18 [?] CRAN (R 4.2.0)
##  P lattice               0.20-45   2021-09-22 [3] CRAN (R 4.2.1)
##    lazyeval              0.2.2     2019-03-15 [1] CRAN (R 4.2.0)
##  P lifecycle             1.0.3     2022-10-07 [?] CRAN (R 4.2.0)
##    limma                 3.54.0    2022-11-01 [1] Bioconductor
##    locfit                1.5-9.6   2022-07-11 [1] CRAN (R 4.2.0)
##    magick              * 2.7.3     2021-08-18 [1] CRAN (R 4.2.0)
##  P magrittr            * 2.0.3     2022-03-30 [?] CRAN (R 4.2.0)
##  P MASS                  7.3-58.1  2022-08-03 [3] CRAN (R 4.2.0)
##  P Matrix                1.5-3     2022-11-11 [3] CRAN (R 4.2.0)
##    MatrixGenerics      * 1.10.0    2022-11-01 [1] Bioconductor
##    matrixStats         * 0.63.0    2022-11-18 [1] CRAN (R 4.2.0)
##    maxLik                1.5-2     2021-07-26 [3] CRAN (R 4.2.0)
##  P memoise               2.0.1     2021-11-26 [?] CRAN (R 4.2.0)
##  P mgcv                  1.8-41    2022-10-21 [3] CRAN (R 4.2.0)
##  P mime                  0.12      2021-09-28 [?] CRAN (R 4.2.0)
##  P miniUI                0.1.1.1   2018-05-18 [?] CRAN (R 4.2.0)
```

```
##    miscTools              0.6-26    2019-12-08 [3] CRAN (R 4.2.0)
##    modeltools             0.2-23    2020-03-05 [1] CRAN (R 4.2.0)
##    MultiAssayExperiment * 1.24.0    2022-11-01 [1] Bioconductor
##    multtest               2.54.0    2022-11-01 [1] Bioconductor
##    munsell                0.5.0     2018-06-12 [1] CRAN (R 4.2.0)
##    network                1.18.0    2022-10-06 [1] CRAN (R 4.2.0)
##  P nlme                   3.1-161   2022-12-15 [3] CRAN (R 4.2.0)
##    NLP                    0.2-1     2020-10-14 [1] CRAN (R 4.2.0)
##    pbs                  * 1.1       2013-06-08 [1] CRAN (R 4.2.0)
##    permute                0.9-7     2022-01-27 [1] CRAN (R 4.2.0)
##    philentropy          * 0.7.0     2022-11-05 [1] CRAN (R 4.2.0)
##    phyloseq             * 1.42.0    2022-11-01 [1] Bioconductor
##  P pillar                 1.8.1     2022-08-19 [?] CRAN (R 4.2.0)
##  P pkgbuild               1.3.1     2021-12-20 [?] CRAN (R 4.2.0)
##  P pkgconfig              2.0.3     2019-09-22 [?] CRAN (R 4.2.0)
##  P pkgload                1.3.2     2022-11-16 [?] CRAN (R 4.2.1)
##    plotly                 4.10.1    2022-11-07 [1] CRAN (R 4.2.0)
##    plyr                   1.8.8     2022-11-11 [1] CRAN (R 4.2.0)
##    png                    0.1-8     2022-11-29 [1] CRAN (R 4.2.0)
##    preprocessCore         1.60.1    2022-12-14 [1] Bioconductor
##  P prettyunits            1.1.1     2020-01-24 [?] CRAN (R 4.2.0)
##  P processx               3.8.0     2022-10-26 [?] CRAN (R 4.2.0)
##  P profvis                0.3.7     2020-11-02 [?] CRAN (R 4.2.0)
##  P promises               1.2.0.1   2021-02-11 [?] CRAN (R 4.2.0)
##  P ps                     1.7.2     2022-10-26 [?] CRAN (R 4.2.0)
##  P purrr                * 0.3.5     2022-10-06 [?] CRAN (R 4.2.0)
##  P R6                     2.5.1     2021-08-19 [?] CRAN (R 4.2.0)
##  P ragg                   1.2.4     2022-10-24 [?] CRAN (R 4.2.0)
##    RColorBrewer         * 1.1-3     2022-04-03 [1] CRAN (R 4.2.0)
##  P Rcpp                   1.0.9     2022-07-08 [?] CRAN (R 4.2.0)
##    RCurl                  1.98-1.9  2022-10-03 [1] CRAN (R 4.2.0)
##    readr                * 2.1.3     2022-10-01 [1] CRAN (R 4.2.0)
##  P remotes                2.4.2     2021-11-30 [?] CRAN (R 4.2.0)
##    reshape2               1.4.4     2020-04-09 [1] CRAN (R 4.2.0)
##    rhdf5                  2.42.0    2022-11-01 [1] Bioconductor
##    rhdf5filters           1.10.0    2022-11-01 [1] Bioconductor
##    Rhdf5lib               1.20.0    2022-11-01 [1] Bioconductor
##  P rlang                  1.0.6     2022-09-24 [?] CRAN (R 4.2.0)
##    rmarkdown              2.19      2022-12-15 [1] CRAN (R 4.2.0)
##    RSQLite              * 2.2.19    2022-11-24 [1] CRAN (R 4.2.0)
##  P rstatix                0.7.1     2022-11-09 [?] CRAN (R 4.2.0)
##  P rstudioapi             0.14      2022-08-22 [?] CRAN (R 4.2.0)
##    S4Vectors            * 0.36.1    2022-12-05 [1] Bioconductor
##    sandwich               3.0-2     2022-06-15 [3] CRAN (R 4.2.0)
##    scales                 1.2.1     2022-08-20 [1] CRAN (R 4.2.0)
##  P sessioninfo            1.2.2     2021-12-06 [?] CRAN (R 4.2.0)
##  P shiny                  1.7.3     2022-10-25 [?] CRAN (R 4.2.0)
##    slam                 * 0.1-50    2022-01-08 [1] CRAN (R 4.2.0)
##    sna                    2.7       2022-06-01 [1] CRAN (R 4.2.0)
##    statnet.common         4.7.0     2022-09-08 [1] CRAN (R 4.2.0)
##  P stringi                1.7.8     2022-07-11 [?] CRAN (R 4.2.0)
##    stringr              * 1.5.0     2022-12-02 [1] CRAN (R 4.2.0)
##    SummarizedExperiment * 1.28.0    2022-11-01 [1] Bioconductor
##  P survival               3.4-0     2022-08-09 [3] CRAN (R 4.2.0)
##  P systemfonts            1.0.4     2022-02-11 [?] CRAN (R 4.2.0)
##  P textshaping            0.3.6     2021-10-13 [?] CRAN (R 4.2.0)
##  P tibble                 3.1.8     2022-07-22 [?] CRAN (R 4.2.0)
##    tidyr                * 1.2.1     2022-09-08 [1] CRAN (R 4.2.0)
##  P tidyselect             1.2.0     2022-10-10 [?] CRAN (R 4.2.0)
```

```
##    tm                    0.7-9     2022-10-19 [1] CRAN (R 4.2.1)
##    topicmodels         * 0.2-12    2021-01-29 [1] CRAN (R 4.2.0)
##    tzdb                  0.3.0     2022-03-28 [1] CRAN (R 4.2.0)
##  P urlchecker            1.0.1     2021-11-30 [?] CRAN (R 4.2.0)
##  P usethis             * 2.1.6     2022-05-25 [?] CRAN (R 4.2.0)
##  P utf8                  1.2.2     2021-07-24 [?] CRAN (R 4.2.0)
##  P vctrs                 0.5.1     2022-11-16 [?] CRAN (R 4.2.1)
##    vegan                 2.6-4     2022-10-11 [1] CRAN (R 4.2.0)
##    viridisLite           0.4.1     2022-08-22 [1] CRAN (R 4.2.0)
##    vsn                 * 3.66.0    2022-11-01 [1] Bioconductor
##  P withr                 2.5.0     2022-03-03 [?] CRAN (R 4.2.0)
##  P xfun                  0.35      2022-11-16 [?] CRAN (R 4.2.1)
##    XML                   3.99-0.13 2022-12-04 [1] CRAN (R 4.2.0)
##  P xml2                  1.3.3     2021-11-30 [?] CRAN (R 4.2.0)
##  P xtable                1.8-4     2019-04-21 [?] CRAN (R 4.2.0)
##    XVector             * 0.38.0    2022-11-01 [1] Bioconductor
##  P yaml                  2.3.6     2022-10-18 [?] CRAN (R 4.2.0)
##    zlibbioc              1.44.0    2022-11-01 [1] Bioconductor
##    zoo                   1.8-11    2022-09-17 [3] CRAN (R 4.2.0)
##
##   [1]  /Users/laurasymul/Documents/Work/Research/Packages/alto/renv/library/R-4.2/x86_64-apple-
darwin17.0
##  [2] /private/var/folders/b9/hxr_p5z56m51ckk_jl995qy00000gn/T/Rtmptswmnv/renv-system-library
##  [3] /Library/Frameworks/R.framework/Versions/4.2/Resources/library
##
##  P -- Loaded and on-disk path mismatch.
##
## ------------------------------------------------------------------------------
```
